# cryoAgent: An agentic workflow for robust and adaptive end-to-end cryo-EM image processing

**DOI:** 10.64898/2026.04.16.718662

**Authors:** Daoyi Li, Shaodi Yang, Taining Pei, Chen Qi, Qian Xiao, Tongxin Niu, Yan Zhang, Yun Zhu, Fei Sun

**Author notes:** Corresponding author: Fei Sun, **Email:**. **Author Contributions:** D.L., F.S., Y.Z., and Y.Z. conceived and led the study. D.L., F.S., and Y.Z. designed the experimental study. D.L. and S.Y. designed and implemented the algorithms and agentic architecture. D.L. and S.Y., T.P. performed experiments across the datasets. S.Y., C.Q., Q.X., T.N., and D.L. prepared the experimental datasets. S.Y. and D.L. refined the 3D models for the density maps and compared local density quality. F.S. and Y.Z. acquired funding, and F.S. supervised project administration. All authors contributed to writing and revising the manuscript. **Competing Interest Statement:** The authors declare no competing interests. **Classification:** Biological Sciences, Biophysics and Computational Biology.

## Abstract

Cryo-electron microscopy (cryo-EM) has become the mainstream method for high-resolution structure determination, yet current automated workflows still require substantial expert intervention for failure recovery, heterogeneity analysis, and parameter optimization. Here, we present cryoAgent, an agentic workflow for autonomous cryo-EM image processing with adaptive tool use to address these challenges. cryoAgent progressively expands its autonomy from a rigid workflow that generates a reliable baseline reconstruction, through guided processing for iterative optimization and heterogeneity analysis, to hypothesis-driven exploration of alternative processing strategies. Across diverse cryo-EM datasets, cryoAgent improves reconstruction quality and outperforms the state-of-the-art automated workflow. The autonomous heterogeneity analysis capability also enables the identification of a previously unreported structural state. cryoAgent therefore provides a scalable framework for autonomous cryo-EM structure determination.

## Introduction

Cryo–electron microscopy (cryo-EM) has become a transformative technique in structural biology, enabling high-resolution structure determination of diverse macromolecules (1). Advances in automated data acquisition now allow modern microscopes to collect tens of thousands of micrographs in a single data collection session (2). As cryo-EM is applied to an increasingly diverse range of biological targets, there is a growing demand for efficient, reliable, and scalable automated image-processing workflows capable of handling large datasets without extensive manual intervention.

Current automated cryo-EM pipelines have substantially reduced manual effort in single-particle analysis, particularly for well-behaved, homogeneous or previously characterized datasets (3–9). However, several key stages still rely heavily on expert judgment. First, existing pipelines lack robust mechanisms for failure recovery. If a step fails, the workflow typically halts or silently degrades without adapting parameters. Second, heterogeneity analysis remains largely manual. It requires users to interpret 3D classification results and decide which classes represent meaningful conformations to target. Third, most automated pipelines do not support iterative optimization. In contrast to expert workflows—which often involve multiple rounds of 2D and 3D classification with progressively refined parameters—automated pipelines usually execute a fixed sequence and terminate after an initial reconstruction, limiting final map quality. Therefore, automated workflows may terminate prematurely or yield reconstructions that fall short of expert-guided outcomes, limiting their utility for final high-quality structure determination.

Addressing these limitations requires moving beyond rigid, predefined workflows toward systems capable of dynamic reasoning and adaptive control. Agentic pipelines powered by large language models (LLMs) introduce decision-making capabilities at key stages of data processing, enabling workflows to evaluate intermediate results, revise strategies, and selectively deploy tools based on contextual cues (10, 11). However, applying open-ended exploration from the beginning is neither reliable nor efficient for cryo-EM image processing, where many steps are already well established and each exploratory branch can require substantial computational resources (12). A guided workflow provides a structured execution framework that encodes established domain knowledge, tool-use procedures, and operational constraints. Such a harness guides the agent along reliable processing pathways while preserving sufficient flexibility to adjust parameters, repeat operations, determine stopping points, and pursue broader hypothesis-driven exploration after a stable result has been obtained (13).

Here, we present cryoAgent, a multi-agent framework that integrates ReAct-based tool use into the cryo-EM single-particle analysis pipeline (14). cryoAgent progressively expands its autonomy across three levels. First, a rigid workflow executes the essential stages of preprocessing, particle picking, and reconstruction to generate a reproducible baseline while monitoring jobs and recovering failed operations. Second, guided processing invokes specialized optimization and heterogeneity agents that adjust parameters, repeat operations, and determine stopping points based on intermediate results. Third, hypothesis-driven exploration uses the accumulated outputs and processing history to formulate and test strategies beyond the encoded workflows. This design ensures reliable execution during initialization while progressively enabling open-ended reasoning as dataset-specific evidence becomes available (Fig. 1).

**Figure 1.**
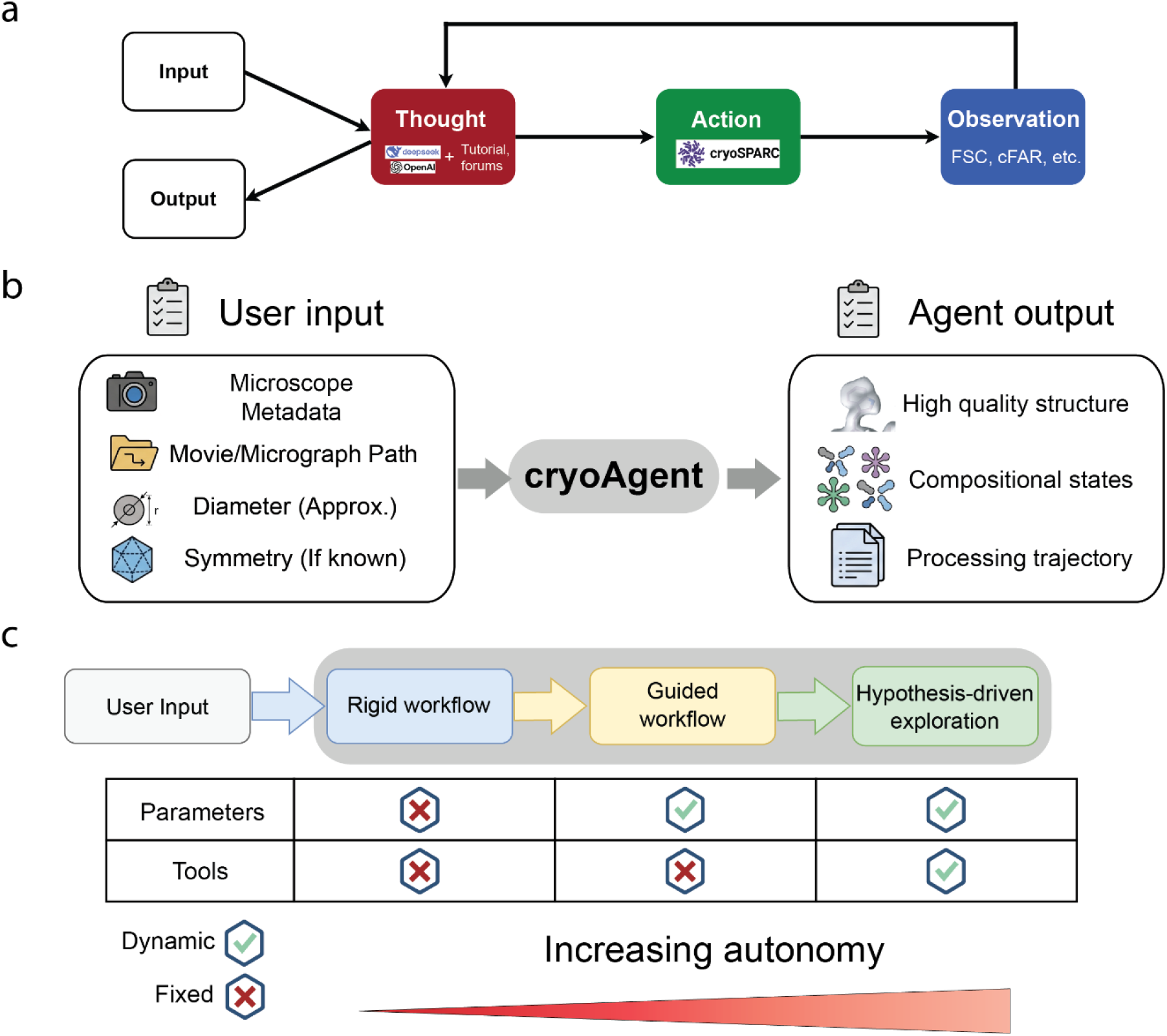
cryoAgent architecture and progressive levels of autonomy. A. cryoAgent operates through a ReAct loop, in which an LLM-based Thought module reasons about the current processing state and determines the next step, the Action module executes external tools, and the Observe module evaluates the resulting outputs to determine whether further iteration is required. B. Input and output of cryoAgent. C. cryoAgent is organized into three processing stages with increasing levels of autonomy. The input data are first processed by a rigid workflow to generate an initial reconstruction. The guided workflow then follows a fixed tool-use sequence while allowing parameter adjustment. Finally, the hypothesis-driven exploration stage formulates and tests additional processing strategies to further improve reconstruction quality.

Across diverse datasets, this progressive architecture improved reconstruction quality relative to the corresponding deposited density maps and achieved state-of-the-art performance compared with existing end-to-end automated method. In addition, the heterogeneity and exploration agents enabled the autonomous identification of a previously overlooked structural state. Together, these results demonstrate that cryoAgent extends cryo-EM automation beyond reliable workflow execution toward adaptive image processing and autonomous structural discovery.

## Results

### cryoAgent architecture and progressive levels of autonomy

cryoAgent integrates LLM-based reasoning with executable cryo-EM processing tools, enabling processing decisions generated within the ReAct loop to be translated directly into computational actions (Fig. 1a). To initiate analysis, users provide only essential experimental information, including microscope and data-acquisition metadata, the micrograph location, the approximate particle diameter, and the expected symmetry (Fig. 1b). A central orchestrator maintains the processing state, coordinates specialized agents, and transfers intermediate results between processing stages. This design separates workflow-level coordination from task-specific analysis while providing a unified interface for tool execution, result interpretation, and decision-making.

To balance processing reliability with adaptive reasoning, cryoAgent progressively expands its autonomy across three processing levels (Fig. 1c). At the first level, it executes a rigid workflow with predetermined processing steps and parameters to efficiently generate baseline reconstruction while monitoring tool execution and recovering failed operations. At the second level, specialized agents operate within stage-specific workflow harnesses to address complementary limitations in compositional heterogeneity analysis and reconstruction quality, with intermediate results guiding parameter adjustment, iterative processing, and stopping decisions. At the third level, cryoAgent uses the accumulated outputs and processing trajectories to generate hypotheses, formulate new processing plans, and adaptively execute available tools beyond the predefined harnesses. The architecture therefore progresses from reliable baseline execution, through structured agentic analysis, to hypothesis-driven exploration.

### Failure recovery supports uninterrupted tool execution

Reliable execution is essential for LLM-based agents that interact with scientific processing tools. In cryo-EM workflows, achieving such reliability requires detecting and recovering from two distinct forms of failure. The first is an explicit execution failure, in which a job terminates because of invalid parameters, incompatible inputs, software errors, or resource limitations. The second is a processing-state failure, in which a job completes successfully but produces an output that is unsuitable for downstream analysis (Fig. 2a). An agentic system must therefore do more than issue tool calls: it must monitor execution, diagnose both explicit and implicit failures, and adapt its actions to restore a viable processing trajectory.

**Figure 2.**
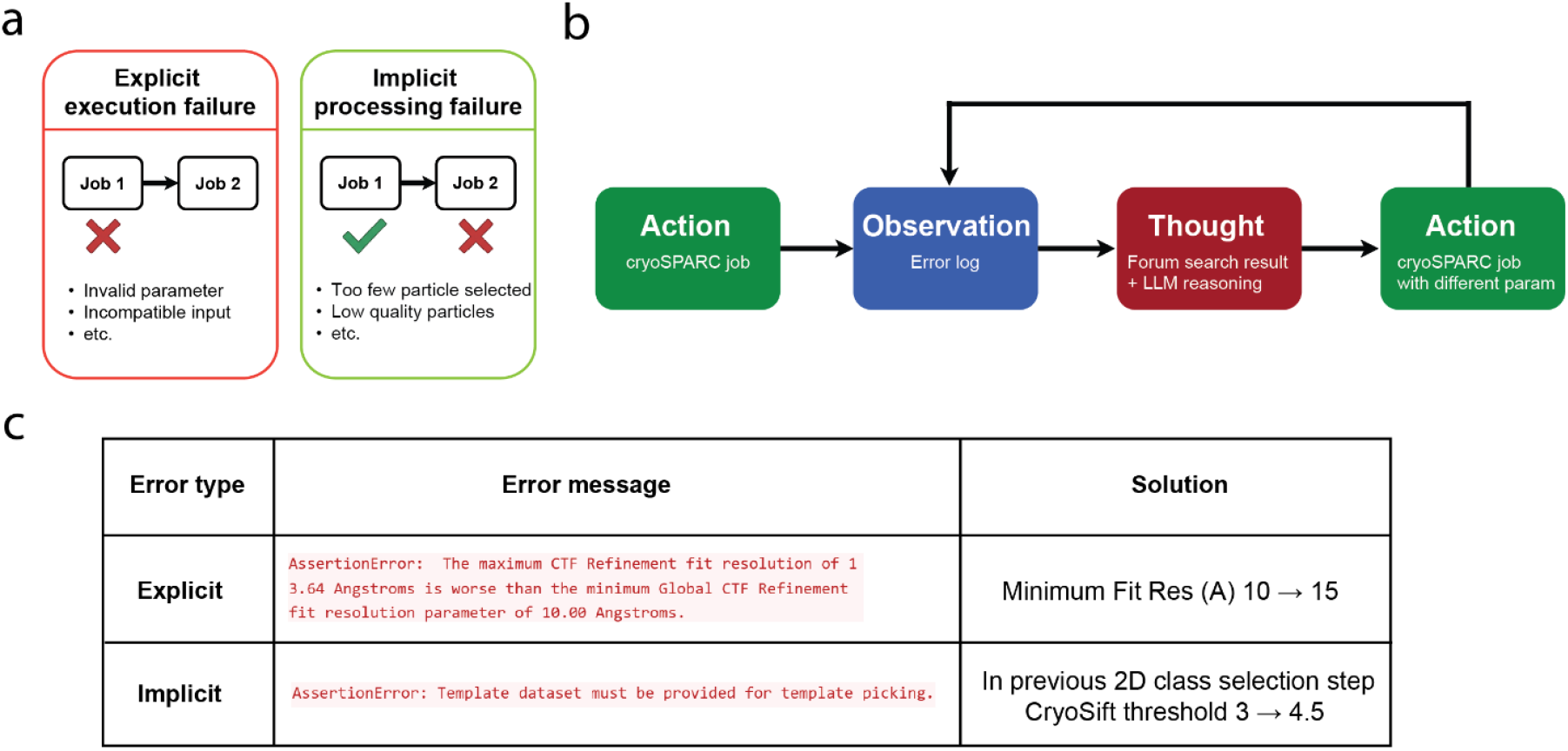
Failure recovery mechanism through an agentic loop. A. Two representative types of processing failure. B. Agentic reasoning process triggered by an error log. C. Representative examples of each error type and the corresponding recovery strategies.

When an execution failure occurs, cryoAgent retrieves the associated error log and uses the reported message to search the cryoSPARC discussion forum for relevant troubleshooting information (15). It examines the top search results, summarizes the proposed causes and solutions, and maps these recommendations onto the parameters exposed by the corresponding cryoAgent tool. The agent then identifies which available parameters are most likely to resolve the failure, modifies the tool call, and re-executes the job. For processing-state failures, cryoAgent detects cases in which a completed job yields insufficient particles for subsequent processing, identifies the limiting selection parameters, relaxes the corresponding thresholds, and repeats the operation. This mechanism enables cryoAgent to recover from both explicit job failures and completed-but-unusable processing outcomes that would otherwise interrupt end-to-end reconstruction (Fig. 2b). Representative examples demonstrate that this recovery mechanism enables cryoAgent to successfully resolve both execution failures (Fig. 2c).

### Structured workflow harnesses improve the reliability and quality of agentic cryo-EM processing

With automated failure recovery in place, cryoAgent could execute an unconstrained agentic process with only minimal guide from the initial input and continue processing despite occasional tool-execution errors. In this setting, the agent first consulted the cryoSPARC tutorials and documentation, generated an end-to-end processing plan, and selected and executed the corresponding tools (Fig. 3a). We evaluated this *de novo* exploration (DE) using the RELION tutorial datasets (16), which contains only 24 movies and therefore provides a compact, well-controlled benchmark for testing agentic workflow execution (Table S1). Because neither processed data nor workflow-guiding prompts were available at initialization, DE sometimes failed to identify and execute an effective end-to-end processing strategy. This instability was more pronounced with lower-capability language models (Table S2). We therefore encoded a rigid workflow with predefined parameters and tool usage, while reserving broader reasoning and exploration for later stages after informative intermediate results had been generated.

**Figure 3.**
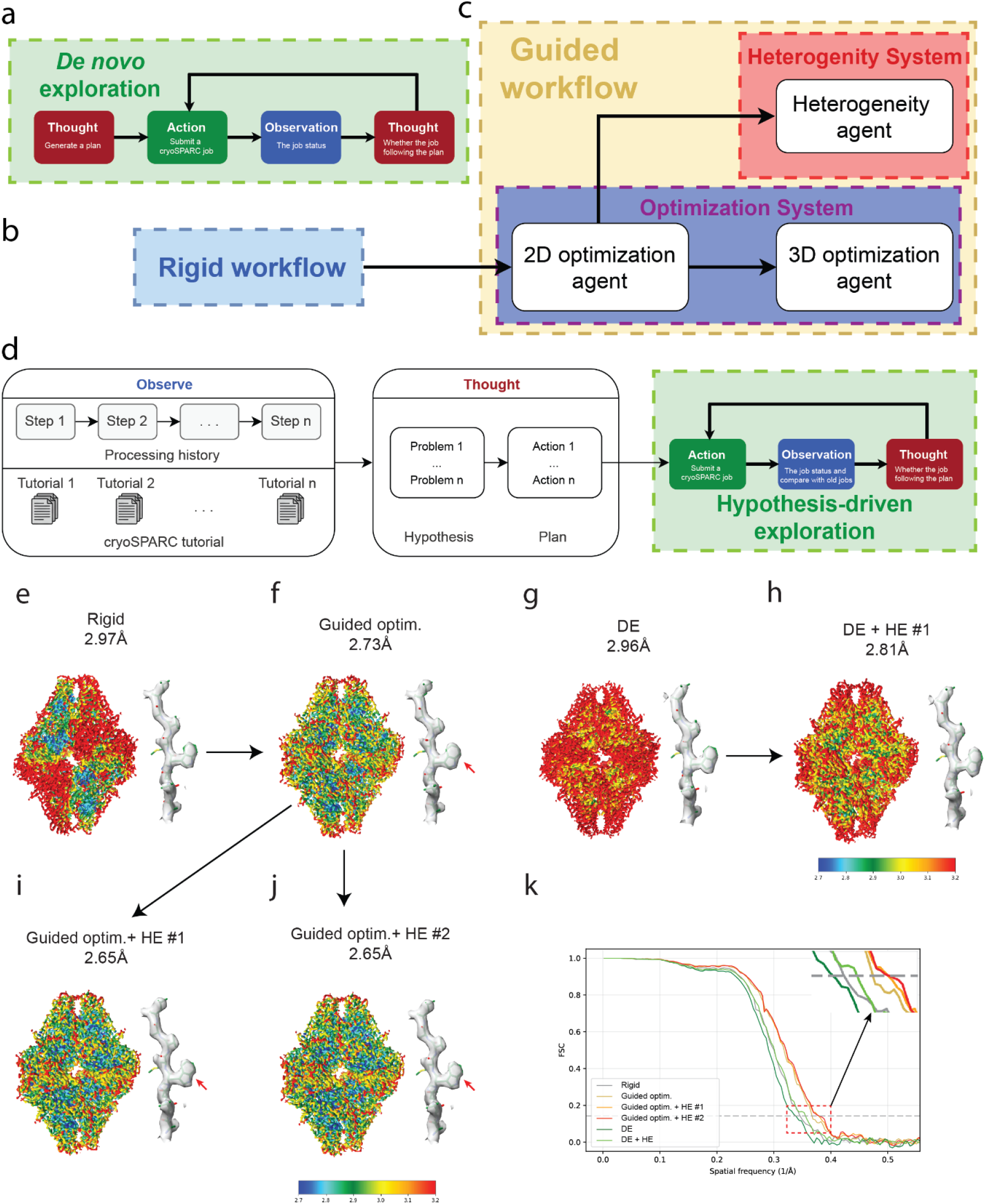
Three levels of autonomy in cryoAgent. A. De novo exploration mode, in which the agent generates a processing plan based on CryoSPARC tutorials and dynamically selects tools and parameters to achieve end-to-end reconstruction. B. Rigid workflow, in which reconstruction is performed using a fixed sequence of tools and predefined parameters. C. Guided workflow, in which the agent follows a predefined tool-use logic after the rigid workflow while adjusting parameters to maximize reconstruction quality and resolve compositional heterogeneity. D. Hypothesis-driven exploration mode. Based on the reconstruction trajectory from either dynamic exploration mode or guided workflow, together with cryoSPARC tutorial knowledge, the agent generates a set of hypotheses and corresponding processing plans. Each hypothesis is then tested to determine whether it improves the final reconstruction. E–J. Reconstruction of the β-galactosidase RELION tutorial dataset using the rigid workflow (E), guided optimization initialized from the rigid-workflow result (F), de novo exploration mode (G), and hypothesis-driven exploration initialized from either the de novo exploration result (H) or the guided-optimization result (I–J). K. FSC curves for the RELION tutorial dataset obtained using different agentic processing strategies. Gray, rigid workflow; brown, rigid workflow followed by guided optimization; yellow, rigid workflow followed by guided optimization and hypothesis-driven exploration in trial 1; red, rigid workflow followed by guided optimization and hypothesis-driven exploration in trial 2; green, de novo exploration; light green, de novo exploration followed by hypothesis-driven exploration.

The rigid workflow is designed to produce a stable initial reconstruction with minimal reasoning and tool-execution overhead (Fig. 3b). Starting from a predefined processing sequence reduces the risk of inefficient or unstable exploration before sufficient information about the dataset has been obtained, while providing a consistent baseline for subsequent optimization and heterogeneity analysis. Within this workflow, the processing stages, tool sequence, and associated parameters are predetermined. The LLM is therefore responsible primarily for executing and monitoring the prescribed tool calls rather than exploring the parameter space or generating alternative processing strategies. This constrained design enables cryoAgent to efficiently obtain an initial reconstruction under standardized processing conditions, with the complete detail shown in Fig. S1.

Starting from the initial reconstruction generated by the rigid workflow, cryoAgent routes the dataset to two specialized agent systems that address complementary limitations of automated cryo-EM processing: compositional heterogeneity and reconstruction quality. Both systems operate within stage-specific workflow harnesses that encode established processing procedures, evaluation criteria, while allowing the agents to explore the relevant parameter space and adapt their decisions to intermediate results.

The heterogeneity agent is designed to identify distinct compositional states and determine how many should be retained for downstream analysis (Fig. 3c). Because LLMs cannot directly interpret three-dimensional density maps, cryoAgent represents the results of each classification round through a graph-based quantitative framework (Fig. S2a). The reconstructed volumes are aligned using cryoAlign (17) and FitMap (18), and pairwise similarity measurements are used to construct a graph in which community detection estimates the number of distinct compositional states (Fig. S3). It iteratively performs 3D classifications using multiple values of K and continues until additional rounds no longer reveal new compositional states. These results are returned to the agent’s ReAct loop to determine the class number and parameters for the next round or to terminate the analysis when no additional compositional state is detected (Fig. S4).

In parallel, the optimization agent improves reconstruction quality through dedicated 2D and 3D agents operating in iterative, dataset-specific workflows (Fig. 3c, S2b). The 2D agent performs repeated rounds of 2D classification and evaluates class quality using cryoSift-derived metrics (Fig. S5), whereas the 3D agent iteratively performs 3D classification and box-size optimization and evaluates the resulting reconstructions using using FSC (Fourier Shell Correlation) and cFAR (Conical FSC Area Ratio) (Fig. S6). These quantitative evaluations are fed back to the respective agents to guide parameter adjustment, further processing, and stopping decisions.

The preceding results established that cryoAgent can reliably execute both rigid and guided processing strategies and that exploration might be more effective when applied after a controlled processing stage with rich processing history (Fig. 3d). In particular, the preliminary reconstruction generated by guided optimization outperformed those produced by both the rigid workflow and DE (Fig. 3e–g, k), providing a stronger starting point for subsequent exploration. We next examined whether it could move beyond these encoded workflows to autonomously formulate and test new strategies for further reconstruction improvement.

### Hypothesis-driven exploration autonomously formulates and executes dataset-specific processing strategies

In the exploration mode, cryoAgent jointly interprets the available tool outputs and processing history to identify factors that may continue to limit reconstruction quality. It then formulates a hypothesis about the underlying limitations and consults the cryoSPARC Guide to identify relevant processing strategies and candidate solutions. The agent translates this hypothesis into an executable plan specifying the sequence of processing steps, tools, and parameters to be evaluated. It subsequently executes the plan, assesses the resulting outputs, and determines whether the strategy improves the reconstruction or whether the hypothesis and plan should be revised. Through this iterative hypothesis–planning–execution–evaluation loop, cryoAgent can construct dataset-specific processing strategies that are not explicitly encoded in the DE or guided workflow (Fig. 3d).

We first asked whether hypothesis-driven exploration (HE) could rescue the suboptimal reconstruction generated from DE and produce a result comparable to that obtained through the rigid and guided workflow. After reviewing the processing trajectory, intermediate reconstructions, and quality measurements, cryoAgent identified three potential limitations: the final reconstruction had been generated using homogeneous refinement rather than non-uniform refinement; no separate CTF refinement had been performed; and the number of retained particles might have been insufficient. The agent therefore proposed three corresponding strategies: non-uniform refinement, additional CTF refinement, and particle repicking using the final density as a reference. Non-uniform refinement improved the reported resolution from 2.96 Å to 2.83 Å. A subsequent standalone CTF-refinement step produced no further improvement, potentially because CTF optimization had already been enabled during non-uniform refinement. Repicking and reconstructing an expanded particle set likewise failed to improve the map quality. Because non-uniform refinement produced a measurable gain, cryoAgent subsequently hypothesized that residual heterogeneity might still limit the reconstruction. The agent noted that no heterogeneity analysis had been performed in the DE and therefore executed heterogeneous refinement, selected the highest-quality particle group, and reconstructed it independently. This additional processing improved the final resolution to 2.81 Å (Fig. S7). Thus, HE successfully identified and tested several plausible limitations and partially rescued the *de novo* exploration result, although the final reconstruction remained inferior to that obtained through the rigid and then guided workflow (Fig. 3e-3f, 3k).

We next applied HE to the reconstruction obtained from the rigid workflow and guided optimization for the same dataset. In this setting, cryoAgent received richer context, including the outputs of individual tools and the stage-specific processing trajectories generated during the preceding analysis. To assess the stability of the exploration process, we performed two independent trials initialized from the same 2.73 Å reconstruction.

In the first trial, cryoAgent identified an opportunities for further improvement: additional CTF refinement. It first performed a separate CTF-refinement step with beam-tilt and trefoil parameters enabled, which marginally improved the resolution from 2.73 Å to 2.71 Å. After evaluating the updated trajectory, the agent recognized that reference-based motion correction had previously produced a substantial improvement and therefore performed an additional round, further improving the resolution to 2.68 Å. It then noted that higher-order CTF terms had not yet been comprehensively optimized and executed another CTF-refinement step with spherical aberration, tetrafoil, and anisotropic magnification enabled. This final step improved the reconstruction to 2.65 Å (Fig. S8).

In the second independent trial, cryoAgent initially hypothesized that the particle-selection threshold had been overly restrictive and relaxed the final selection criteria. Although this retained more particles, the resulting reconstruction was worse, leading the agent to reject this strategy. It next identified incomplete higher-order CTF optimization as a potential limitation and performed CTF refinement using the same higher-order parameters as in the first trial, improving the reconstruction to 2.66 Å. The agent subsequently performed ab initio reconstruction with K=3, used the class containing the largest particle population to initialize heterogeneous refinement, regrouped the resulting classes into two particle subsets, and selected the larger subset for non-uniform refinement. This trajectory further slightly improved the resolution to 2.65 Å. Finally, the agent generated templates from the resulting density, repeated particle picking and curation, and reconstructed the expanded particle set; however, this strategy produced no additional improvement (Fig. S8).

Despite following different reasoning and tool-execution trajectories, both independent trials converged on reconstructions of approximately 2.65 Å. It results in noticeable changes in the improvement of the local resolution and density quality of tryptophan (Fig. 3f, 3i-j). This result demonstrates that hypothesis-driven exploration can test alternative strategies, reject unproductive branches, and reproducibly improve an already optimized reconstruction.

### Autonomous heterogeneity analysis with cryoAgent reveals previously overlooked structural states

We evaluated whether the cryoAgent heterogeneity agent could autonomously determine the number of non-redundant structural states present in cryo-EM datasets. Across datasets, the heterogeneity profiles recovered by cryoAgent were consistent with those reported in the original studies (Fig. S9-S10). This agreement indicates that recursive classification did not systematically produce unsupported classes or over-segment the datasets.

Application of this workflow to EMPIAR-10028 (19) revealed a compositional state that had not been reported in the original analysis. Whereas the deposited reconstruction represented an 80S ribosome, recursive agent-guided classification separated the particles into two reproducible structural types (Fig.4a-d). Type 1 closely matched the deposited 80S ribosome, whereas Type 2 corresponded to an isolated 60S ribosomal subunit. The principal difference was the absence of the 40S subunit in Type 2, accompanied by difference of density around the inter subunit bridge region that connects the 60S and 40S subunits in the assembled 80S ribosome (Fig. 4e-f). These features support the interpretation of Type 2 as a compositionally distinct 60S population rather than an incompletely resolved 80S reconstruction.

**Figure 4.**
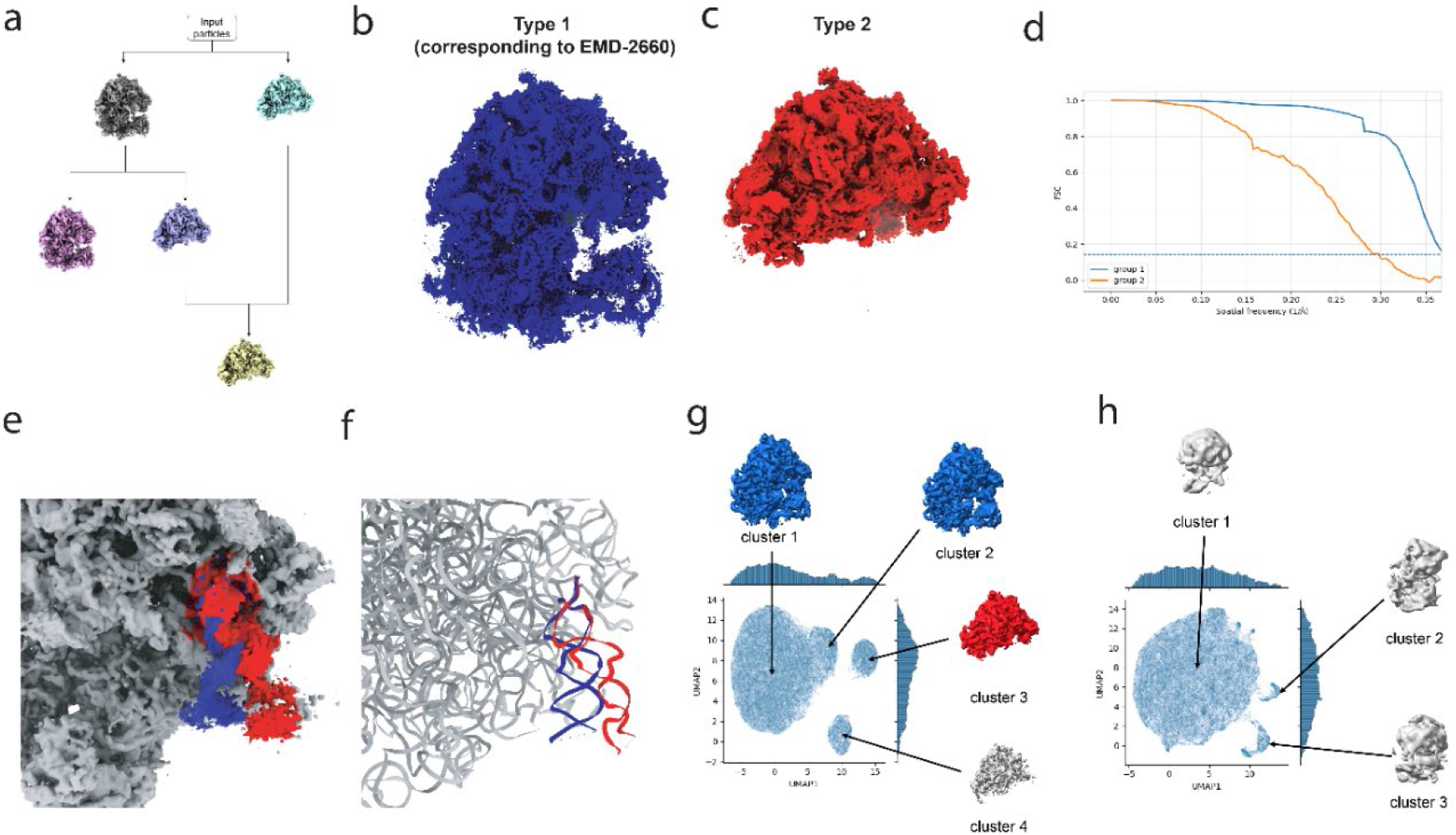
Autonomous heterogeneity analysis reveals a previously overlooked compositional state. A. Discovery workflow of the cryoAgent heterogeneity system under the guided workflow. B–C. In EMPIAR-10028, the heterogeneity agent identified two types of density. B. Type 1 corresponds to the deposited 80S ribosome map, EMDB-2660. C. Type 2 represents a newly identified 60S ribosome population. D. FSC curves for the different compositional states. E–F. Density and atomic-model comparisons reveal local conformational differences between the identified states.

We independently evaluated the identified heterogeneity using DRGN-AI (20). In fixed-pose DRGN-AI analysis, the latent distribution contained four clusters. Two clusters corresponded to major states identified in the previous cryoDRGN analysis (21), whereas another produced a density consistent with the isolated 60S state recovered by cryoAgent (Fig. 4c,4g). By contrast, pose-free DRGN-AI analysis did not produce an interpretable cluster separation (Fig. 4h). The agreement between cryoAgent and independent fixed-pose latent-variable analysis provides additional support for the previously overlooked 60S population.

### Cross-dataset benchmarking demonstrates generalizable end-to-end reconstruction ability

In EMPIAR-10181 (22), iterative agent-driven optimization progressively improved resolution, outperforming both the originally deposited reconstruction and cryoWizard (8), a state-of-the-art automated end-to-end cryo-EM image processing workflow (Fig. 5a-c, S11). With the hypothesis-driven exploration mode, the resolution improved slightly but not very much (Fig. 5c, S12). In more challenging cases, such as EMPIAR-10217, the optimization agent produced higher overall density quality relative to the rigid automated workflow (Fig. 2d-e).

**Figure 5.**
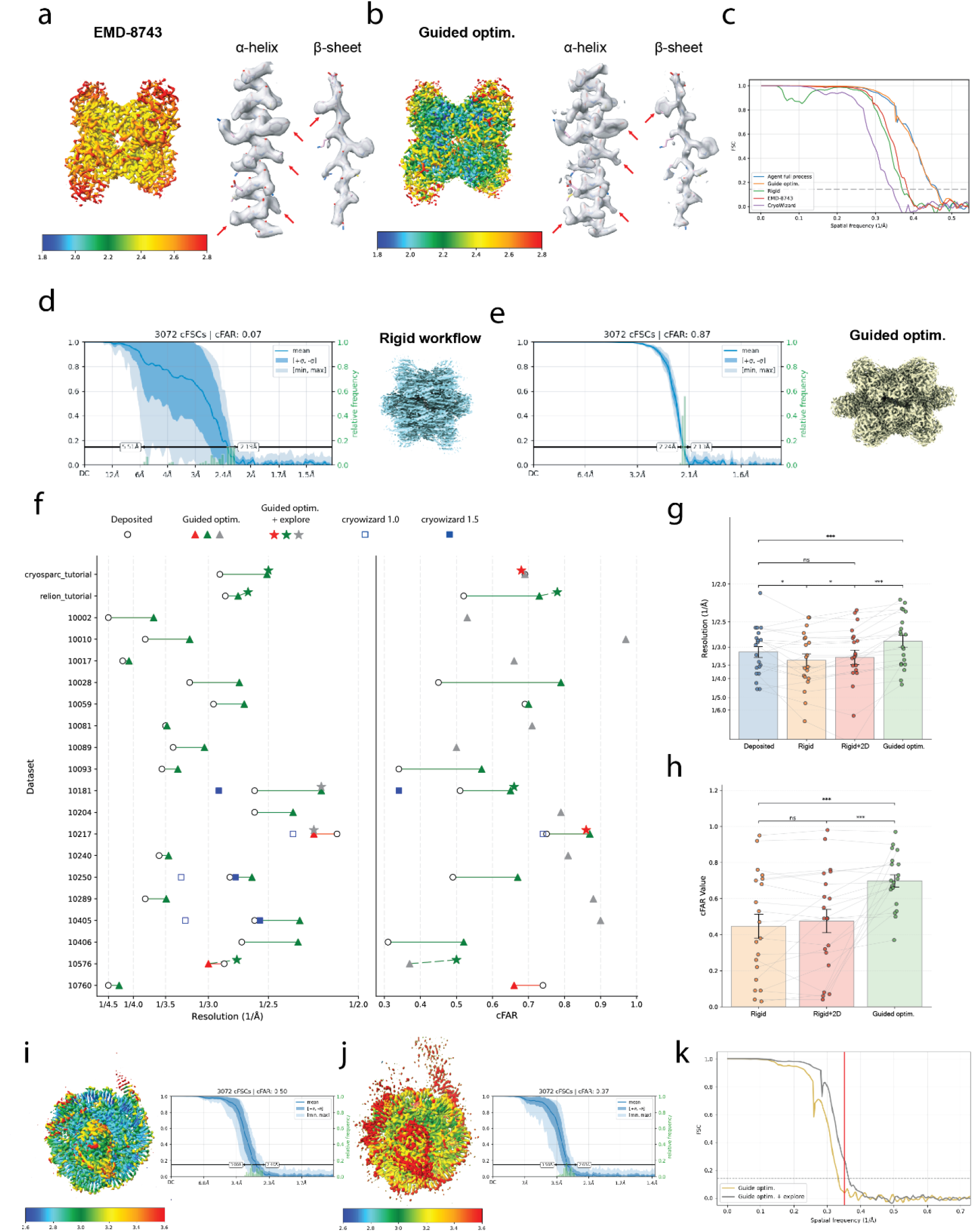
Cross-dataset benchmarking demonstrates generalizable reconstruction performance. a–b: Comparison between a. the deposited EMDB density map, EMDB-8743, and b. the cryoAgent-processed EMPIAR-10181 reconstruction. Density maps are colored by local resolution estimated from half maps. Representative regions containing α-helices and β-sheets are shown; red markers indicate residues with substantially improved local density quality. c: FSC curves for EMPIAR-10181 under different levels of agent-driven optimization. Green, rigid workflow; orange, rigid workflow + guided optimization; red, deposited EMDB FSC curve; blue, rigid workflow + guided optimization + HE; purple, cryoWizard. d–e: Density maps and cFAR scores for EMPIAR-10217 obtained using d. the rigid workflow and e. the full cryoAgent workflow. f–h: Across datasets, cryoAgent generally improves both resolution and cFAR scores. f. Dataset-level comparison of deposited maps, cryoAgent reconstructions, and cryoWizard results. Hollow circles indicate deposited maps; filled circles indicate cryoAgent reconstructions; blue squares indicate cryoWizard results. g–h. Box plots summarizing performance gains in g. resolution and h. cFAR across processing stages, with statistical significance assessed using the Wilcoxon signed-rank test. i-k: Comparison between the reconstruction and cFAR of the EMPIAR-10576 with guided optimization system and further hypothesis-driven reconstruction mode. Density maps are colored by local resolution estimated from half maps.

We evaluated cryoAgent on 20 cryo-EM datasets representing a broad range of single-particle reconstruction scenarios (Table S1). The targets ranged in molecular mass from 64 kDa to 6.7 MDa and contained approximately 10,000 to 1.8 million particles. The benchmark included complexes with different symmetries, particle abundances, initial reconstruction qualities, and degrees of structural heterogeneity. For each dataset, cryoAgent began from the minimal experimental inputs and first generated baseline reconstruction through the rigid workflow, followed by guided optimization and, where applicable, hypothesis-driven exploration.

Guided optimization improved reconstruction quality compared to the deposited densities in 18 of the 20 benchmark datasets, together with some reconstruction result from cryoWizard (Fig. S11, S13). Across the complete benchmark, paired comparisons between the original density map and guided optimization reconstructions showed significant improvements in both reciprocal resolution and cFAR (Fig. 5f-h, S14–S33). These results indicate that guided optimization consistently improved the baseline reconstructions across the benchmark rather than benefiting only a small subset of datasets. The two remaining datasets, EMPIAR-10217 and EMPIAR-10576, along with additional representative datasets, were further subjected to HE analysis (Figs.S10, S12, S34– S35). HE also led to a substantial improvement in the reconstruction quality of EMPIAR-10576 (Fig. 5i-k).

We next compared cryoAgent with existing automated and expert-guided processing approaches. Among the datasets previously processed by cryoWizard and available through EMDB-China, cryoAgent produced higher-resolution reconstructions and improved reconstruction-quality metrics for all evaluated datasets (Fig. 5f and Fig. S13). We additionally benchmarked cryoAgent on a dataset that had been processed independently by expert. Using only the rigid workflow and guided optimization, cryoAgent achieved reconstruction quality comparable to expert-curated results (Fig. S36). These comparisons demonstrate that cryoAgent can deliver competitive reconstruction performance on previously unseen datasets without access to target-specific prior knowledge or deposited density maps.

The improvements were accompanied by dataset-specific changes in the processing trajectory. The final particle populations selected by cryoAgent were not simply retained subsets of the particles produced by the rigid workflow; depending on the dataset, optimization removed low-quality particles, recovered particles excluded by earlier thresholds, or generated alternative subsets through additional classification and selection (Fig. S37). The final extraction box sizes also frequently differed from both the rigid-workflow settings and the corresponding deposited values (Fig. S38). Together with the variation in classification, refinement, and particle-selection trajectories across datasets, these results indicate that cryoAgent adapted its processing decisions to the intermediate evidence rather than applying a single predetermined optimization recipe.

Overall, these benchmark results demonstrate that cryoAgent can generalize across diverse cryo-EM datasets and consistently improve reconstructions from minimal experimental inputs. Its progressive architecture combines the reliability of rigid workflow execution with the adaptability of harness-guided optimization and hypothesis-driven exploration, enabling dataset-specific processing decisions without access to deposited maps or target-specific prior knowledge. By achieving reconstruction quality competitive with existing automated workflows and expert-guided analyses, cryoAgent provides a scalable framework for reliable end-to-end cryo-EM processing across a broad range of reconstruction scenarios.

## Discussion

In this study, we developed cryoAgent as a multi-agent framework that integrates large language model–based reasoning with executable cryo-EM processing tools. Its principal contribution is a progressive architecture that expands autonomy from rigid workflow execution, through guided workflow, to hypothesis-driven exploration. This design enabled reliable end-to-end processing, automated failure recovery, dataset-specific reconstruction optimization, and autonomous analysis of compositional heterogeneity. Across diverse cryo-EM datasets, cryoAgent improved reconstruction quality relative to its rigid-workflow baseline and achieved performance competitive with existing automated and expert-guided analyses. It also recovered a previously overlooked 60S ribosomal population in EMPIAR-10028, illustrating how agentic processing can extend beyond workflow completion toward discovery-oriented structural analysis.

A central implication of these results is that unrestricted agentic control is not equally useful throughout a cryo-EM image processing workflow. Since the agent has access to only limited experimental metadata from scratch, providing little evidence to propose meaningful dataset-specific decisions plan. *De novo* exploration frequently introduced redundant or unstable tool use. Encoding a rigid workflow improved reproducibility and efficiency without removing agent-controlled execution or failure recovery. Once intermediate reconstructions and quantitative evaluations became available, dataset-specific reasoning became more informative: guided workflow allowed controlled adaptation within established procedures, whereas hypothesis-driven exploration enabled strategies beyond those procedures when additional improvement was required. More broadly, these findings suggest that scientific agents should begin within harnessed workflows structured by expert knowledge and progressively acquire greater autonomy as reliable evidence accumulates, rather than operating with full autonomy from the outset (23–25).

The benchmark further clarifies the complementary roles of the adaptive processing levels. Guided workflow addressed most reconstruction-quality limitations through established classification, selection, and refinement procedures, whereas hypothesis-driven exploration provided an additional route for datasets that remained suboptimal after these procedures. The guided heterogeneity agent extended this structured adaptation toward structural-state discovery by recovering a previously overlooked 60S population independently supported by DRGN-AI. Thus, guided optimization provides an efficient solution for commonly encountered reconstruction problems, while hypothesis-driven exploration becomes particularly valuable when the encoded procedures are insufficient to achieve highest reconstruction quality.

Several major limitations remain. First, the current implementation is closely coupled with the cryoSPARC ecosystem and relies on its software interfaces, documentation, and Discussion Forum for tool execution and troubleshooting. Extending the framework to RELION, cisTEM, Scipion, and mixed-software workflows will be necessary to establish broader portability. Second, the external knowledge available to cryoAgent is currently dominated by the cryoSPARC Guide and Discussion Forum, which represents only a limited subset of available cryo-EM image-processing knowledge. Future systems should support broader retrieval from the scientific literature and enable agents to extract, compare, and operationalize processing strategies reported in previous studies. Third, and most importantly, cryoAgent interprets density maps indirectly through FSC, cFAR, cryoSift-derived measurements, similarity graphs, and other structured summaries. Although these representations support quantitative evaluation of global reconstruction quality, they may not capture subtle local defects that an experienced researcher could recognize visually. Models capable of identifying locally heterogeneous or poorly resolved regions could support the autonomous design of local masks and the targeted use of focused classification, local refinement, and multibody refinement.

cryoAgent suggests that reliable scientific-agent systems do not require the removal of established procedural constraints. Instead, expert-defined workflows can provide a dependable foundation from which autonomy is progressively expanded as informative intermediate evidence becomes available, allowing reproducible execution and open-ended reasoning to serve complementary roles. Although demonstrated here for image processing, this principle could ultimately be extended across the broader cryo-EM workflow, including automated data acquisition, real-time quality assessment, atomic model building, structural validation, and data deposition. Integrating these capabilities could provide a path toward a truly end-to-end autonomous structure-determination pipeline.

## Materials and Methods

### Thought–Action–Observation loop

All decision-making in cryoAgent follows a structured Thought–Action–Observation (TAO) protocol that couples language-model–based reasoning with external cryo-EM processing tools (Fig. 1a) (14). In each iteration, an agent receives a structured summary of the current workflow state, including job status, quality metrics, and log-derived signals, and generates a thought that describes the current state assessment and the next permissible operation. The thought process is explicitly constrained by the agent’s role and the set of actions supported by the underlying tools, ensuring that decisions remain grounded and reproducible.

Based on the selected action, the agent executes the corresponding tool operation and subsequently collects observations from the execution, including job state, numerical quality metrics (e.g., FSC and cFAR), and diagnostic information extracted from log files. These observations are normalized and fed back into the agent’s context for the next iteration. The TAO loop continues until predefined tssssermination criteria are met, such as task completion, convergence of quality metrics, or repeated failure, enabling adaptive yet interpretable control of long-running cryo-EM image processing workflows.

### Integration of cryo-EM software as agentic tools in cryoAgent

cryoAgent is designed to operate on top of widely adopted cryo-EM image-processing software rather than replacing existing, well-validated tools. In the current implementation, cryoAgent integrates cryoSPARC (26) as its primary processing engines. They are invoked programmatically through the APIs, allowing cryoAgent to submit jobs, monitor execution status, and retrieve intermediate results without manual intervention.

To enable systematic evaluation of intermediate and final processing results, cryoAgent also integrates additional analysis and assessment tools beyond the primary reconstruction engines. These tools provide structured quantitative observations that can be incorporated into the agent’s reasoning process, including decisions about class merging or splitting and convergence assessment. A complete list of the external tools integrated into cryoAgent, together with their software versions, is provided in Table S3.

By abstracting tool-specific formats and execution details into representations interpretable by the LLM, cryoAgent allows agent-level reasoning and decision-making to remain independent of the underlying processing software. This abstraction facilitates adaptive tool selection and cross-platform optimization within a unified multi-agent workflow.

### Memory representation of intermediate results

To support iterative reasoning across multi-stage cryo-EM workflows, cryoAgent employs differentiated memory strategies tailored to specific component requirements. The monitoring system maintains a dense contextual history of execution logs, warnings, and recovery actions, since effective recovery often depends on detailed log information and the recent execution context. Although long contexts may raise concerns about loss of earlier information, workflow failures are typically localized to a failed tool invocation and a few preceding steps. Consequently, recovery depends primarily on recent local context, and forgetting older context does not substantially reduce recovery performance (27).

In contrast, the heterogeneity and optimization systems utilize a compact structured memory strategy. Because these systems navigate long iterative trajectories that often involve tens of discrete tool executions, retaining the full raw logs would introduce significant noise and diminish reasoning efficiency. Instead, cryoAgent extracts high-signal observations, including quantitative metrics (e.g., resolution, cFAR, CryoSift scores, and terminal FSC profiles) and critical hyperparameters (e.g., box size, symmetry, and class number). These structured records function as a persistent working memory, enabling the agent to perform cross branch comparisons in heterogeneity system and track optimization convergence effectively. This design is consistent with prior agent-memory work showing that local decision-making benefits from retaining detailed recent processing history, whereas long-horizon reasoning is better supported by compact structured memory or hierarchical memory management to avoid context dilution (28).This strategy reduces unnecessary context and token usage while preserving the information required for subsequent decision-making.

### Conventional rigid automated workflow used as the procedural backbone

This rigid workflow is adapted from the cryoSPARC tutorial (26, 29), where the automated tools are used on essential steps. To define this procedural backbone clearly, we specify the parameter settings used at each stage of the fixed-step workflow.

During preprocessing, cryoSPARC-based workflows use Patch Motion Correction for movie alignment and Patch CTF Estimation for CTF determination. In RELION-based workflows, movies are processed using the RELION implementation of MotionCor2 (30), and CTF is estimated using CTFFIND4 (31). Micrographs are then filtered using a maximum fitted resolution threshold of 6 Å, which is the recommended value used by relion_it.py (7).

For particle picking, cryoAgent first uses blob picking to identify candidate particle locations. The particle diameter for picking is estimated from the box size of the corresponding EMDB entry minus 150 Å. The initial box size is set to match the deposited EMDB box size in angstroms. A first round of 2D classification is performed to generate templates for template-based picking.

Templates are selected using either a CryoSift score threshold of 3, corresponding to the default cutoffs used in the CryoSift (9). These templates are then used to identify high-quality particle locations. A second round of 2D classification is subsequently performed to enrich high-quality particles, which are again selected using the same CryoSift thresholds.

Finally, particles are reconstructed by ab initio reconstruction in cryoSPARC under C1 symmetry, followed by refinement using the deposited symmetry reported for the corresponding EMDB density map.

### Identifying the distinct type of states among densities

To determine how many distinct conformational states are represented in a single 3D classification run, the heterogeneity agent applies a post-classification clustering tool based on density similarity. Since 3D classification often yields overlapping or structurally redundant volumes, the agent first aligns all resulting density maps—using cryoAlign (17) and Fitmap (18) when no reference is provided—and computes pairwise FSC scores (Fig. S3a). FSC values at the 0.5 threshold are used to quantify similarity, from which a graph is constructed linking volumes with high similarity. Edges corresponding to differences greater than 20 Å are removed, as this is the more than the threshold for domain-level differences (Fig. S3b) (32). A community detection algorithm is then applied to this graph to estimate the number of distinct structural clusters present in that classification result (33). This provides an interpretable estimate of the essential number of non-redundant classes produced in a single round, independent of the redundancy of K value chosen.

### Agentic Heterogeneity Analysis via Recursive hierarchical Classification in guided heterogeneity analysis system

To determine overall dataset heterogeneity, the agent initiates a reasoning loop that uses the above clustering tool as its decision backbone. The process begins with 3D classification at K = 2 and K = 3. If the K = 3 classification is found to produce more distinct clusters than K = 2, the agent continues processing from K = 3 until the LLM determines that the process has converged (Fig. S4a). This logic is applied recursively. For each resulting cluster, the agent performs subclassification, again starting from K = 2 and K = 3, and evaluates structural diversity using the same graph-based clustering method and updates the plan for subsequent parameter selection (Fig. S4b). Subdivision continues until no further gain in heterogeneity is observed—at which point the agent assigns a cluster as a terminal “leaf.” After all branches have converged, the agent performs a final cross-leaf comparison to merge redundant structures, yielding a final set of distinct, high-confidence conformational states derived through adaptive, agentic reasoning (Fig. S4).

### Multi-round 2D optimization by the 2D optimization agent

To improve particle quality while minimizing the loss of valid signal, cryoAgent implements a dedicated 2D optimization agent that performs multiple rounds of 2D classification on extracted particle images (Fig. S5a). In each round, 2D classification is performed using a relaxed selection threshold relative to the picking process in the rigid workflow, allowing the agent to retain borderline classes that may contain meaningful signal but would otherwise be discarded in a single-pass workflow. After each round, the agent evaluates two complementary criteria derived from the classification results. First, it assesses changes in the distribution of particle quality scores using the median value and a Wilcoxon signed-rank test to determine whether the overall quality has significantly degraded relative to the previous iteration. If a statistically significant deterioration is detected, the agent interprets this as over-pruning and terminates further 2D optimization (Fig. S5b, S5c). Second, particularly during the first two rounds, when many particles remain junk and model-based contextual assessments may be less reliable, the agent applies an empirical safeguard based on the fraction of particles removed in each iteration (5). If fewer than 10% of particles are discarded, the agent issues a potential stop signal.

Rather than applying fixed termination rules across all datasets, the 2D optimization agent reasons over the context of the entire workflow. This allows the agent to adapt its stopping behavior across datasets with different characteristics, and to balance particle retention against quality improvement in a data-dependent manner. By integrating statistical evaluation with contextual reasoning, the 2D optimization agent enables robust, adaptive multi-round 2D refinement.

### Iterative reconstruction refinement by the 3D optimization agent

To further improve reconstruction quality beyond particle-level optimization, cryoAgent implements a dedicated 3D optimization agent that operates on three-dimensional density reconstructions and refinement results. Unlike the 2D optimization agent, which focuses on particle quality and class consistency in image space, the 3D optimization agent reasons over volumetric reconstructions and global and local quality metrics to guide iterative refinement at the reconstruction stage. Estimation of the true number of clusters is achieved by the heterogeneity agent, and the resulting heterogeneity analysis is subsequently used by the 3D optimization agent.

The 3D optimization agent decomposes iterative reconstruction refinement into three sequential stages, each targeting a distinct source of quality degradation. In the first stage, the agent performs multiple rounds of non-referenced heterogeneous refinement by agentic loop. This step is designed to identify and separate particles corresponding to junk protein, contaminants, or non-proteinaceous density that may not be captured by reference-guided classification. By avoiding structural bias at this stage, the agent enables robust removal of grossly inconsistent particle populations. In the second stage, the agent performs referenced heterogeneous refinement with an adaptively selected number of classes. Here, the agent treats 3D classification as a fine-grained sorting process, using reference-guided separation to further discard low-quality or poorly aligned particles while preserving structurally consistent populations. The number of classes in the above analysis is not a fixed priori but adjusted based on observed class separation and convergence behavior. In the final stage, the 3D optimization agent evaluates and adjusts the particle box size, which may be suboptimal in the initial processing workflow. When the box size is too small, CTF-delocalized signal extending beyond the extraction boundary is lost and cannot be fully recovered^34,35^. Conversely, unnecessarily large extraction boxes can include excessive background noise and, particularly in crowded micrographs, signal from adjacent particles, which can interfere with particle alignment and classification and thereby compromise reconstruction quality ^36^. To balance preservation of CTF-delocalized signal against unnecessary signal from surrounding image regions, the 3D optimization agent explores alternative box sizes around the baseline setting and evaluates their impact on reconstruction quality (Fig. S6a).

Across all three stages, agent decisions are guided primarily by the convergence of resolution, assessed through FSC trends across iterations (Fig. S6b). In addition, changes in cFAR are tracked as a complementary indicator of map quality; significant improvements in cFAR, particularly when the value is below 0.5, are explicitly recorded and considered evidence of meaningful quality improvement even when global resolution gains are modest (Fig. S6c). This staged, metric-driven strategy allows the 3D optimization agent to adaptively refine reconstructions while avoiding overfitting or unnecessary parameter exploration.

### Comparison of reconstruction quality across workflows

To compare reconstruction quality across different datasets (22, 37–39, 19, 40–54) with different processing strategies, we evaluated density maps generated by the full agentic workflow, the rigid automated image processing backbone, and the deposited reference densities. FSC, local resolution, and cFAR were calculated in cryoSPARC using the same evaluation procedure for densities produced by cryoAgent, densities produced by the rigid workflow, and deposited datasets with available half maps, ensuring consistency (55). When half maps for deposited densities were not available, direct recalculation of these metrics was not performed; instead, the reported FSC curves or resolutions provided with the deposited structures in EMDB were used for comparison (Fig. S14-S33). The local resolution for these deposited EMDB densities are calculated with ResMap using single map mode (56).

### External knowledge tools: cryoSPARC Discuss and the cryoSPARC Guide

CryoAgent uses two external knowledge sources in an advisory capacity, cryoSPARC forum and tutorial. For failure recovery, when a cryoSPARC job fails, the corresponding stage agent may search the cryoSPARC Discuss forum using error messages extracted from job.log. The agent retrieves the first five relevant discussion threads, identifies suggested causes or parameter changes, and uses this information to determine whether and how to retry the same job type. The retrieved information does not directly modify the workflow; rather, it is interpreted by the agent and translated into candidate recovery actions.

For the hypothesis generation, agents in the DE and HE systems consult the official cryoSPARC Guide and forum. The improvement agent first summarizes the current processing state and the strategies that have already been attempted, and then queries the knowledge for processing approaches relevant to that situation. The retrieved recommendations are interpreted in the context of the current dataset and converted into one or more processing hypotheses. Each hypothesis is subsequently translated into an executable plan consisting of a sequence of cryoSPARC jobs and associated parameters. After execution, the resulting reconstruction is compared with the current baseline using resolution and cFAR. If the result improves the evaluation metrics, it becomes the new baseline for the next iteration; otherwise, the attempted branch is archived as the context for further rounds.

### Evaluation of the performance of cryoWizard

EMPIAR-10181 was selected for presentation in the main figure, and cryoWizard was therefore run on this dataset to allow direct comparison with cryoAgent. The reconstruction quality of cryoWizard is better than the rigid workflow but worse than the cryoAgent full process. In the original publication of cryoWizard, 3 out of 6 datasets are available in the EMDB-China. We then use these three datasets for further direct comparison. For these three datasets, the resolution for cryoWizard 1.0 was directly taken from the original publication, whereas the performance of cryoWizard 1.5 was obtained from the cryoWizard GitHub README page. Density maps for comparison were downloaded from the Zenodo link provided by cryoWizard (8).

### Resource requirements and runtime estimation

cryoAgent was evaluated under two practical hardware configurations: an NVIDIA RTX 5090 GPU with SSD caching and an NVIDIA RTX 3090 GPU without SSD caching. These configurations represent computing resources that are accessible in typical laboratory settings. The estimated token consumption was approximately 3.5 million tokens per dataset on average, although the exact runtime and token usage varied by dataset. Runtime was measured from the initialization of each agent to the generation of its summary report, therefore it included both LLM reasoning and tool execution (Tables S4 and S5).

### Statistical analysis and exact P values

Because the normal distribution of the differences between paired process methods cannot be assumed, statistical comparisons were performed using two-sided paired Wilcoxon signed-rank tests. For reconstruction resolution, comparisons were made among the deposited reconstruction, the rigid automated workflow, the rigid workflow with 2D optimization, and full cryoAgent optimization; the exact P values were as follows: deposited versus rigid, P = 0.0484; deposited versus rigid+2D, P = 0.398; deposited versus full cryoAgent, P = 0.000322; rigid versus rigid+2D, P = 0.0347; and rigid+2D versus full cryoAgent, P = 0.000131. For cFAR, comparisons were made among the rigid automated workflow, the rigid workflow with 2D optimization, and full cryoAgent optimization; the exact P values were as follows: rigid versus rigid+2D, P = 0.354; rigid versus full cryoAgent, P = 0.000577; and rigid+2D versus full cryoAgent, P = 0.00055. These exact P values are provided for transparency and to facilitate direct interpretation of the magnitude of the statistical evidence.

## Code availability

The code is available at github.com/smallelephant9516/cryoAgent

## Data availability

The movies and micrographs analyzed in this study are available through EMPIAR (https://www.ebi.ac.uk/empiar). The RELION tutorial dataset is available at the RELION tutorial website (https://relion.readthedocs.io/en/latest/SPA_tutorial/Introduction.html). The cryoSPARC tutorial dataset and associated processing parameters are available at the cryoSPARC introductory tutorial website (https://guide.cryosparc.com/processing-data/get-started-with-cryosparc-introductory-tutorial). Deposited density maps are available through EMDB (https://www.ebi.ac.uk/emdb/). Final density maps generated by cryoAgent could be downloaded at https://zenodo.org/records/19348651.

## Acknowledgments

We thank Yingying Zhu at the Guangzhou Institutes of Biomedicine and Health, Chinese Academy of Sciences, and Renmin Han at Shandong University for valuable discussions regarding this project. This work is supported by National Natural Science Foundation of China (32525005 to F.S.), National Key Research and Development Program of China (2021YFA1301500 to F.S., 2021YFF0704300 to Yan Z., 2024YFA1307402 to Yun Z.), Basic Research Program Based on Major Scientific Infrastructures, CAS (JZHKYPT-2021-05 to F.S.)

## Supplementary Data

**Figure S1.**
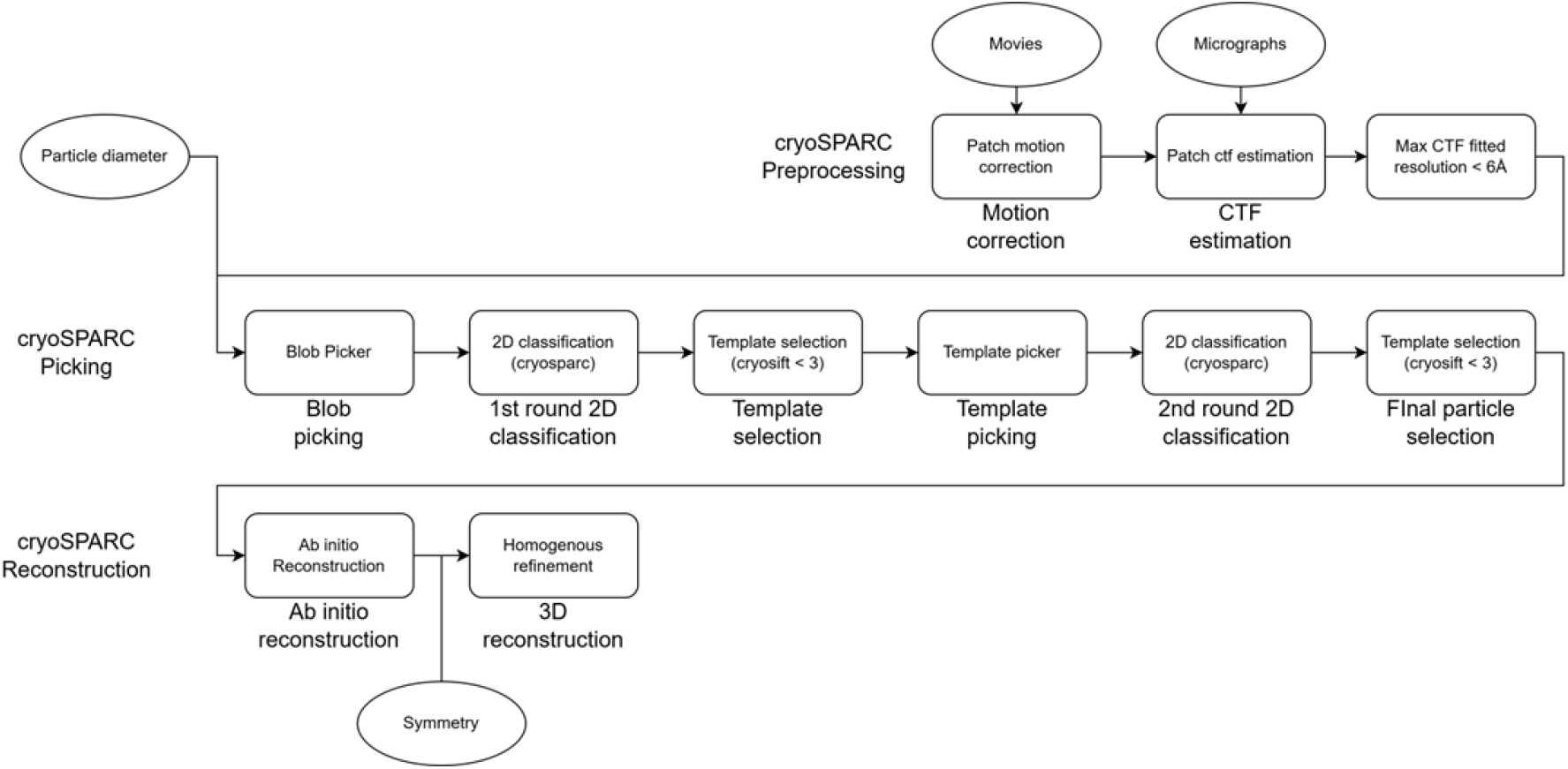
The tool and parameter usage in the rigid workflow.

**Figure S2.**
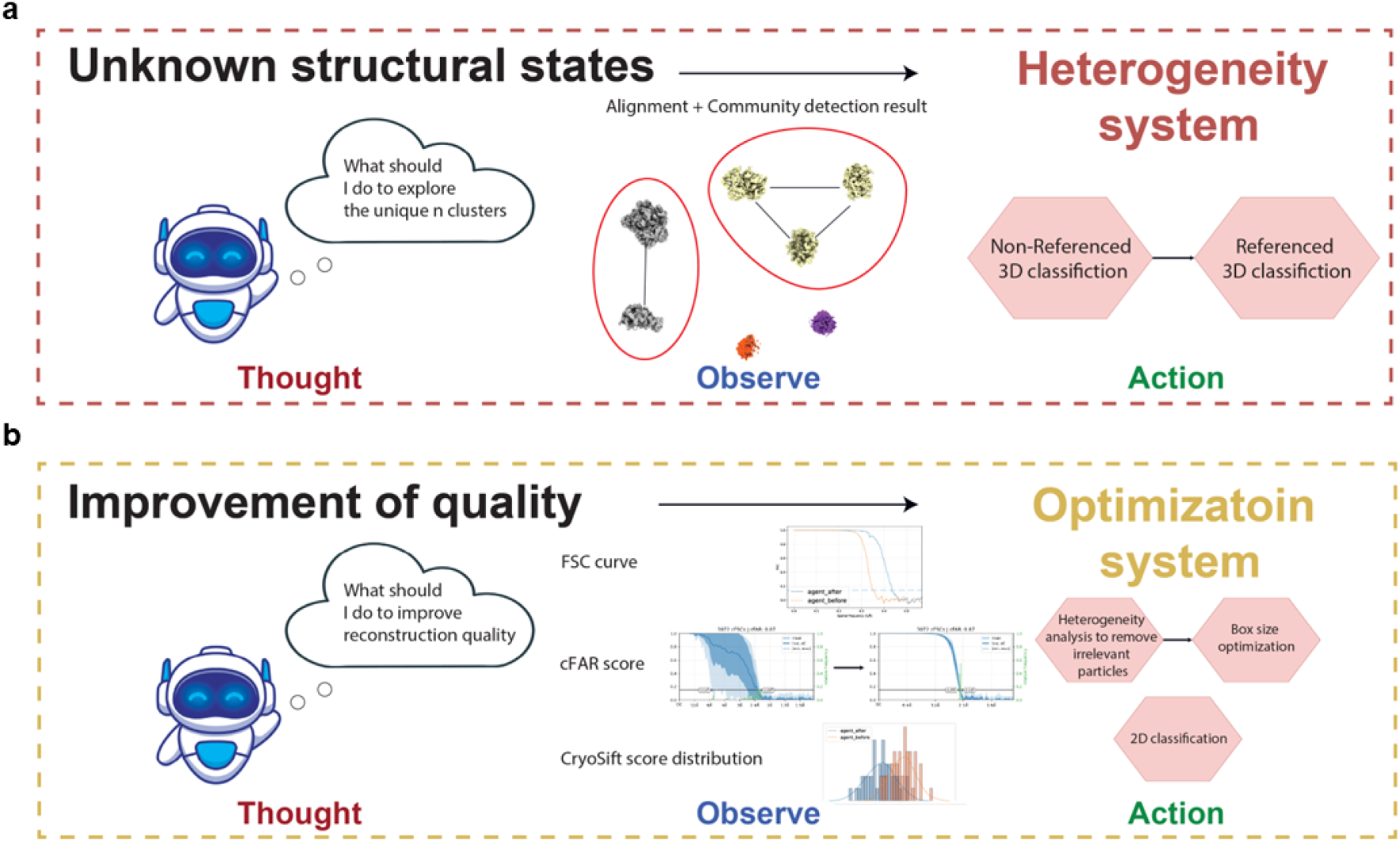
The systems in the guided workflow A, In the heterogeneity system, the agent inspects the similarity graph from the heterogeneity analysis and determines the essential number of structural states. B, In the optimization system, the agent iteratively removes junk particles through 2D/3D classification and optimizes box size to maximize the reconstruction quality.

**Figure S3.**
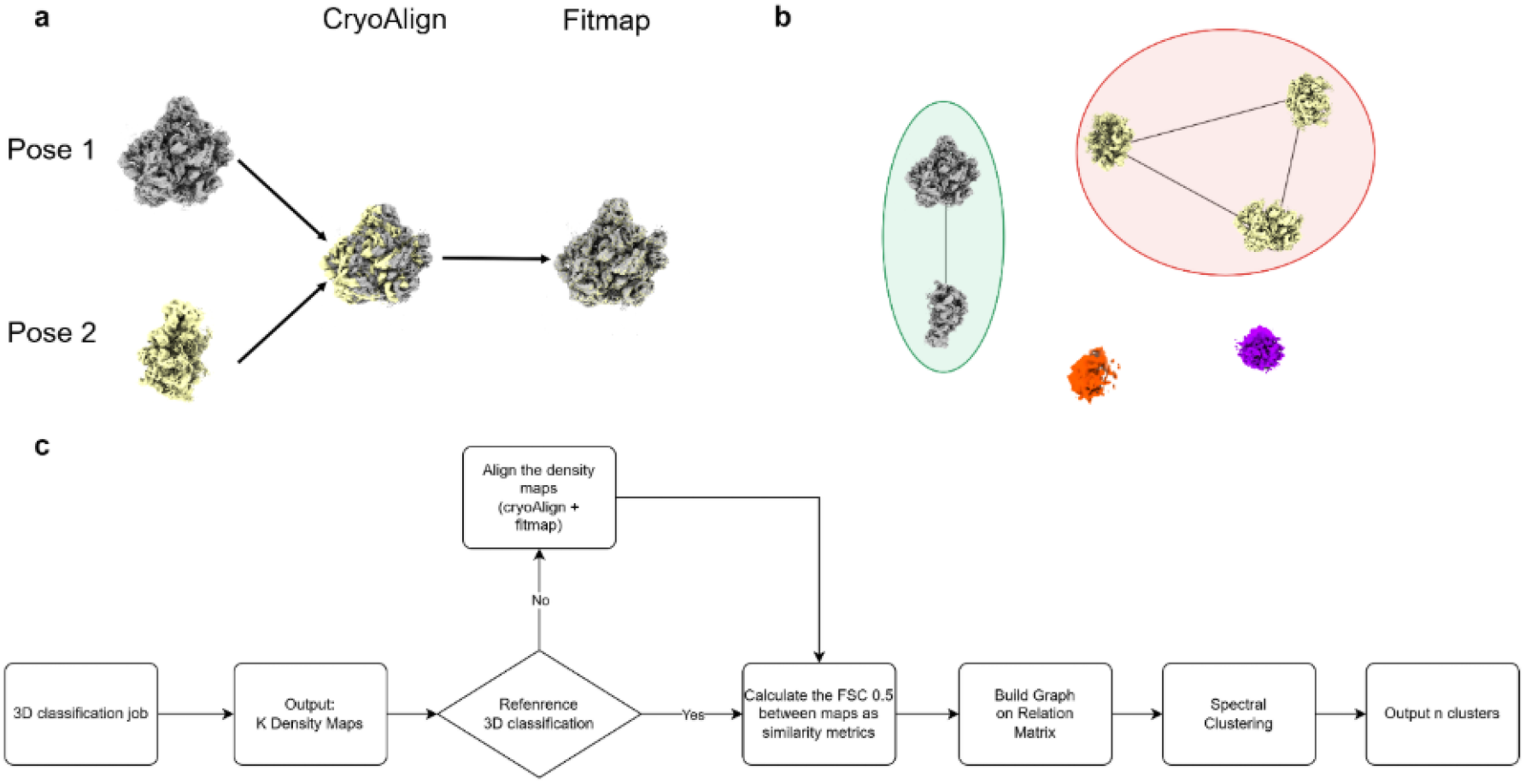
Alignment and clustering of density maps from 3D classification. A, Schematic illustration of density-map alignment using a combination of cryoAlign and fitmap to align two reconstructed volumes. B, Graph constructed from the pairwise similarity matrix of aligned density maps, with density types identified using graph-based community detection. C, Conceptual diagram of the graph-theoretic framework used to estimate the number of unique structural clusters.

**Figure S4.**
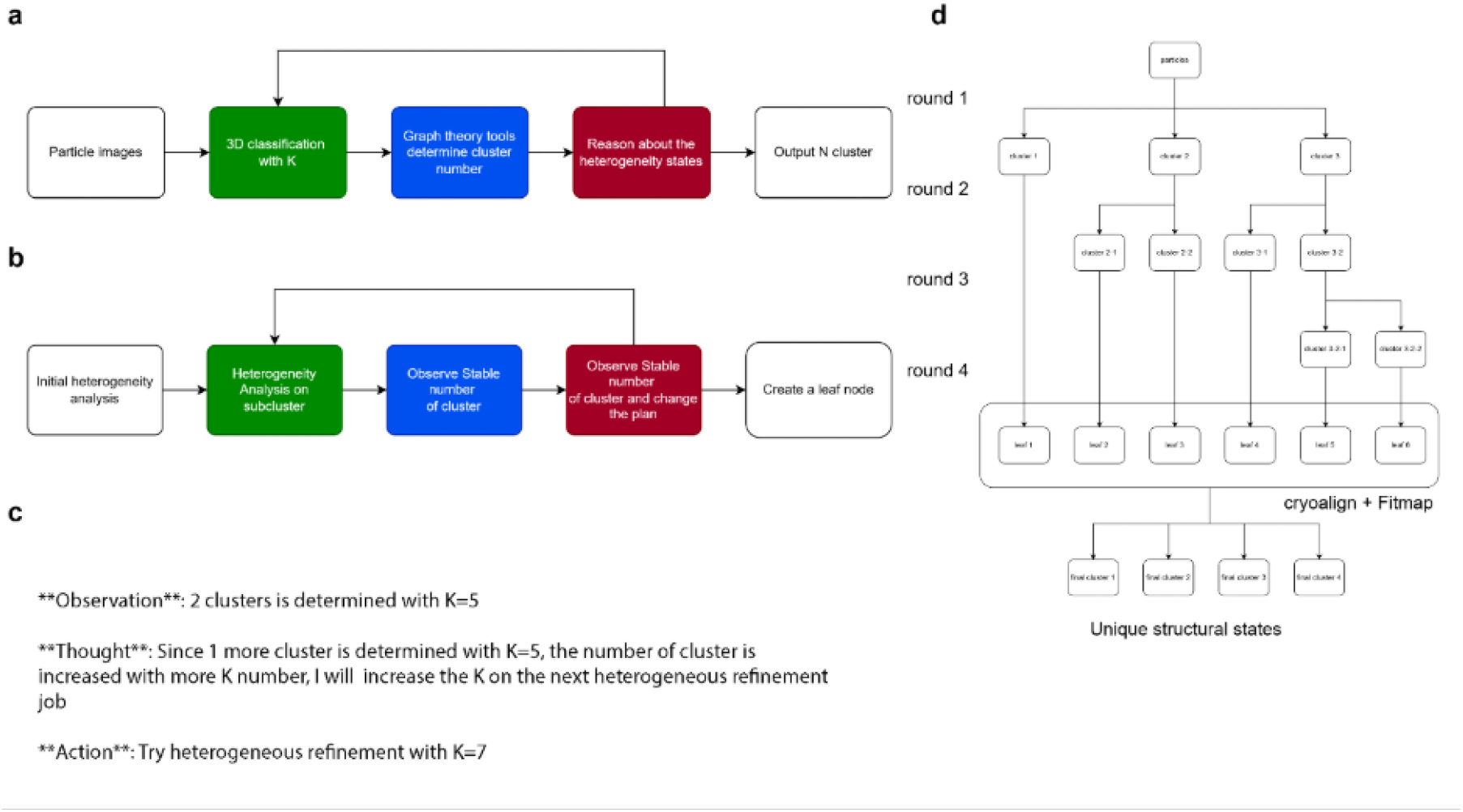
Agentic process for iterative heterogeneity analysis. a, Reasoning logic used by the language model to determine the classification number K for a single round of heterogeneity analysis. b, Reasoning process for identifying stable clusters and creating leaf nodes in the hierarchical, recursive agentic workflow. c, Reasoning and response generated by the language model in response to the detected number of unique clusters. d, Hierarchical tree generated by the heterogeneity agent. Leaf nodes are compared using cryoAlign and fitmap to assess pairwise similarity and determine the final number of unique structural states.

**Figure S5.**
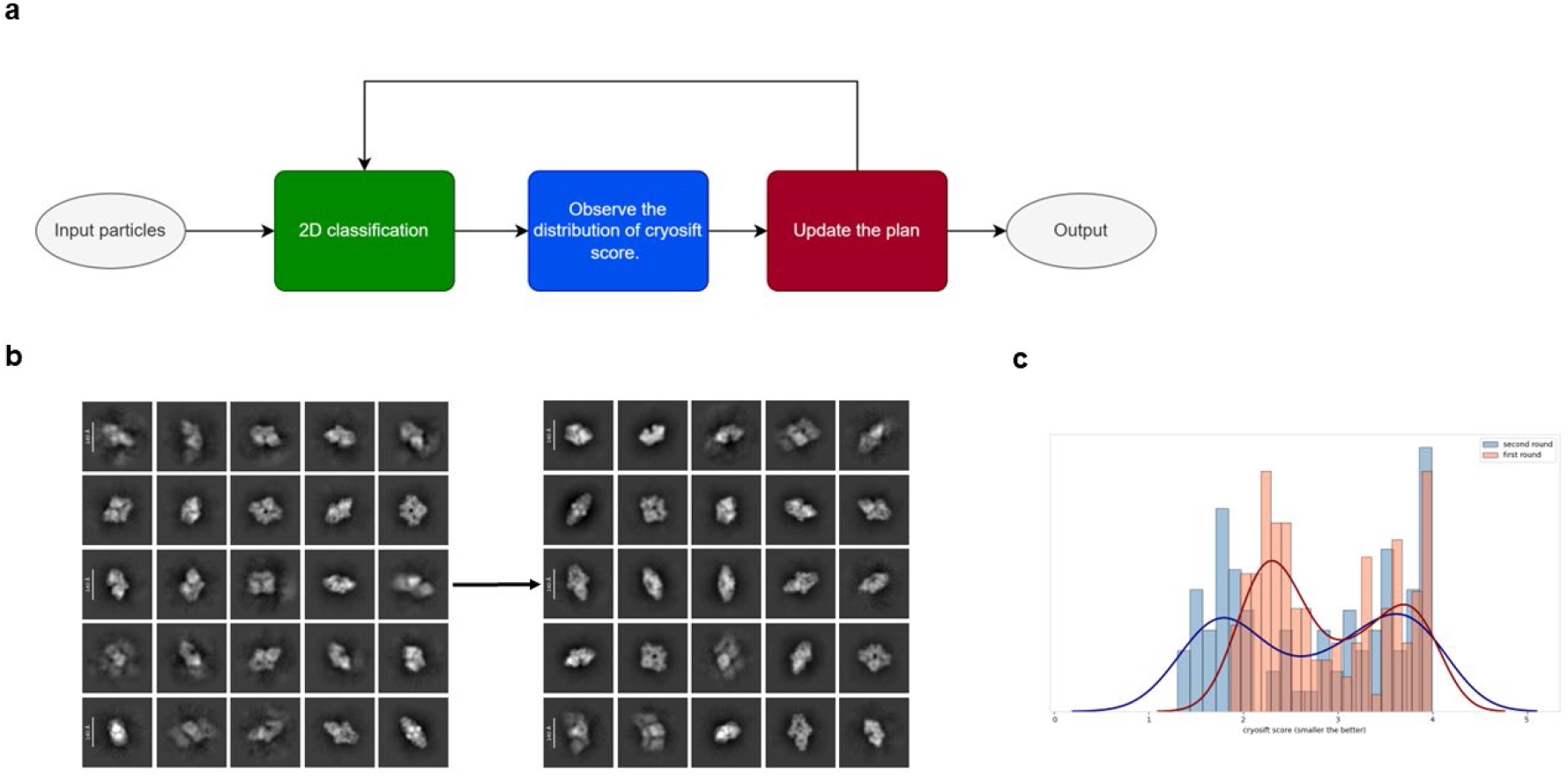
Agentic process for 2D optimization. A, Reasoning logic used by the language model to determine whether the quality of 2D classification has converged and whether additional rounds are required. B, Representative 2D classes selected after the first round (left) and second round (right) of agent-driven 2D classification on RELION tutorial dataset. C, Distribution of Cryosift scores for particles after the first and second rounds of 2D classification of the 2D classification result in b.

**Figure S6.**
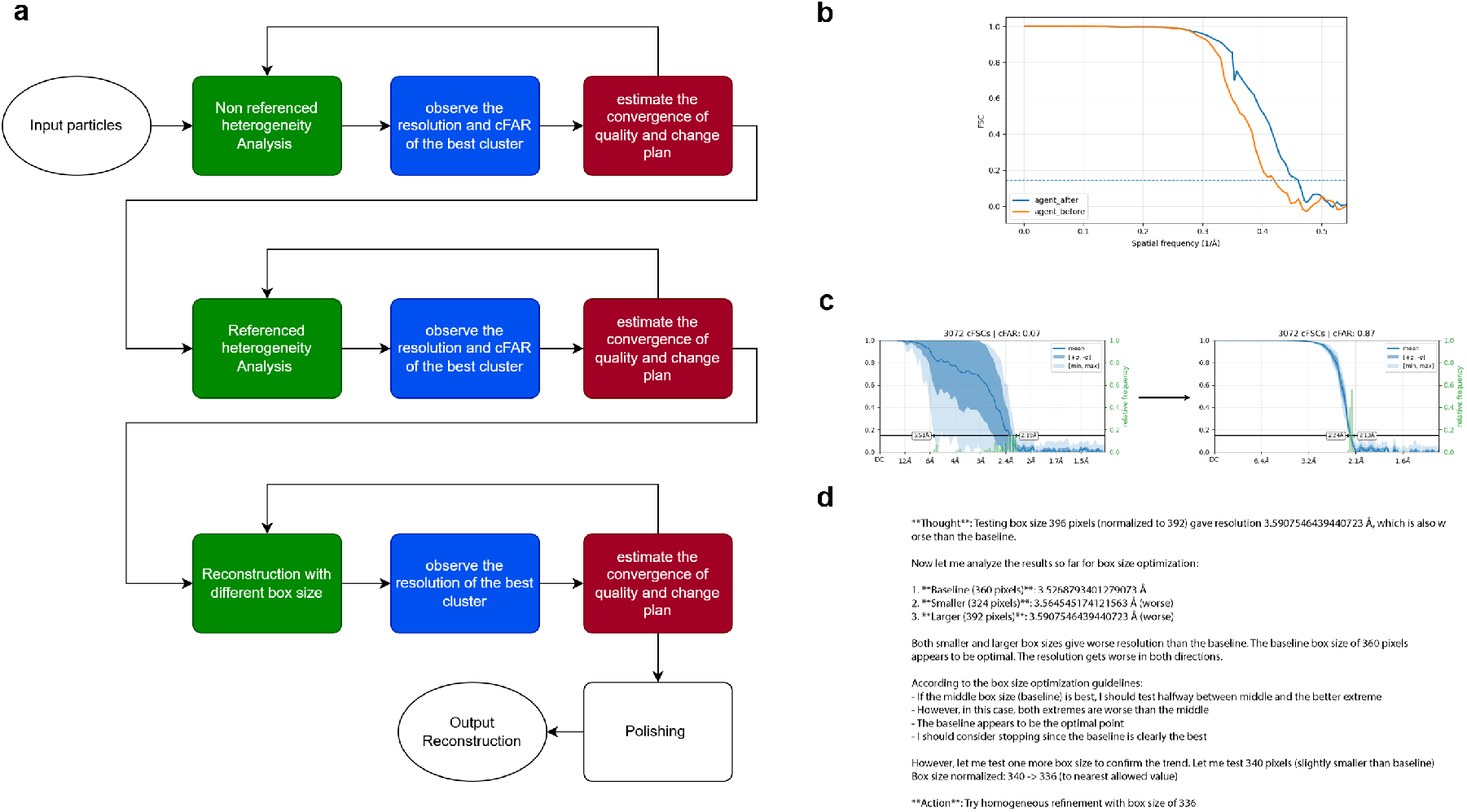
Agentic process for multiple rounds of 3D classification and box size optimization. a, Reasoning logic used by the language model to determine whether heterogeneity analysis and box-size optimization have converged and whether additional rounds are required. b, Representative changes in FSC between the first and second rounds of heterogeneity analysis. c, Representative changes in cFAR score between the first and second rounds of heterogeneity analysis. d, Reasoning and response generated by the optimization agent in reaction to changes in cFAR and resolution metrics.

**Figure S7.**
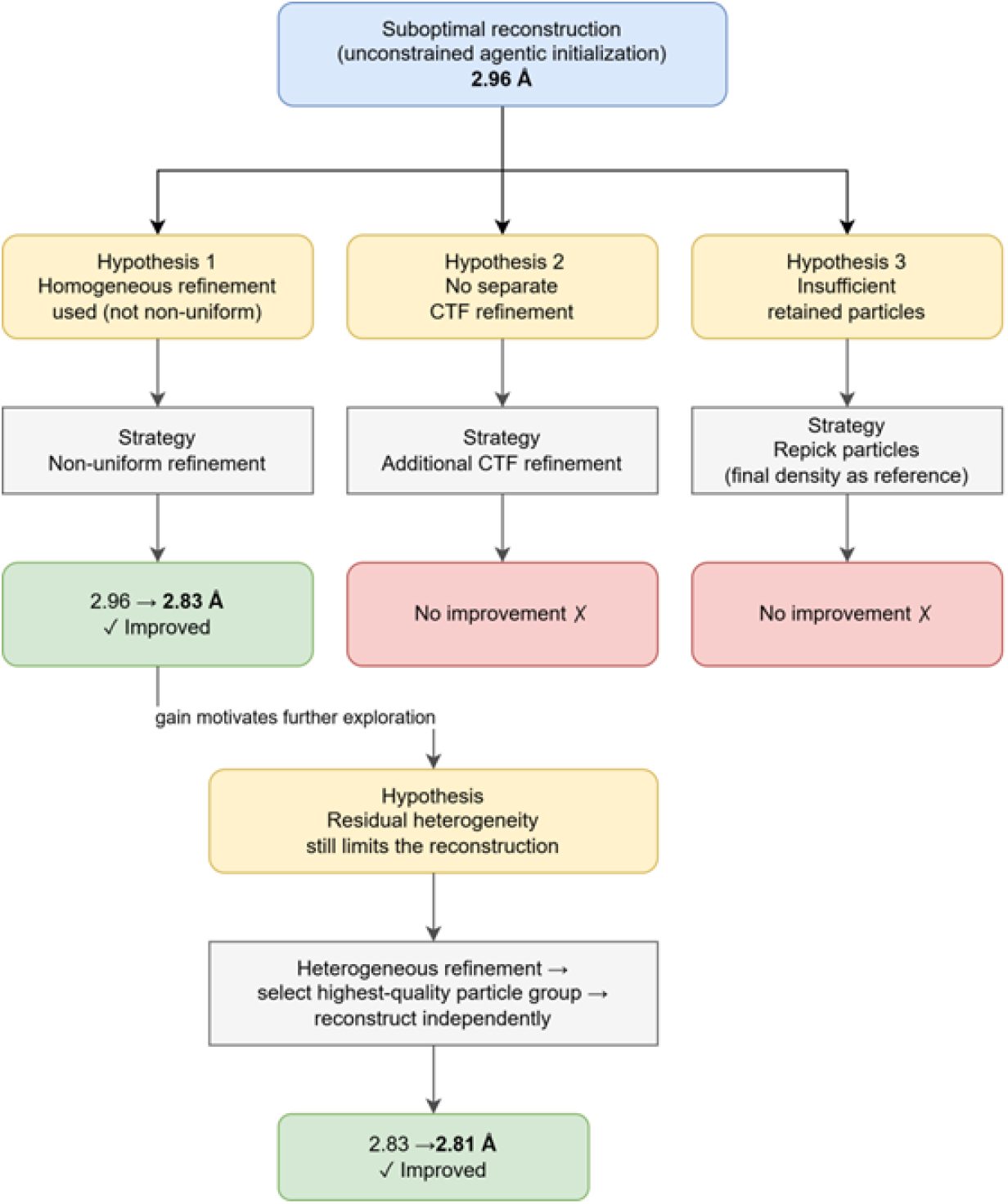
Hypothesis-driven exploration trajectory initialized from the dynamic-exploration result on the RELION tutorial dataset. The workflow shows the exploration path followed by the hypothesis-driven exploration mode after initialization from the reconstruction generated by dynamic exploration on the RELION tutorial dataset.

**Figure S8.**
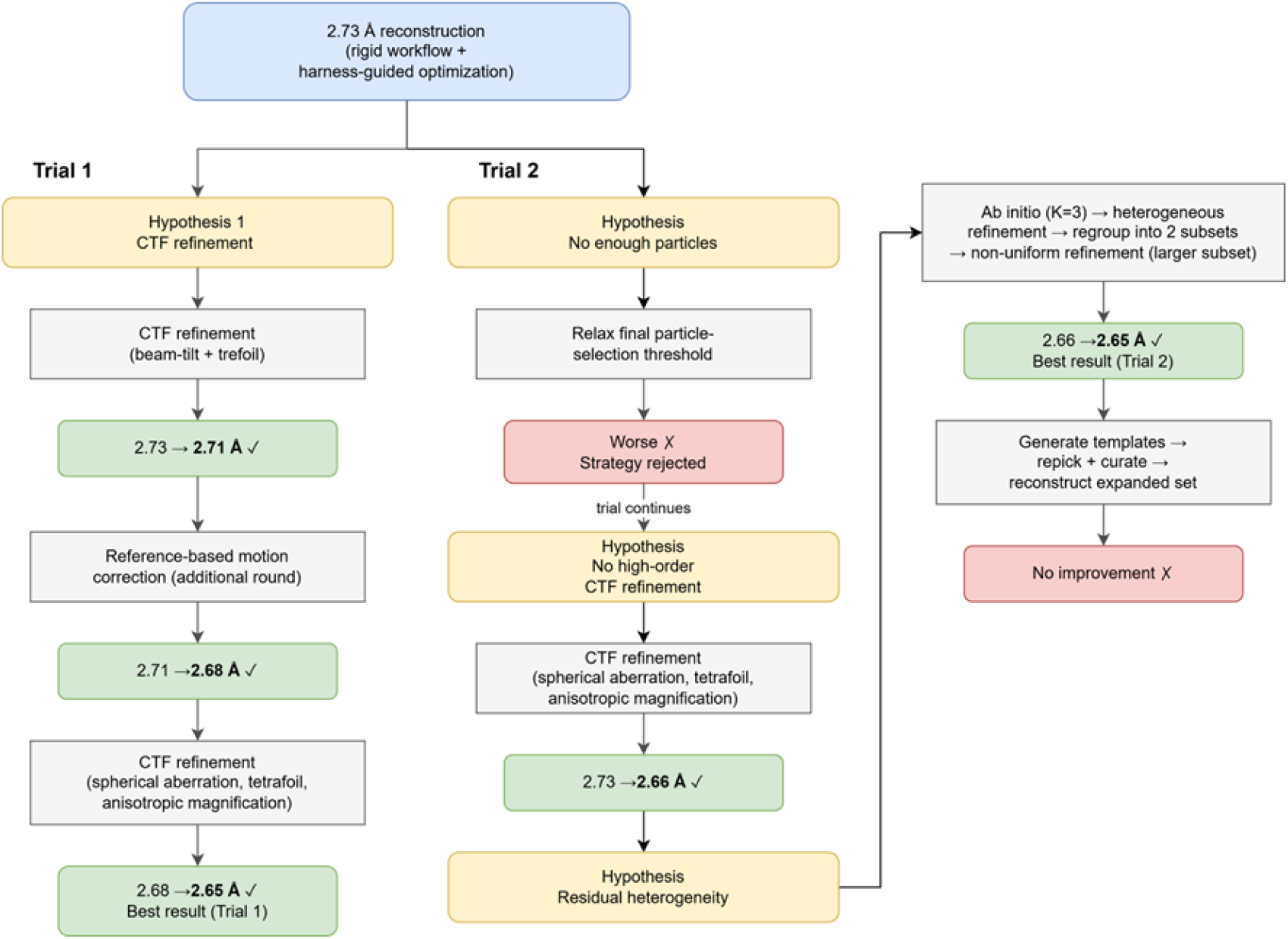
Two hypothesis-driven exploration trials initialized from the guided optimization result on the RELION tutorial dataset. The two trials show distinct exploration paths followed by the hypothesis-driven exploration mode when starting from the reconstruction generated by guided optimization on the RELION tutorisal dataset.

**Figure S9.**
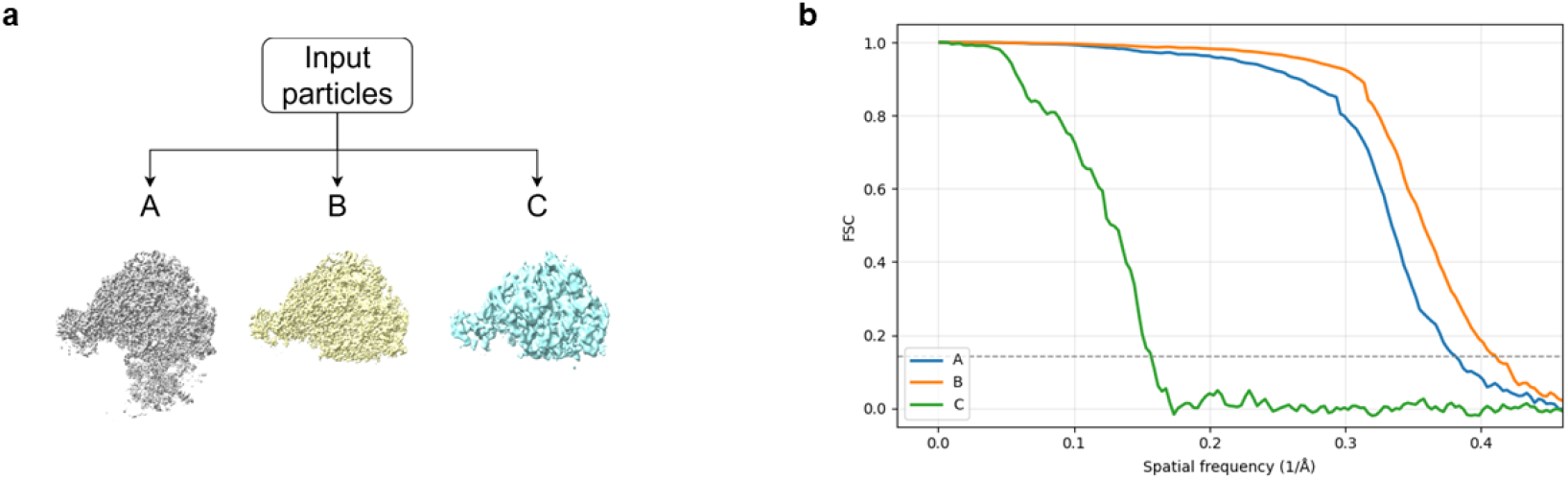
Discovery tree structure of EMPIAR-10406 generated by the heterogeneity agent. a, Hierarchical branching of subsets for EMPIAR-10406 produced by the heterogeneity agent. Input particles are recursively partitioned into progressively refined subpopulations, resulting in multiple structurally distinct reconstructions. b, FSC curves of reconstructions from different branches of EMPIAR-10406 dataset.

**Figure S10.**
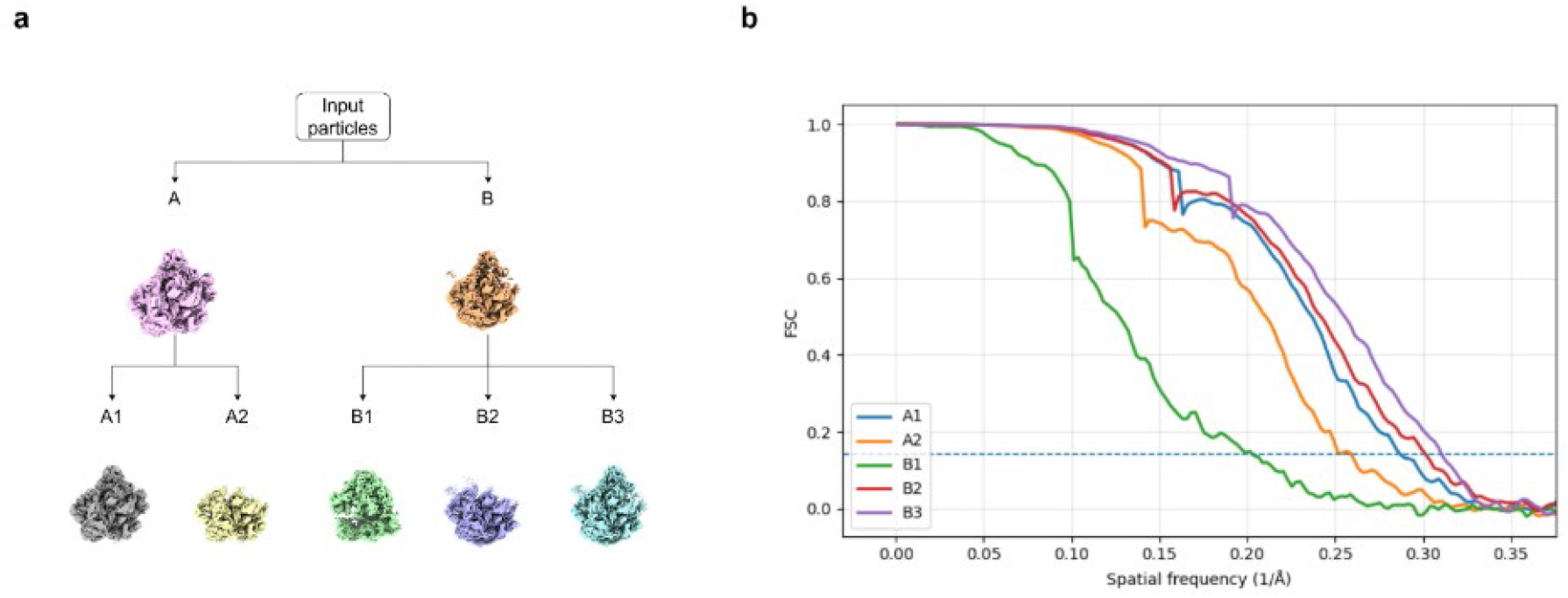
Discovery tree structure of EMPIAR-10076 generated by the heterogeneity agent. a, Hierarchical branching of subsets for EMPIAR-10076 produced by the heterogeneity agent. Input particles are recursively partitioned into progressively refined subpopulations, resulting in multiple structurally distinct reconstructions. b, FSC curves of reconstructions from different branches of EMPIAR-10076 dataset.

**Figure S11.**
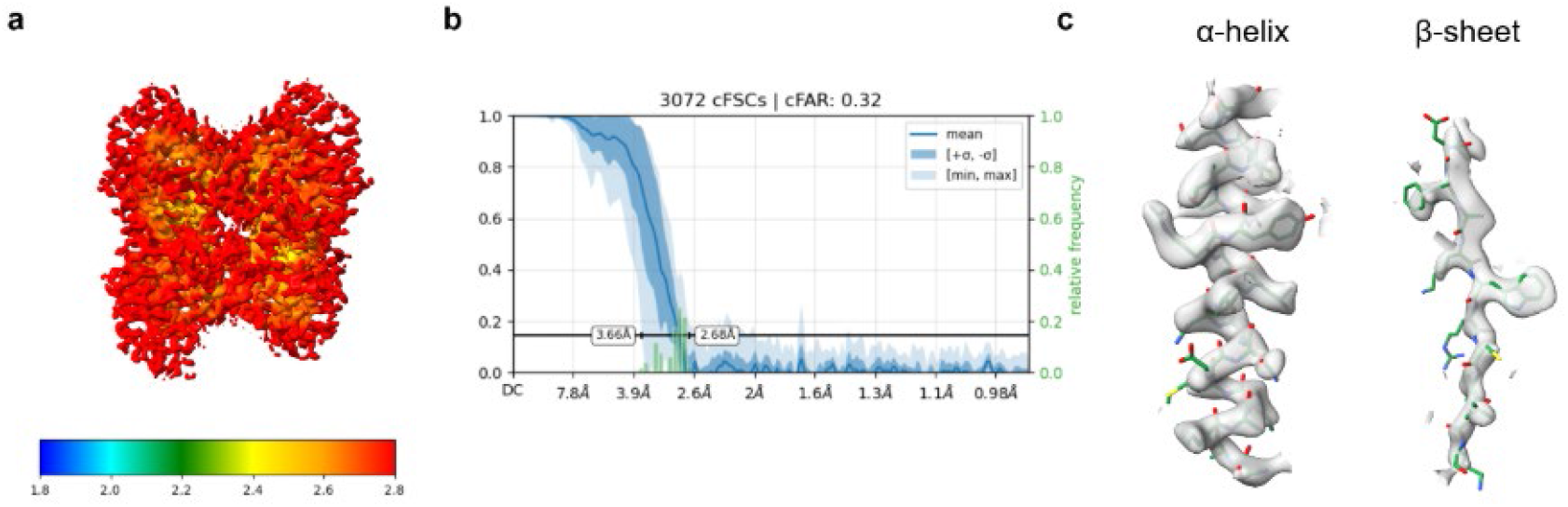
Illustration of the reconstruction results produced by the cryoWizard on the EMPIAR-10181 dataset a, Density map from cryoWizard default mode. b, cFAR score panel of the density map from the cryoWizard default mode c, Representative of the same α-helices and β-sheets region as Fig.5a-5b are shown;

**Figure S12.**
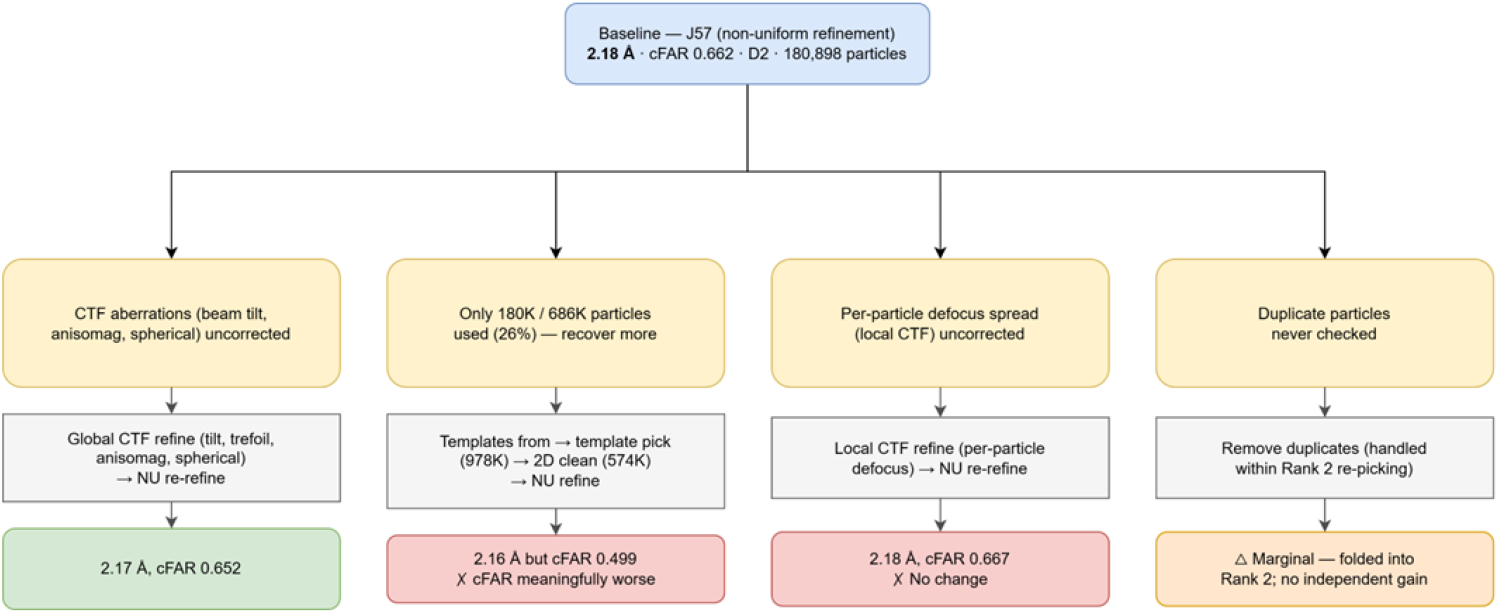
Hypothesis-driven exploration modes from the guided optimization result on the EMPIAR-10181 dataset. The workflow shows the exploration path followed by the hypothesis-driven exploration mode after initialization from the reconstruction generated by guided optimization on the EMPIAR-10181 dataset.

**Figure S13.**
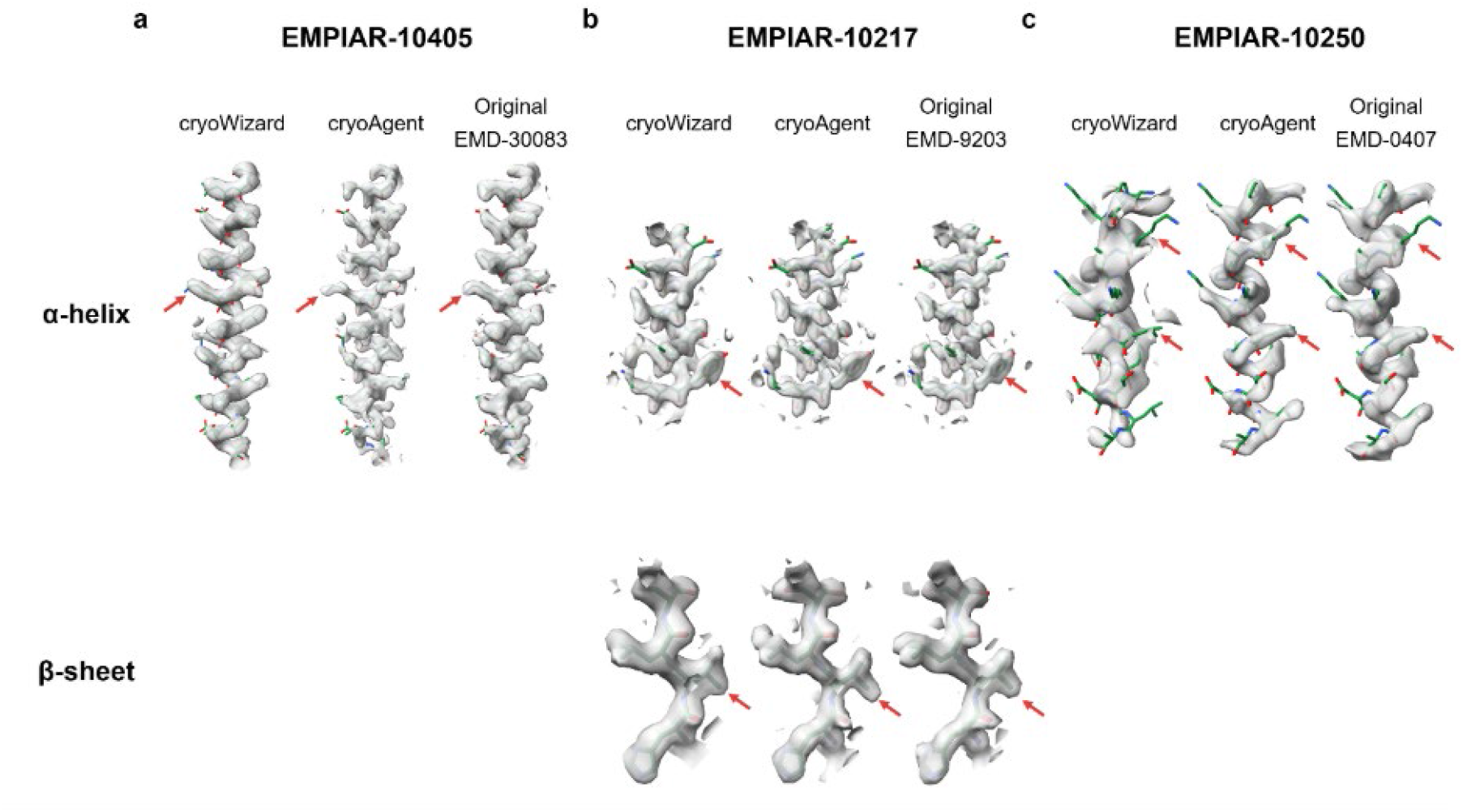
Comparison of local density quality between deposited cryoWizard reconstructions, corresponding EMDB entries, and densities produced by cryoAgent. Representative regions are shown for (a) EMPIAR-10405, (b) EMPIAR-10217, and (c) EMPIAR-10250. For each dataset, regions containing α-helices and β-sheets are compared across densities produced by cryoWizard, deposited in EMDB, and generated by cryoAgent. Red markers indicate residues with substantially improved local density quality.

**Figure S14.**
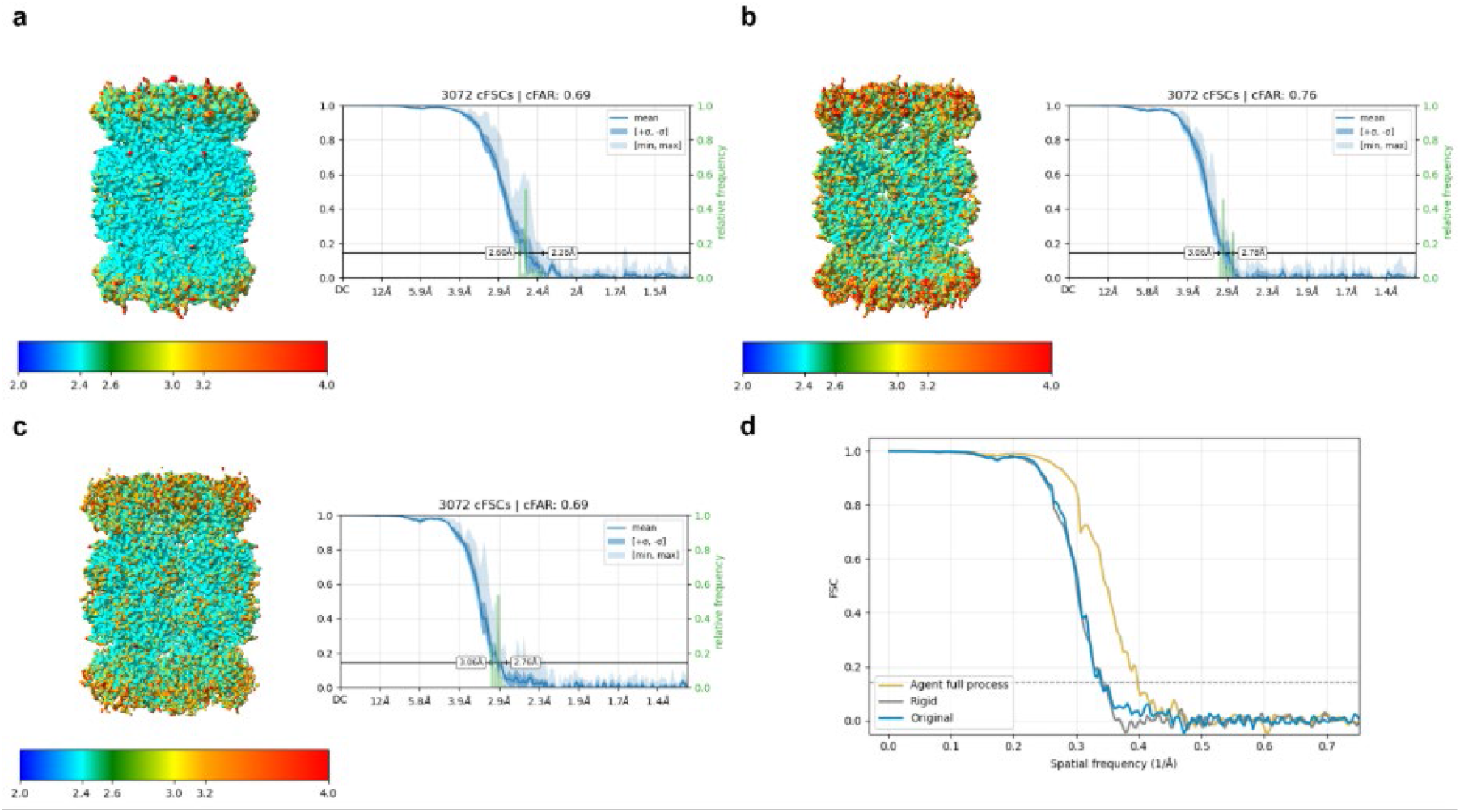
Comparison of reconstructions of T20S proteasome produced by different processing workflows on cryoSPARC tutorial dataset (38)**. a–c,** Density maps from the full cryoAgent workflow (a), conventional rigid workflow (b), and manual image processing with cryoSPARC tutorial parameters (c), shown with corresponding local resolution and cFAR score maps. **d,** FSC curves comparing reconstructions obtained from the full cryoAgent workflow, the conventional rigid automated workflow, and cryoSPARC automatic processing for the T20S with tutorial parameters.

**Figure S15.**
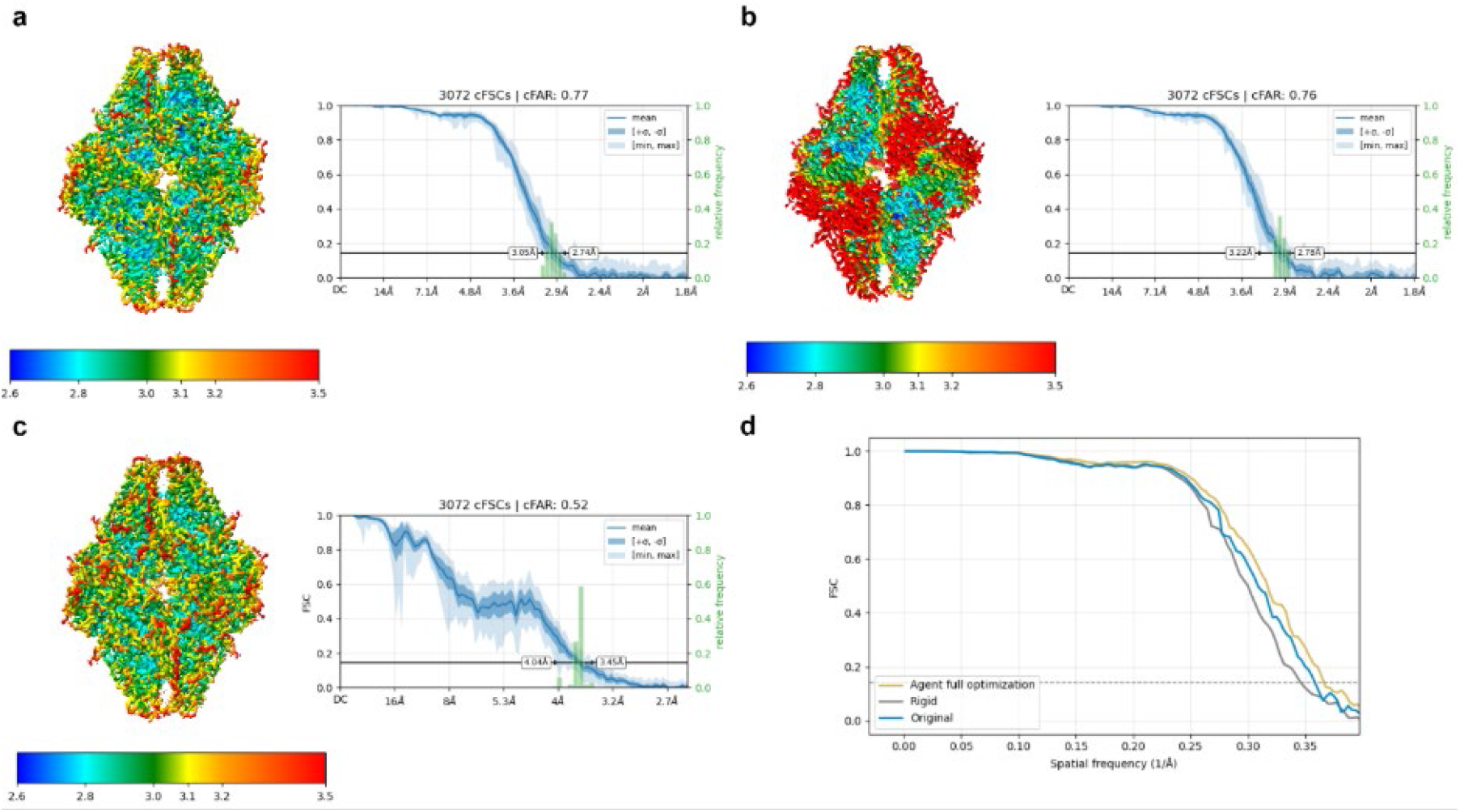
Comparison of reconstructions of β-galactosidase produced by different processing workflows on RELION tutorial dataset(39). a–c, Density maps from the full cryoAgent workflow (a), conventional rigid workflow (b), and cryoSPARC automatic processing (c), shown with corresponding local resolution and cFAR score maps. d, FSC curves comparing reconstructions obtained from the full cryoAgent workflow, the conventional rigid automated workflow, and the final refined density from the RELION 4 tutorial workflow.

**Figure S16.**
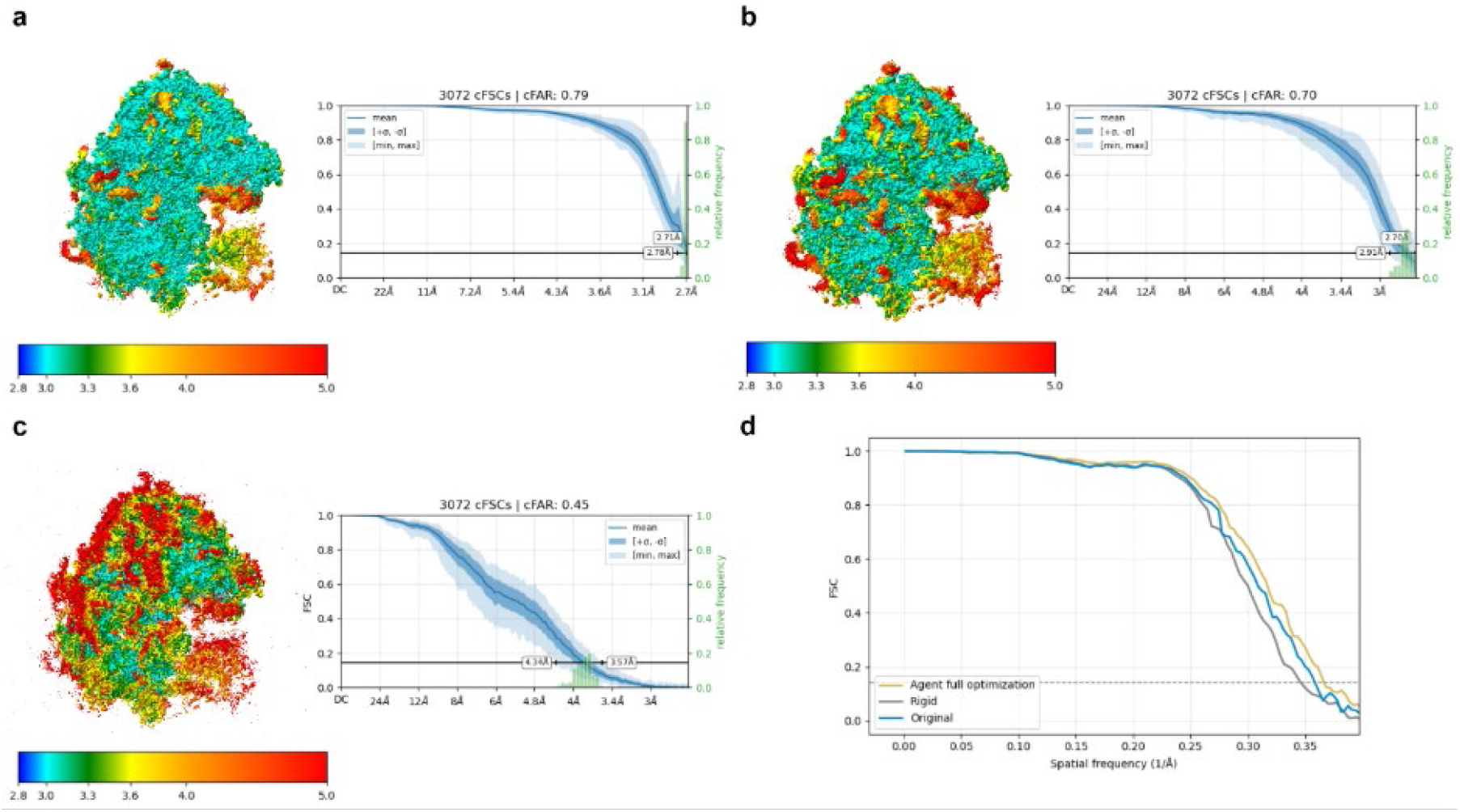
Comparison of reconstructions of Plasmodium falciparum 80S ribosome produced by different processing workflows on EMPIAR-10028 dataset(19). a–c, Density maps from the full cryoAgent workflow (a), conventional rigid workflow (b), EMD-2660 (c), shown with corresponding local resolution and cFAR score maps. d, FSC curves comparing reconstructions obtained from the full cryoAgent workflow, the conventional rigid automated workflow, and the corresponding EMDB density.

**Figure S17.**
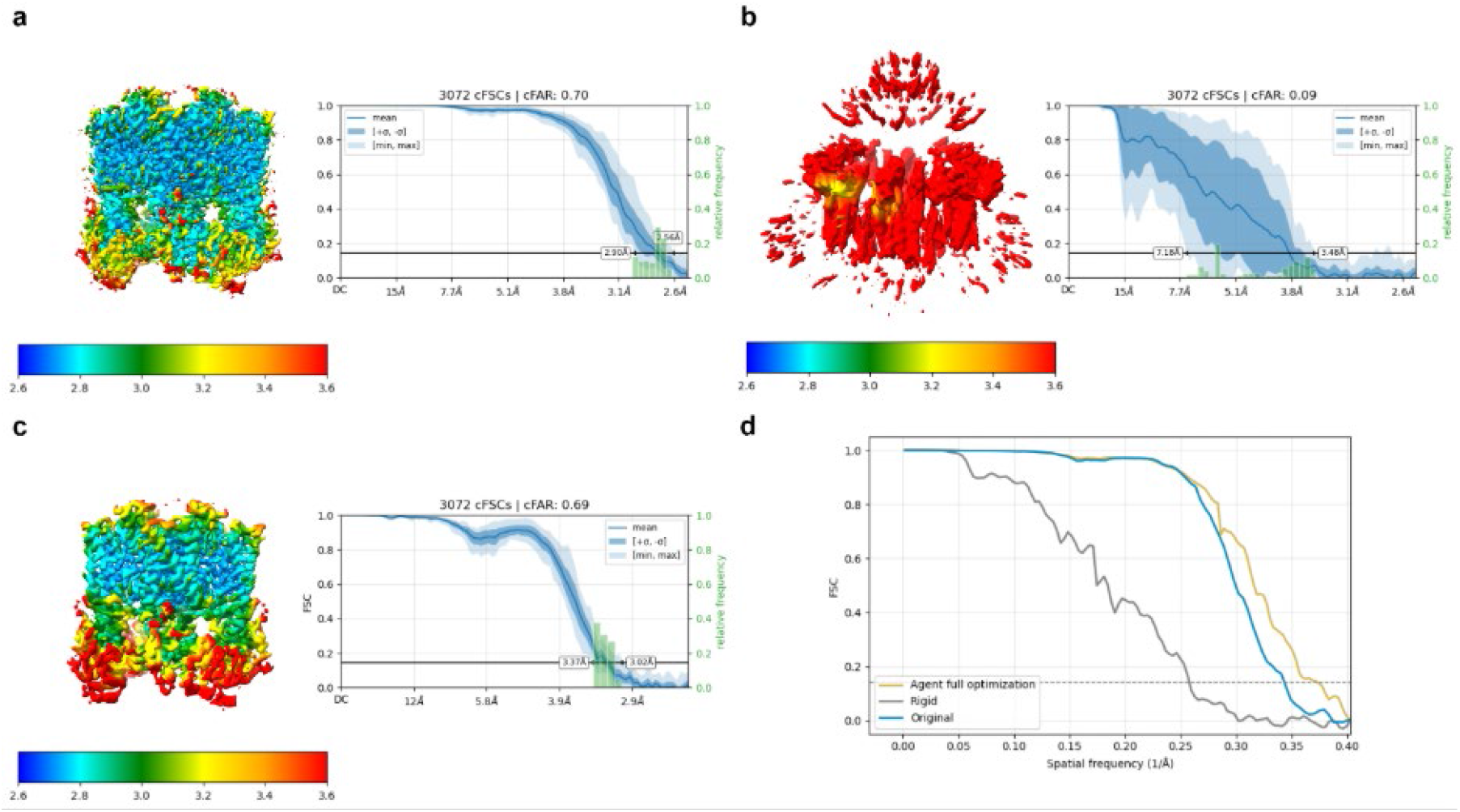
Comparison of reconstructions of TRPV1 produced by different processing workflows on the EMPIAR-10059 dataset(40). a–c, Density maps from the full cryoAgent workflow (a), conventional rigid workflow (b), and EMD-8117 (c), shown with corresponding local resolution and cFAR score maps. d, FSC curves comparing reconstructions obtained from the full cryoAgent workflow, the conventional rigid automated workflow, and the corresponding EMDB density.

**Figure S18.**
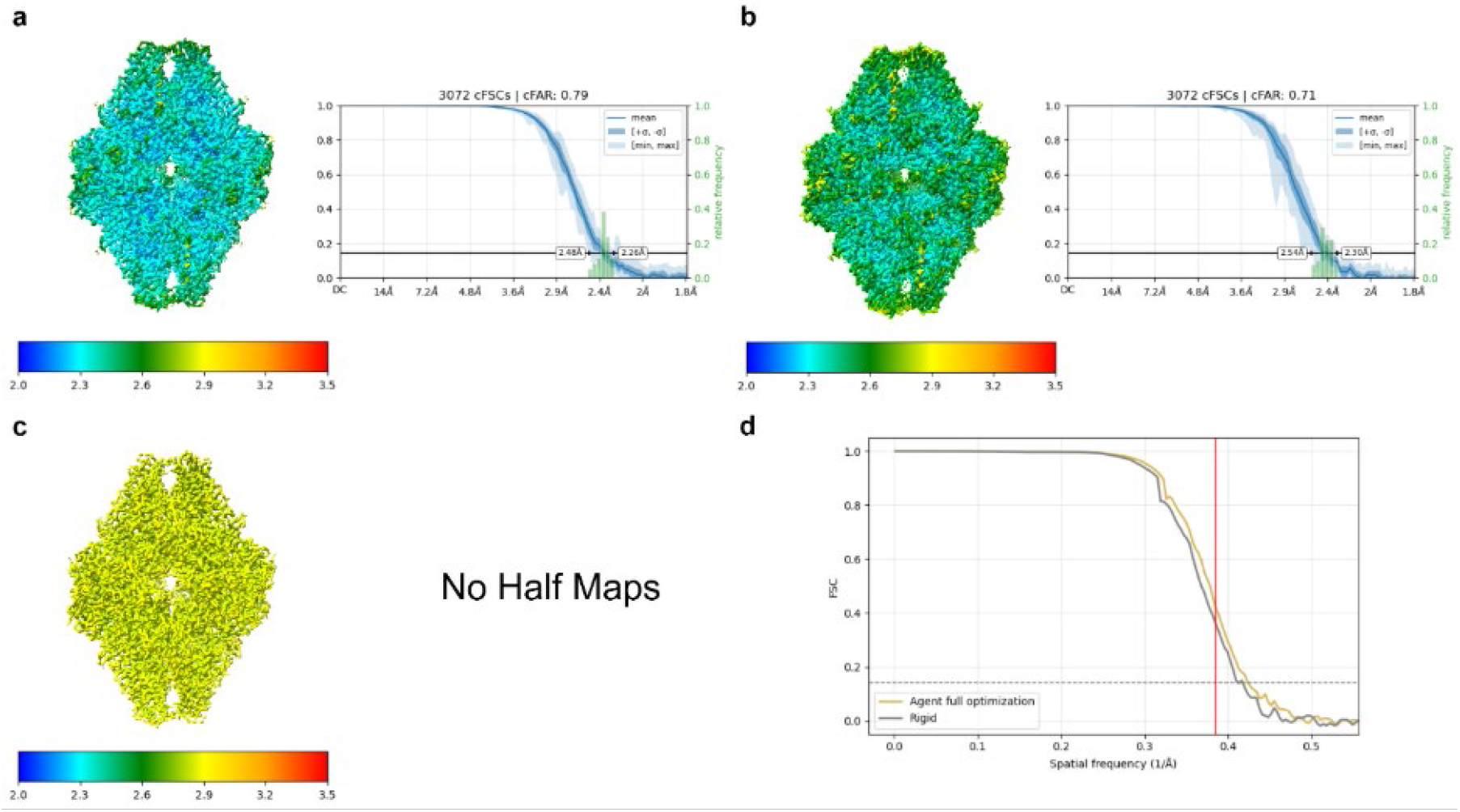
Comparison of reconstructions of β-galactosidase produced by different processing workflows on the EMPIAR-10204 dataset(41). a–b, Density maps from the full cryoAgent workflow (a) and conventional rigid workflow (b) shown with corresponding local resolution and cFAR score maps. c, EMD-6840 density map. As the half-maps were not available, local resolution was estimated using ResMap in single-map mode, and cFAR was not evaluated. d, FSC curves comparing reconstructions obtained from the full cryoAgent workflow, the conventional rigid automated workflow. The red line indicates the deposited resolution for the corresponding EMDB density.

**Figure S19.**
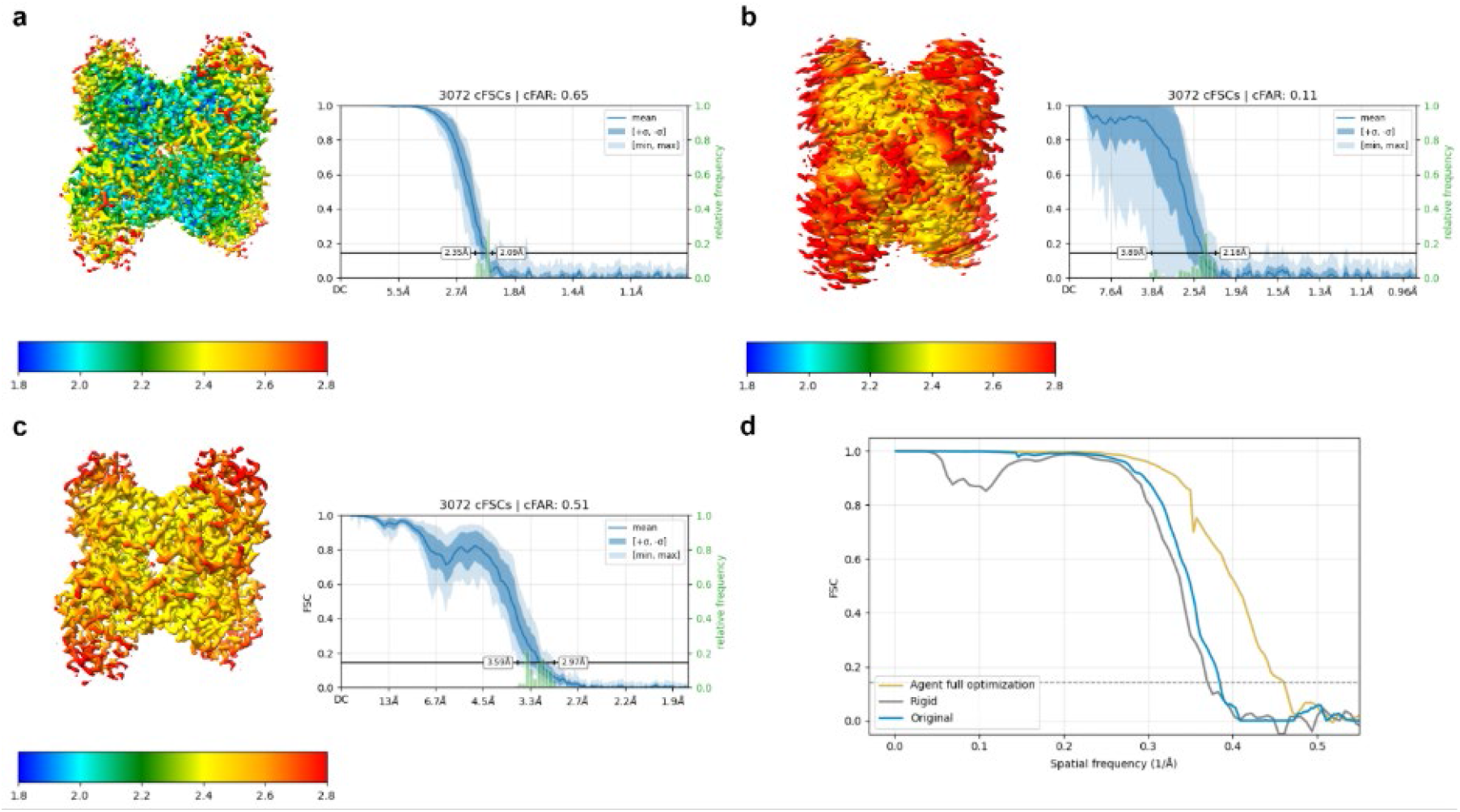
Comparison of reconstructions of Rabbit Muscle Aldolase produced by different processing workflows on the EMPIAR-10181 dataset(22). a–c, Density maps from the full cryoAgent workflow (a), conventional rigid workflow (b), and EMD-8743 (c), shown with corresponding local resolution and cFAR score maps. d, FSC curves comparing reconstructions obtained from the full cryoAgent workflow, the conventional rigid automated workflow, and the corresponding EMDB density.

**Figure S20.**
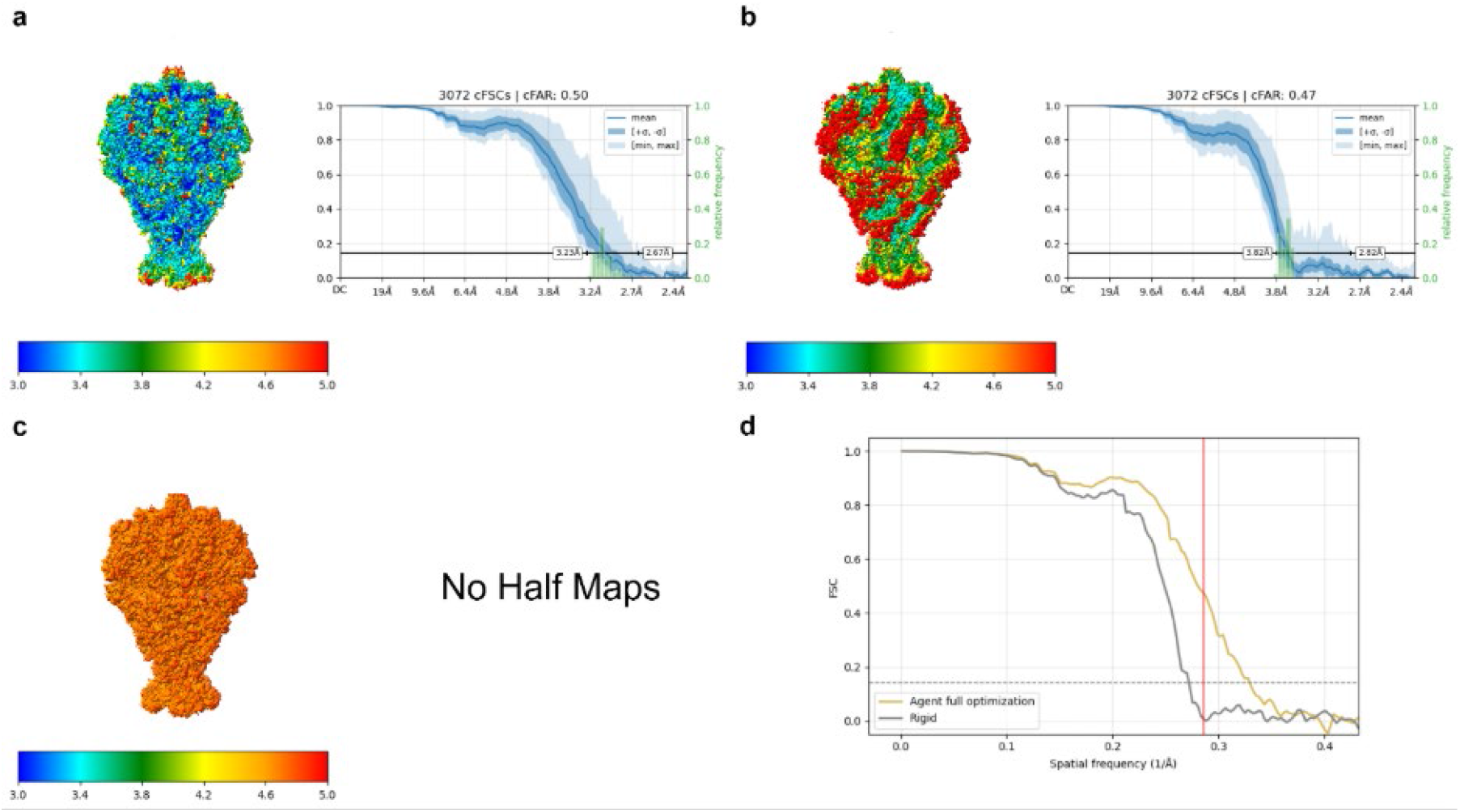
Comparison of reconstructions of TcdA1 produced by different processing workflows on the EMPIAR-10089 dataset(42). a–b, Density maps from the full cryoAgent workflow (a) and conventional rigid workflow (b) shown with corresponding local resolution and cFAR score maps. c, EMD-3645 density map. As the half-maps were not available, local resolution was estimated using ResMap in single-map mode, and cFAR was not evaluated. d, FSC curves comparing reconstructions obtained from the full cryoAgent workflow, the conventional rigid automated workflow. The red line indicates the deposited resolution for the corresponding EMDB density.

**Figure S21.**
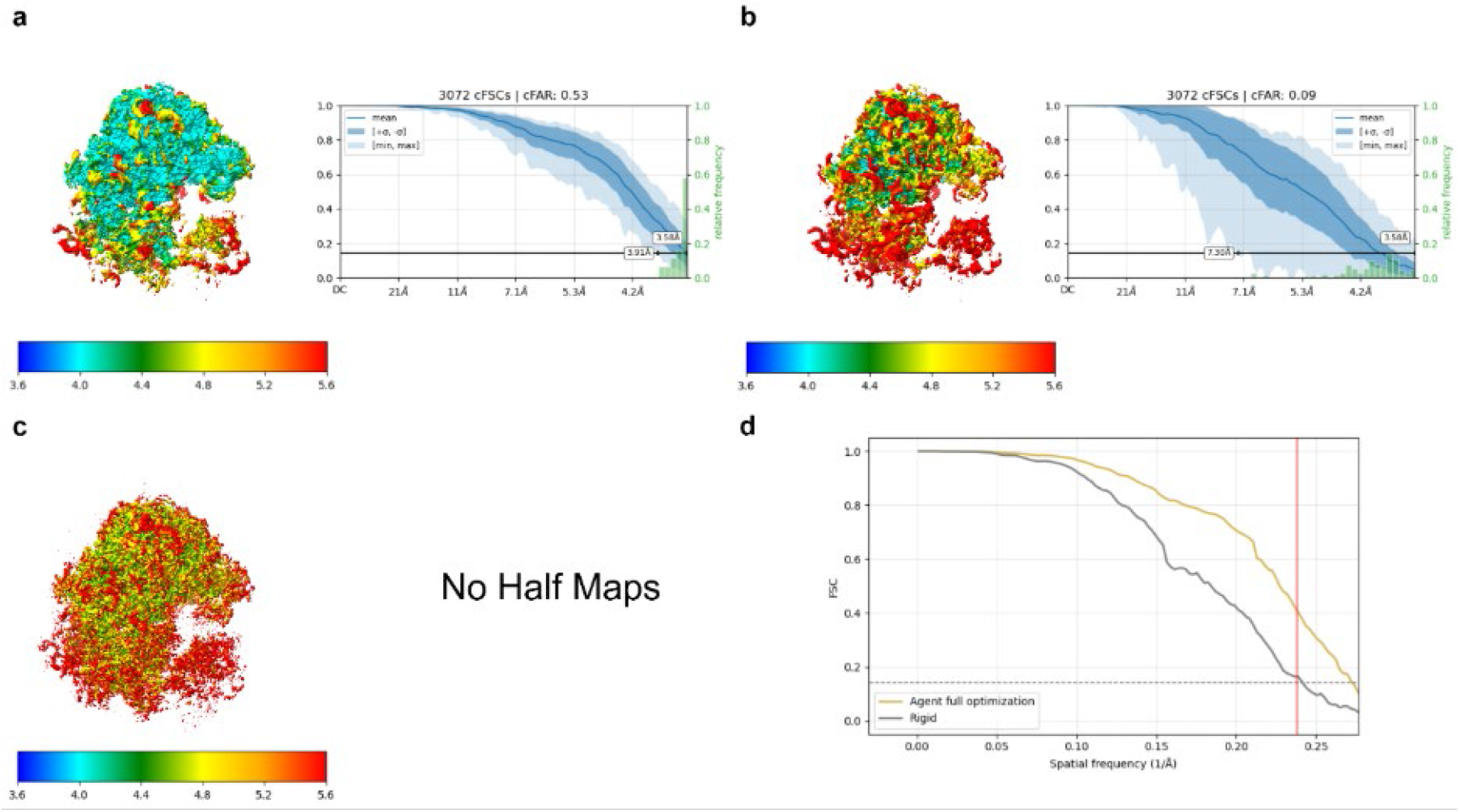
| Comparison of reconstructions of S.cereviseae 80S ribosome produced by different processing workflows on thes EMPIAR-10002 dataset(43). a–b, Density maps from the full cryoAgent workflow (a) and conventional rigid workflow (b) shown with corresponding local resolution and cFAR score maps. c, EMD-2275 density map. As the half-maps were not available, local resolution was estimated using ResMap in single-map mode, and cFAR was not evaluated. d, FSC curves comparing reconstructions obtained from the full cryoAgent workflow, the conventional rigid automated workflow. The red line indicates the deposited resolution for the corresponding EMDB density.

**Figure S22.**
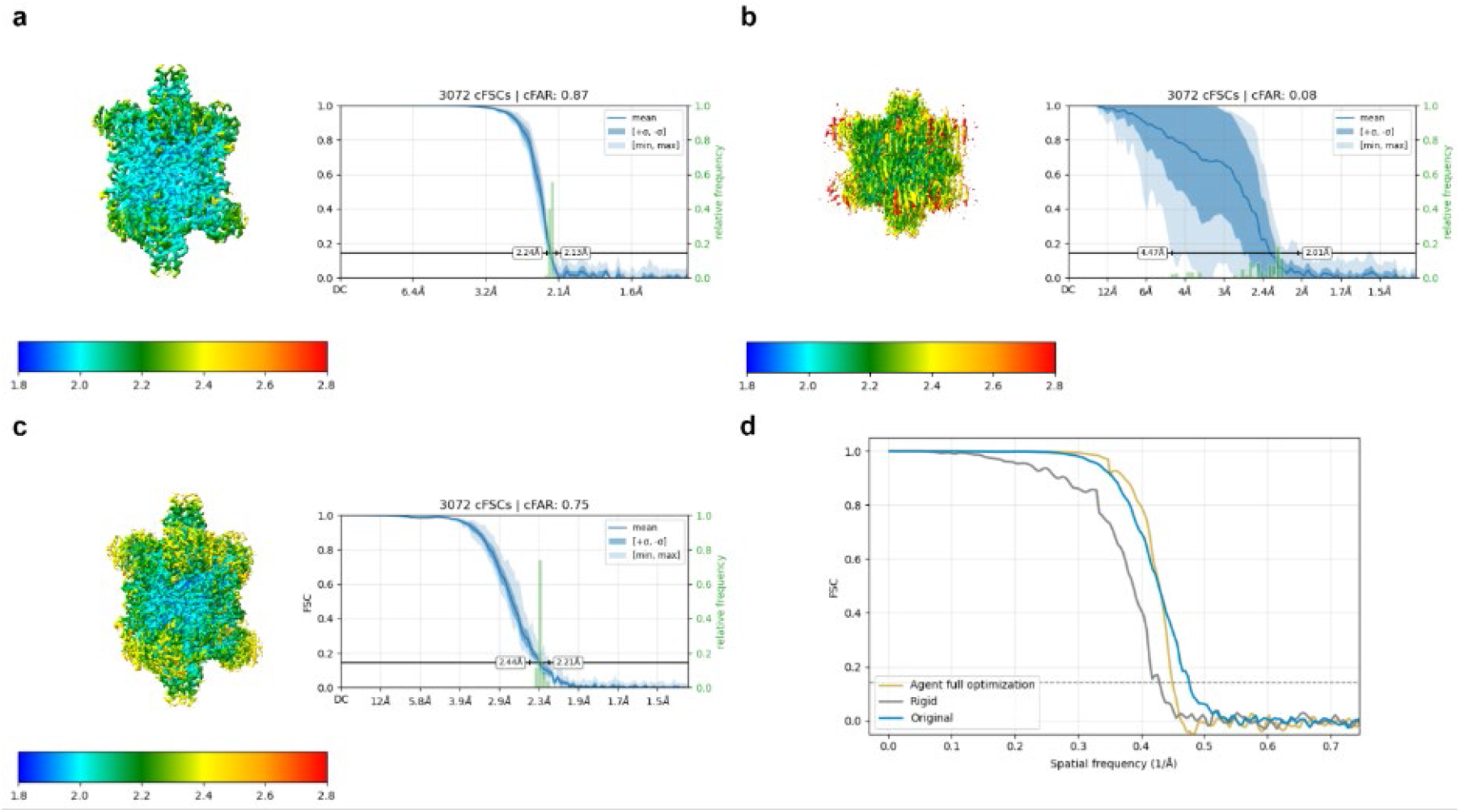
Comparison of reconstructions of bovine liver glutamate dehydrogenase produced by different processing workflows on the EMPIAR-10217 dataset(37) a–c, Density maps from the full cryoAgent workflow (a), conventional rigid workflow (b), and EMD-9203 (c), shown with corresponding local resolution and cFAR score maps. d, FSC curves comparing reconstructions obtained from the full cryoAgent workflow, the conventional rigid automated workflow, and the corresponding EMDB density.

**Figure S23.**
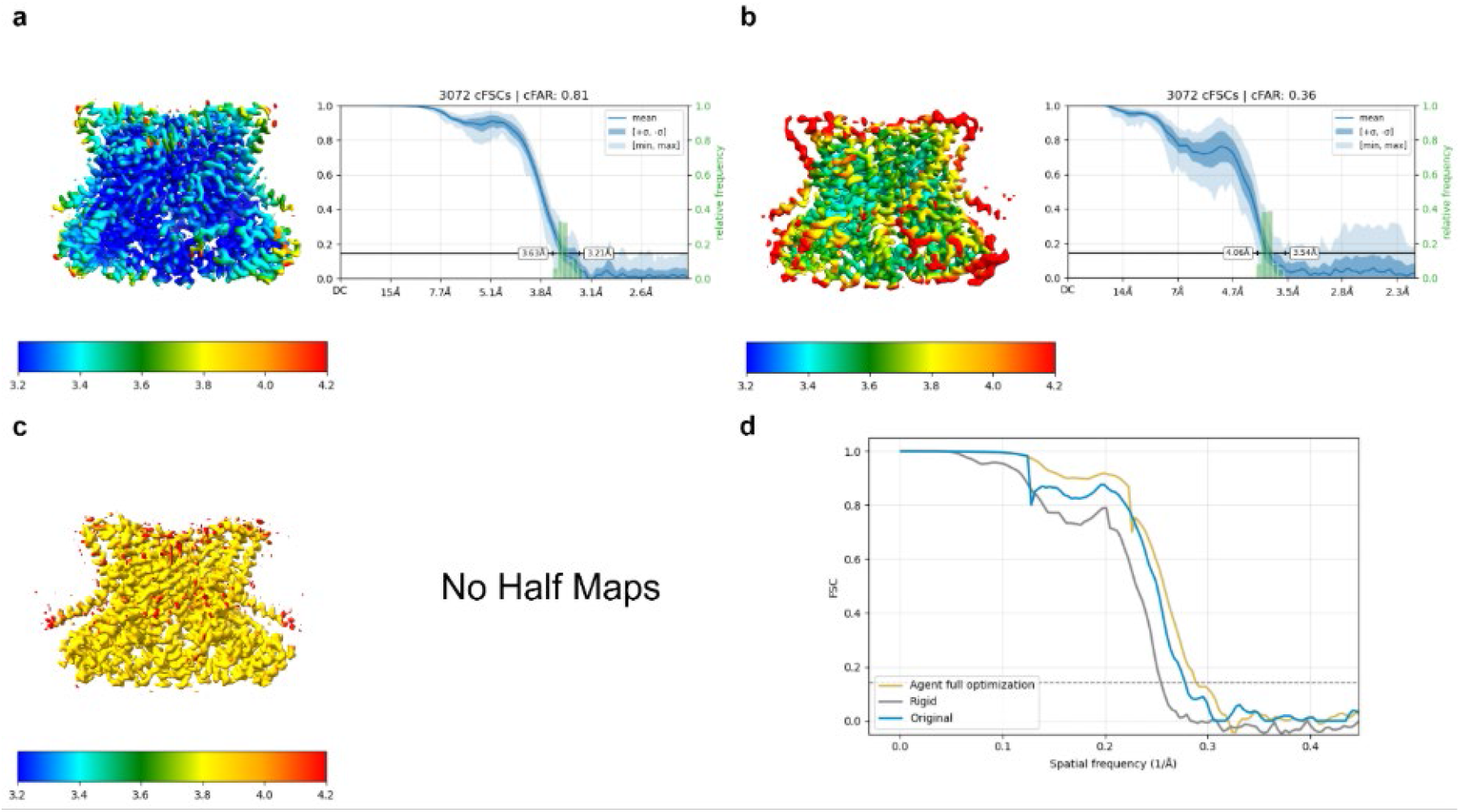
Comparison of reconstructions of afTMEM16 produced by different processing workflows on the EMPIAR-10240 dataset(44). a–b, Density maps from the full cryoAgent workflow (a) and conventional rigid workflow (b) shown with corresponding local resolution and cFAR score maps. c, EMD-8959 density map. As the half-maps were not available, local resolution was estimated using ResMap in single-map mode, and cFAR was not evaluated. d, FSC curves comparing reconstructions obtained from the full cryoAgent workflow, the conventional rigid automated workflow, and the corresponding EMDB density.

**Figure S24.**
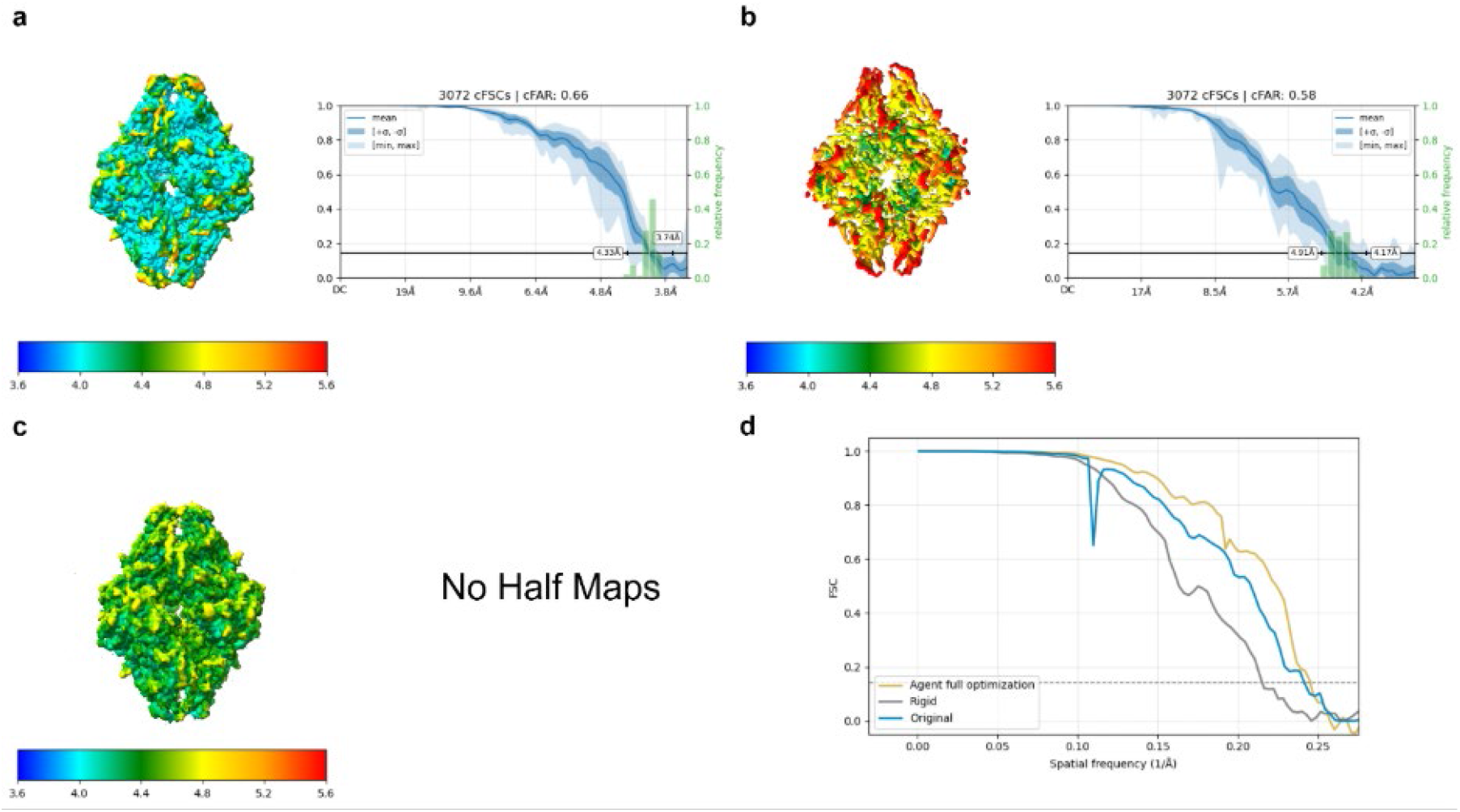
Comparison of reconstructions produced by different processing workflows on the EMPIAR-10017 dataset(45). a–b, Density maps from the full cryoAgent workflow (a) and conventional rigid workflow (b) shown with corresponding local resolution and cFAR score maps. c, EMD-2824 density map. As the half-maps were not available, local resolution was estimated using ResMap in single-map mode, and cFAR was not evaluated. d, FSC curves comparing reconstructions obtained from the full cryoAgent workflow, the conventional rigid automated workflow, and the corresponding EMDB density.

**Figure S25.**
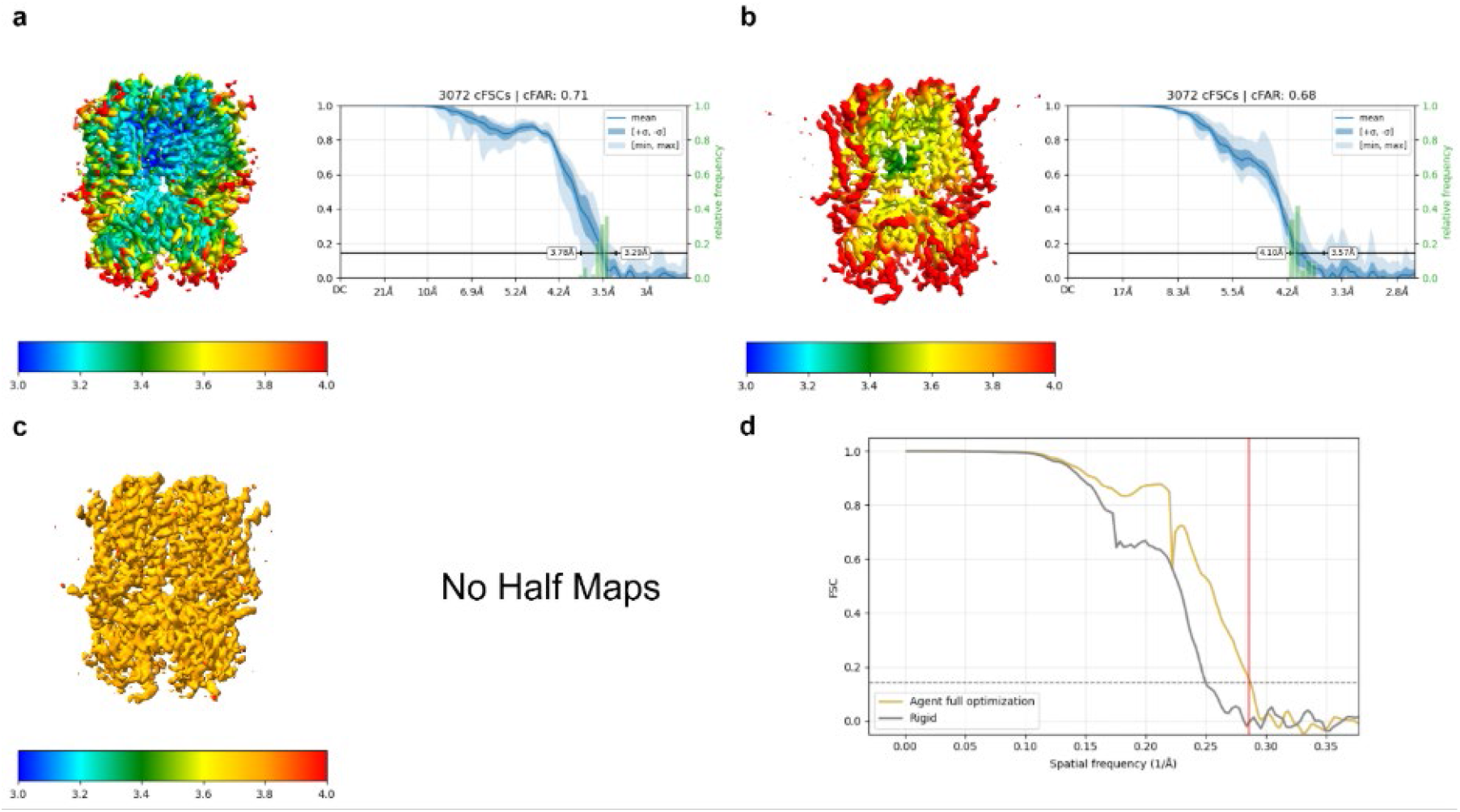
Comparison of reconstructions of Human HCN1 hyperpolarization-activated cyclic nucleotide-gated ion channel produced by different processing workflows on the EMPIAR-10081 dataset(46). a–b, Density maps from the full cryoAgent workflow (a) and conventional rigid workflow (b) shown with corresponding local resolution and cFAR score maps. c, EMD-8511 density map. As the half-maps were not available, local resolution was estimated using ResMap in single-map mode, and cFAR was not evaluated. d, FSC curves comparing reconstructions obtained from the full cryoAgent workflow, the conventional rigid automated workflow. The red line indicates the deposited resolution for the corresponding EMDB density.

**Figure S26.**
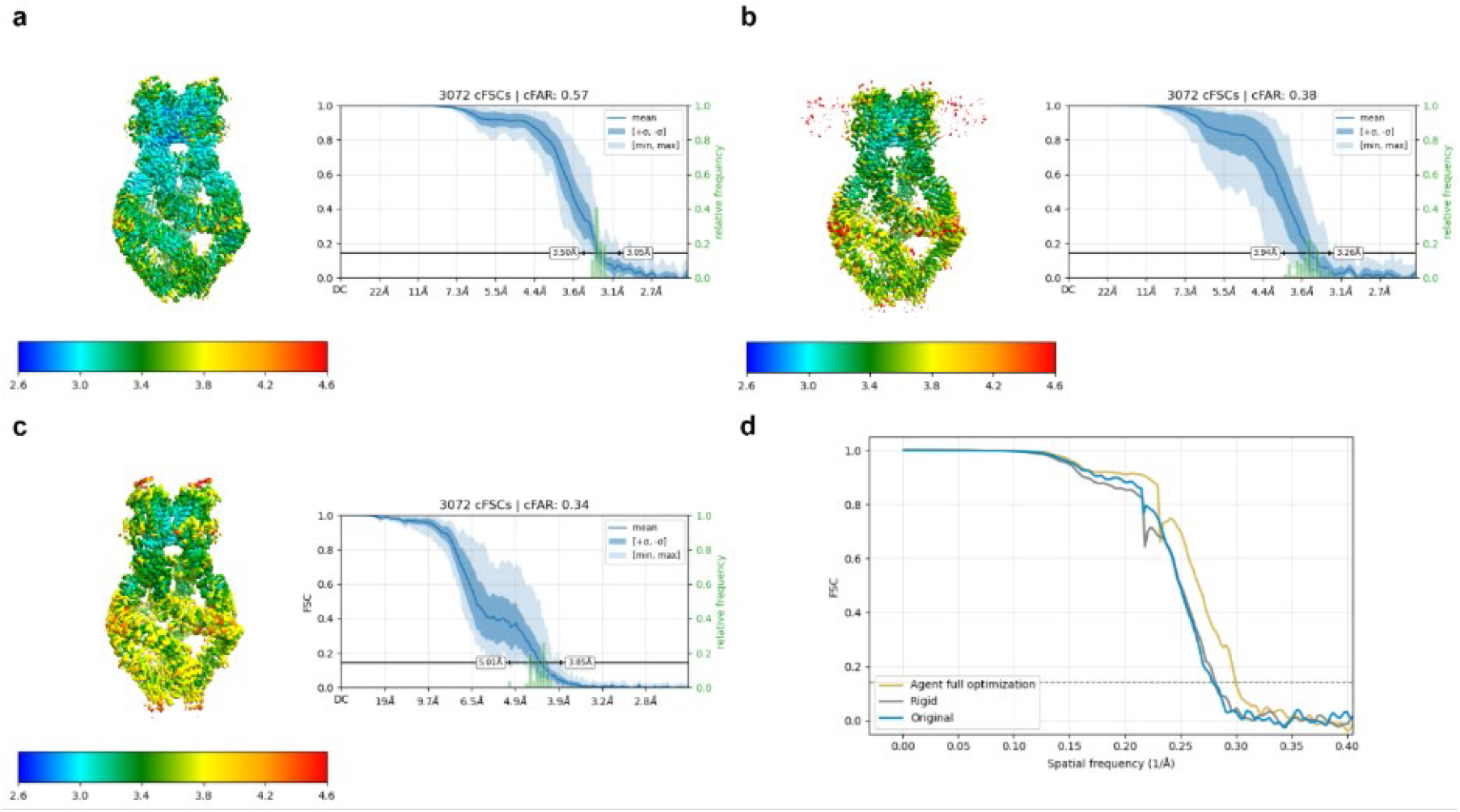
Comparison of reconstructions of Mechanotransduction channel NOMPC produced by different processing workflows on the EMPIAR-10093 dataset(47). a–c, Density maps from the full cryoAgent workflow (a), conventional rigid workflow (b), and EMD-8702 (c), shown with corresponding local resolution and cFAR score maps. d, FSC curves comparing reconstructions obtained from the full cryoAgent workflow, the conventional rigid automated workflow, and the corresponding EMDB density.

**Figure S27.**
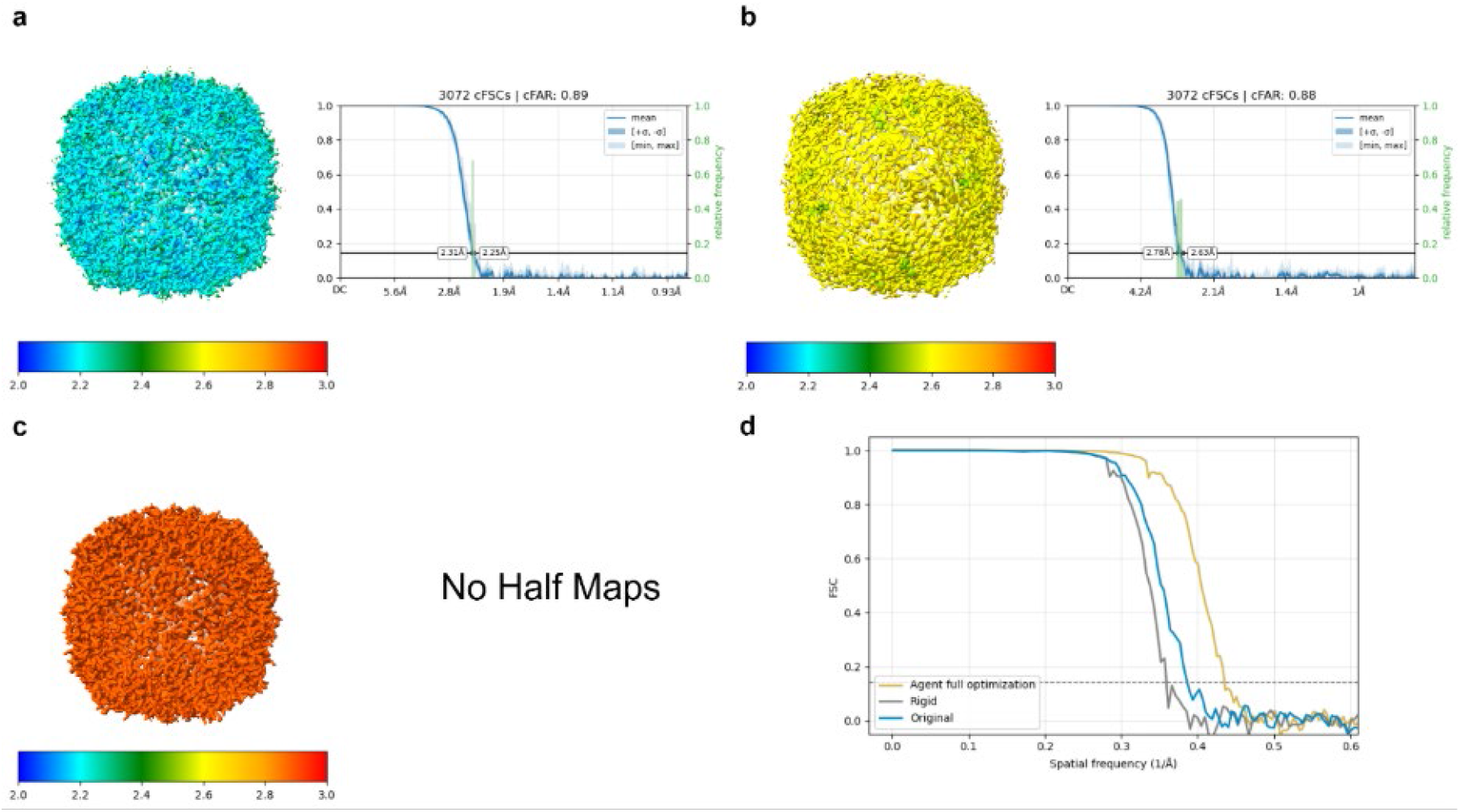
Comparison of reconstructions of apoferritin produced by different processing workflows on the EMPIAR-10405 dataset(48). a–b, Density maps from the full cryoAgent workflow (a) and conventional rigid workflow (b) shown with corresponding local resolution and cFAR score maps. c, EMD-30083 density map. As the half-maps were not available, local resolution was estimated using ResMap in single-map mode, and cFAR was not evaluated. d, FSC curves comparing reconstructions obtained from the full cryoAgent workflow, the conventional rigid automated workflow, and the corresponding EMDB density.

**Figure S28.**
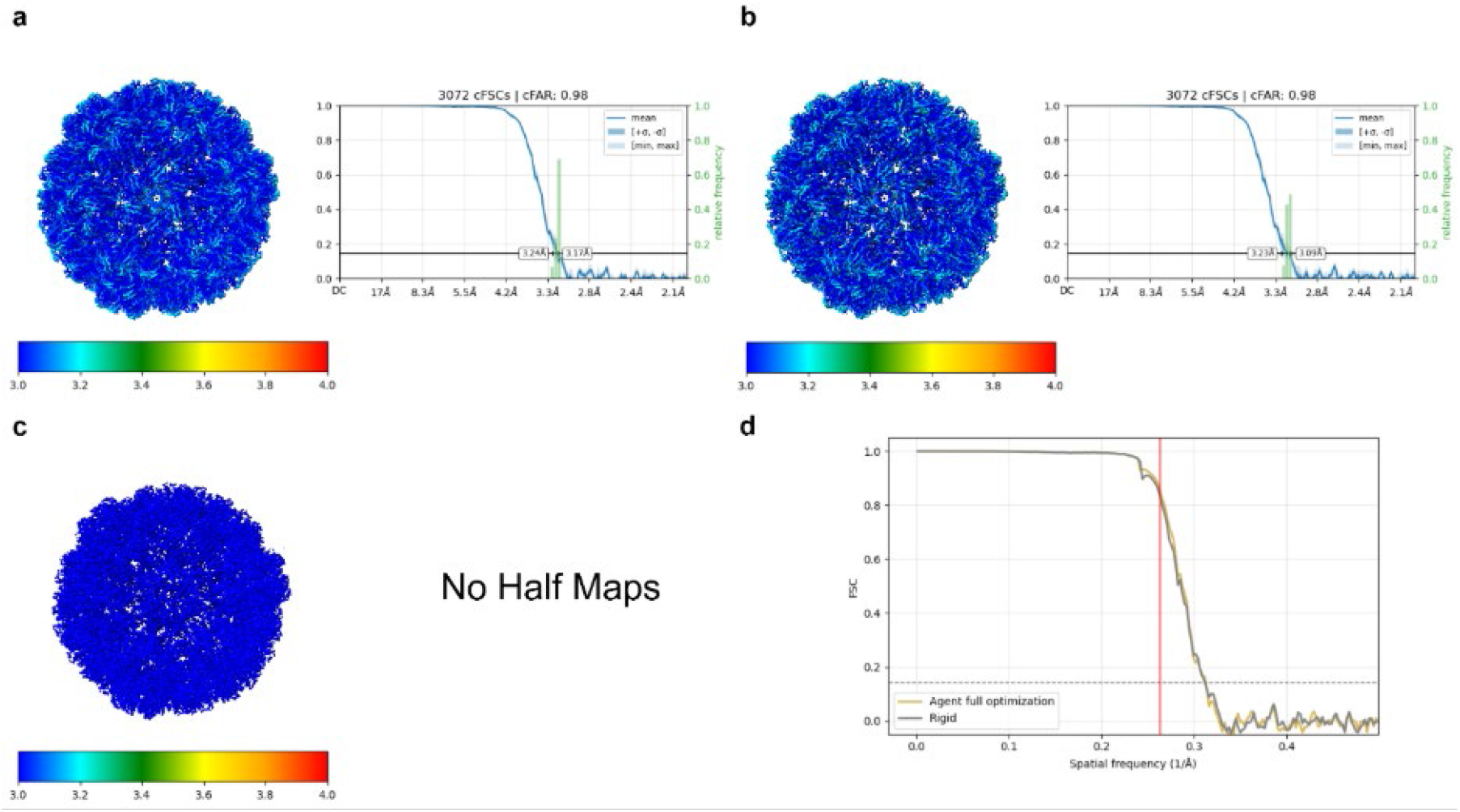
Comparison of reconstructions of brome mosaic virusss produced by different processing workflows on the EMPIAR-10010 dataset(49). a–b, Density maps from the full cryoAgent workflow (a) and conventional rigid workflow (b) shown with corresponding local resolution and cFAR score maps. c, EMD-6000 density map. As the half-maps were not available, local resolution was estimated using ResMap in single-map mode, and cFAR was not evaluated. d, FSC curves comparing reconstructions obtained from the full cryoAgent workflow, the conventional rigid automated workflow. The red line indicates the deposited resolution for the corresponding EMDB density.

**Figure S29.**
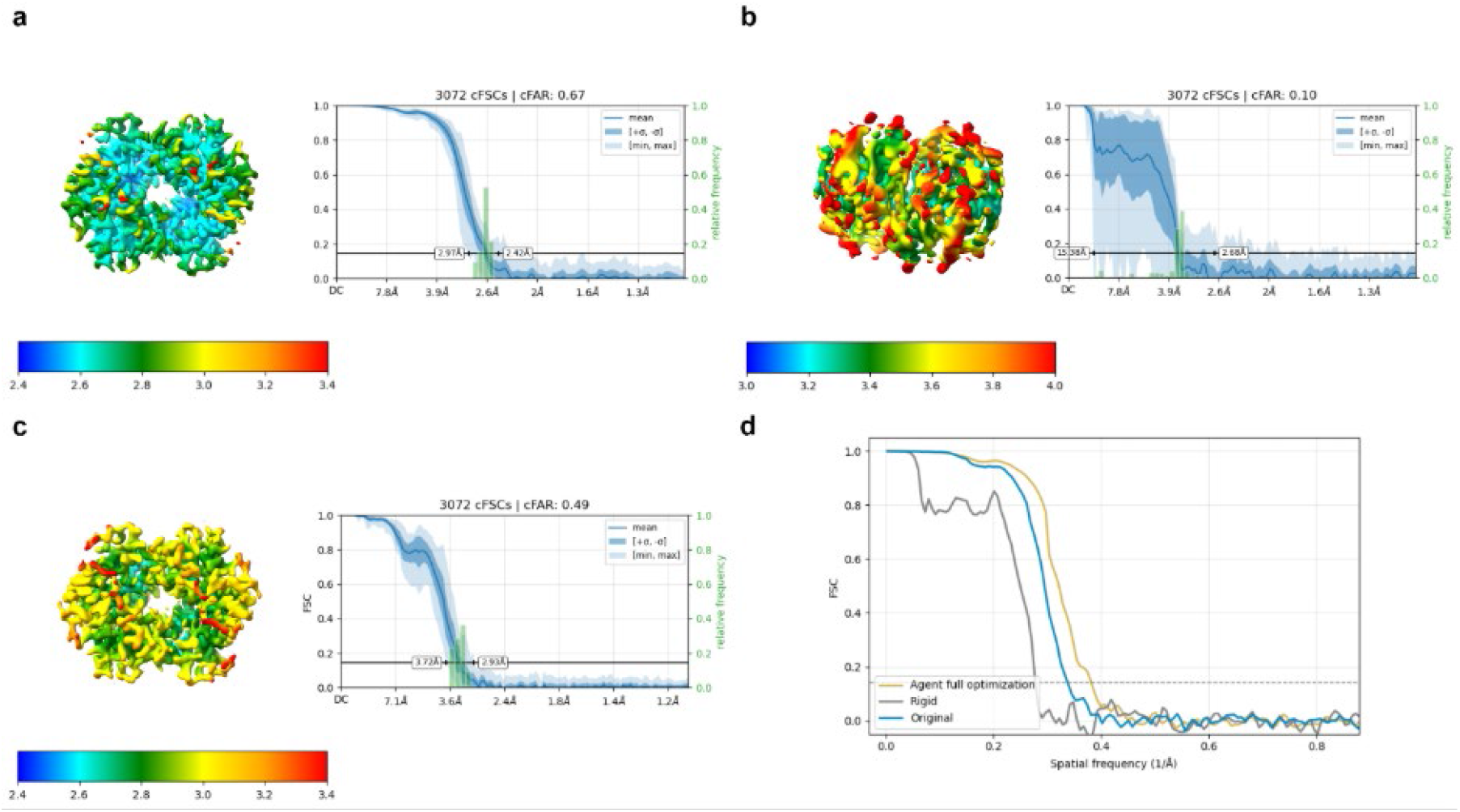
Comparison of reconstructions of human methemoglobin produced by different processing workflows on the EMPIAR-10250 dataset(50). a–c, Density maps from the full cryoAgent workflow (a), conventional rigid workflow (b), and EMD-0407 (c), shown with corresponding local resolution and cFAR score maps. d, FSC curves comparing reconstructions obtained from the full cryoAgent workflow, the conventional rigid automated workflow, and the corresponding EMDB density.

**Figure S30.**
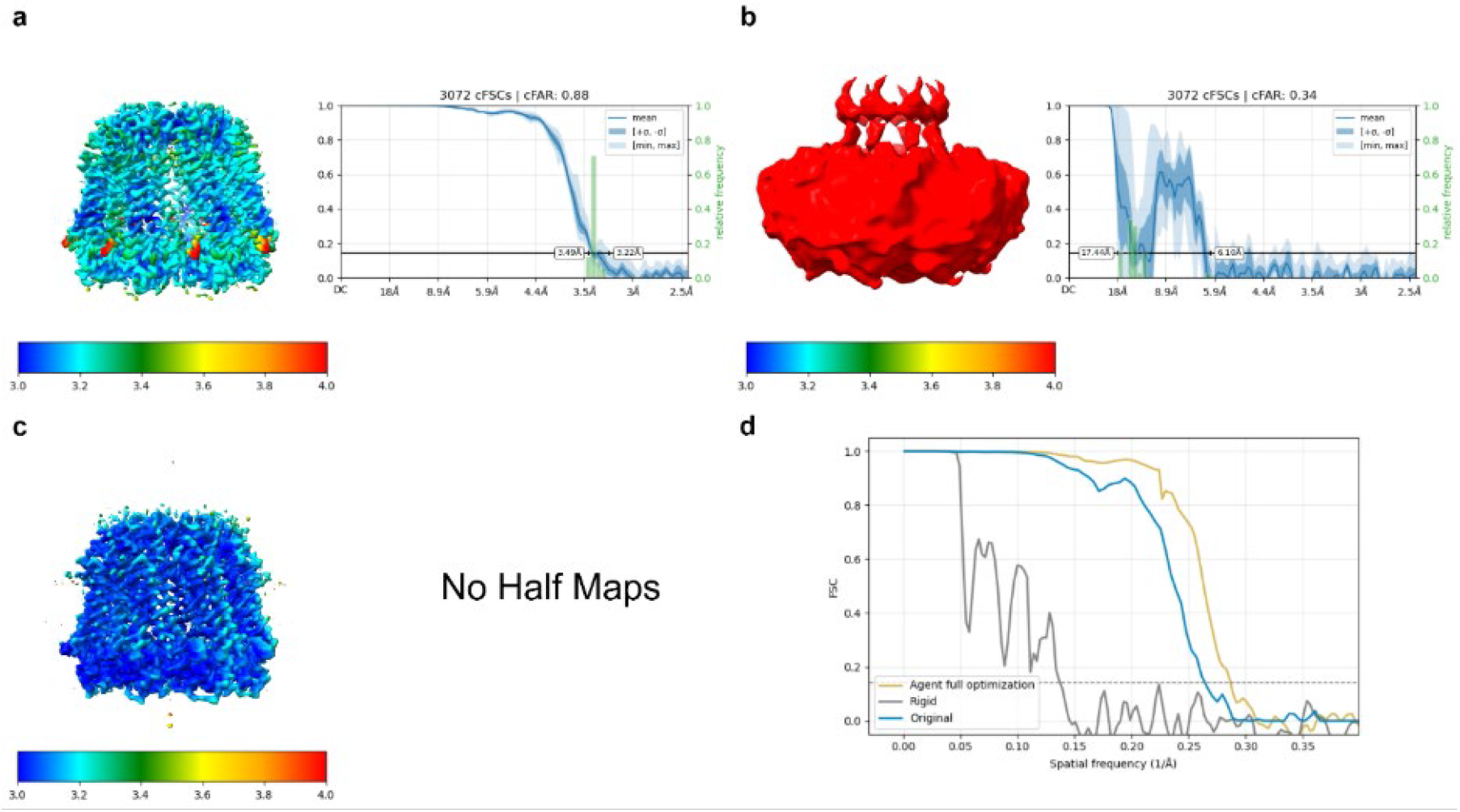
Comparison of reconstructions of octameric complex of innexin-6 produced by different processing workflows on the EMPIAR-10289 dataset(51). a–b, Density maps from the full cryoAgent workflow (a) and conventional rigid workflow (b), shown with corresponding local resolution and cFAR score maps. c, EMD-9971 density map. As the half-maps were not available, local resolution was estimated using ResMap in single-map mode, and cFAR was not evaluated. d, FSC curves comparing reconstructions obtained from the full cryoAgent workflow, the conventional rigid automated workflow, and the corresponding EMDB density.

**Figure S31.**
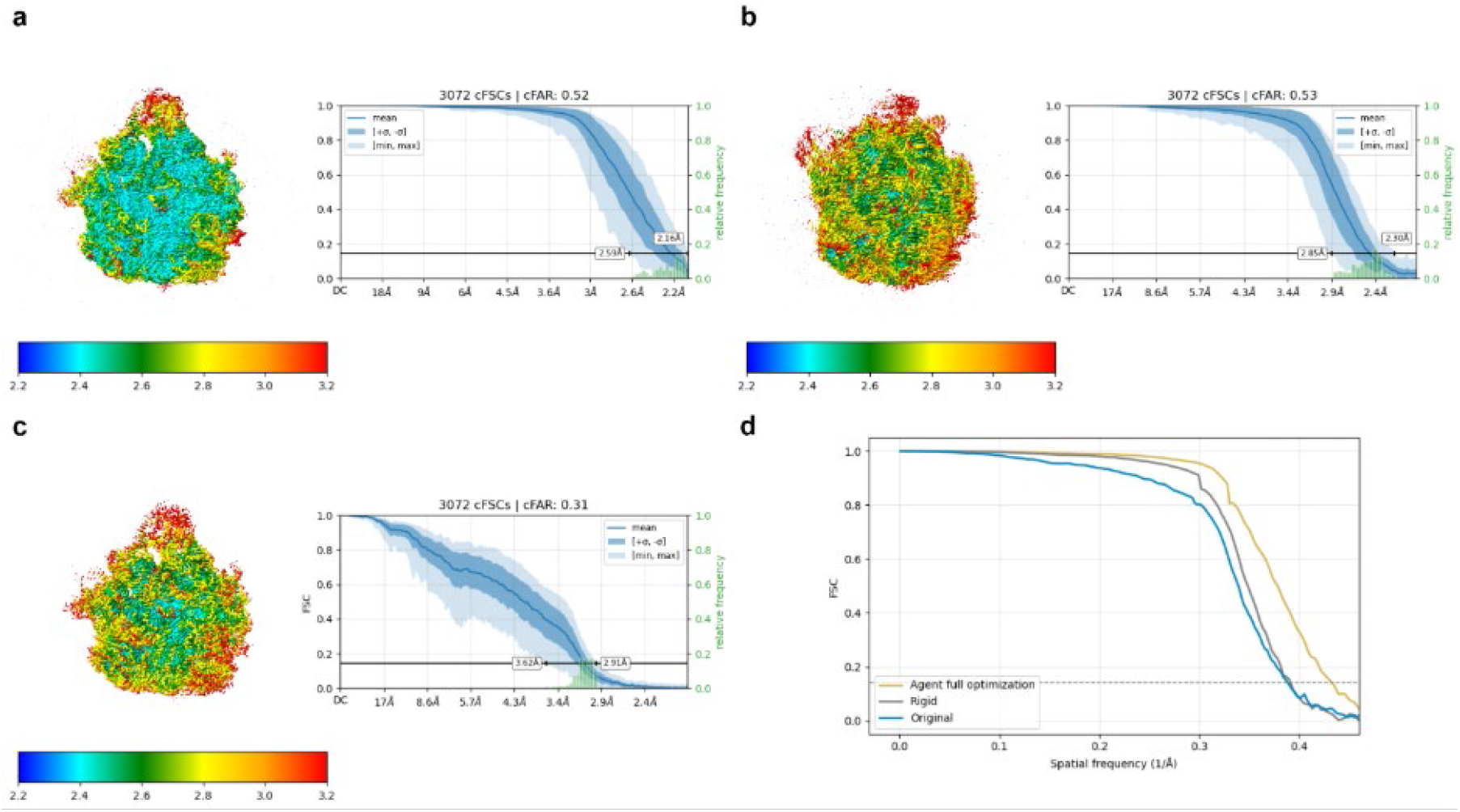
Comparison of reconstructions of acinetobacter baumannii ribosome-amikacin complex - 50S subunit produced by different processing workflows on the EMPIAR-10406 dataset(52). a–c, Density maps from the full cryoAgent workflow (a), conventional rigid workflow (b), and EMD-10809 (c), shown with corresponding local resolution and cFAR score maps. d, FSC curves comparing reconstructions obtained from the full cryoAgent workflow, the conventional rigid automated workflow, and the corresponding EMDB density.

**Figure S32.**
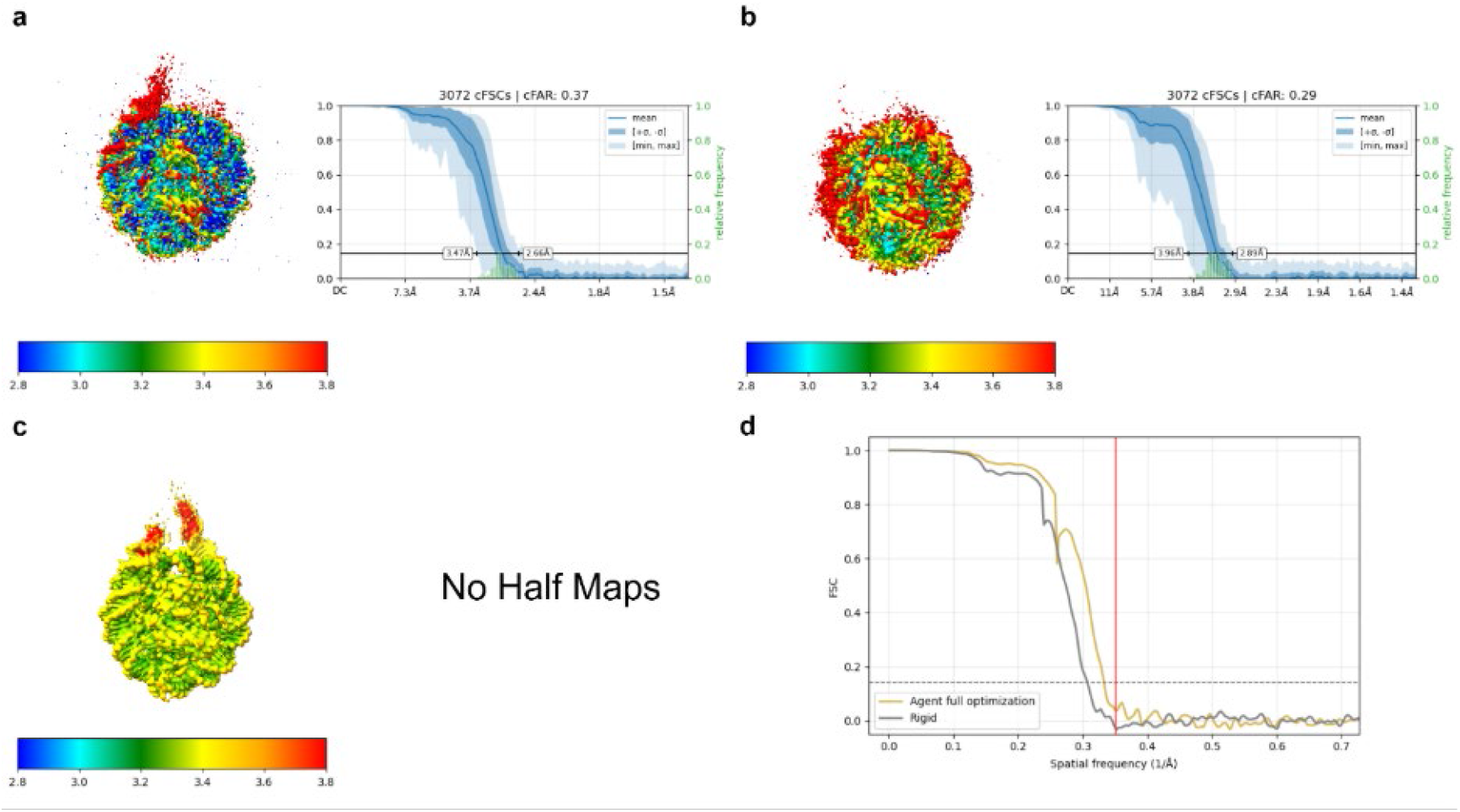
Comparison of reconstructions of 197bp nucleosome produced by different processing workflows on the EMPIAR-10576 dataset(53). a–b, Density maps from the full cryoAgent workflow (a) and conventional rigid workflow (b) shown with corresponding local resolution and cFAR score maps. c, EMD-22686 density map. As the half-maps were not available, local resolution was estimated using ResMap in single-map mode, and cFAR was not evaluated. d, FSC curves comparing reconstructions obtained from the full cryoAgent workflow, the conventional rigid automated workflow. The red line indicates the deposited resolution for the corresponding EMDB density.

**Figure S33.**
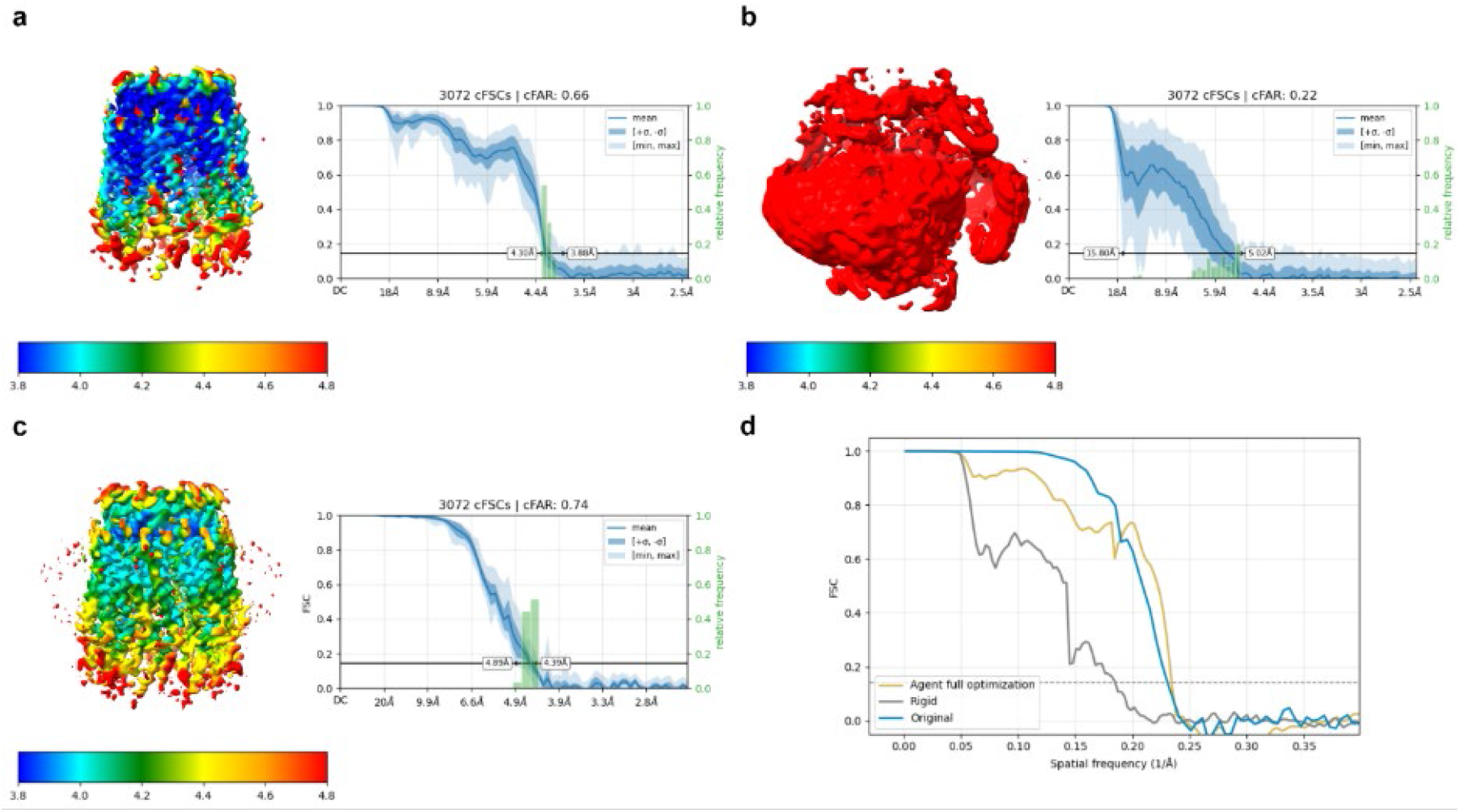
Comparison of reconstructions of human pannexin-1 produced by different processing workflows on the EMPIAR-10760 dataset(54). a–c, Density maps from the full cryoAgent workflow (a), conventional rigid workflow (b), and EMD-32768 (c), shown with corresponding local resolution and cFAR score maps. d, FSC curves comparing reconstructions obtained from the full cryoAgent workflow, the conventional rigid automated workflow, and the corresponding EMDB density.

**Figure S34.**
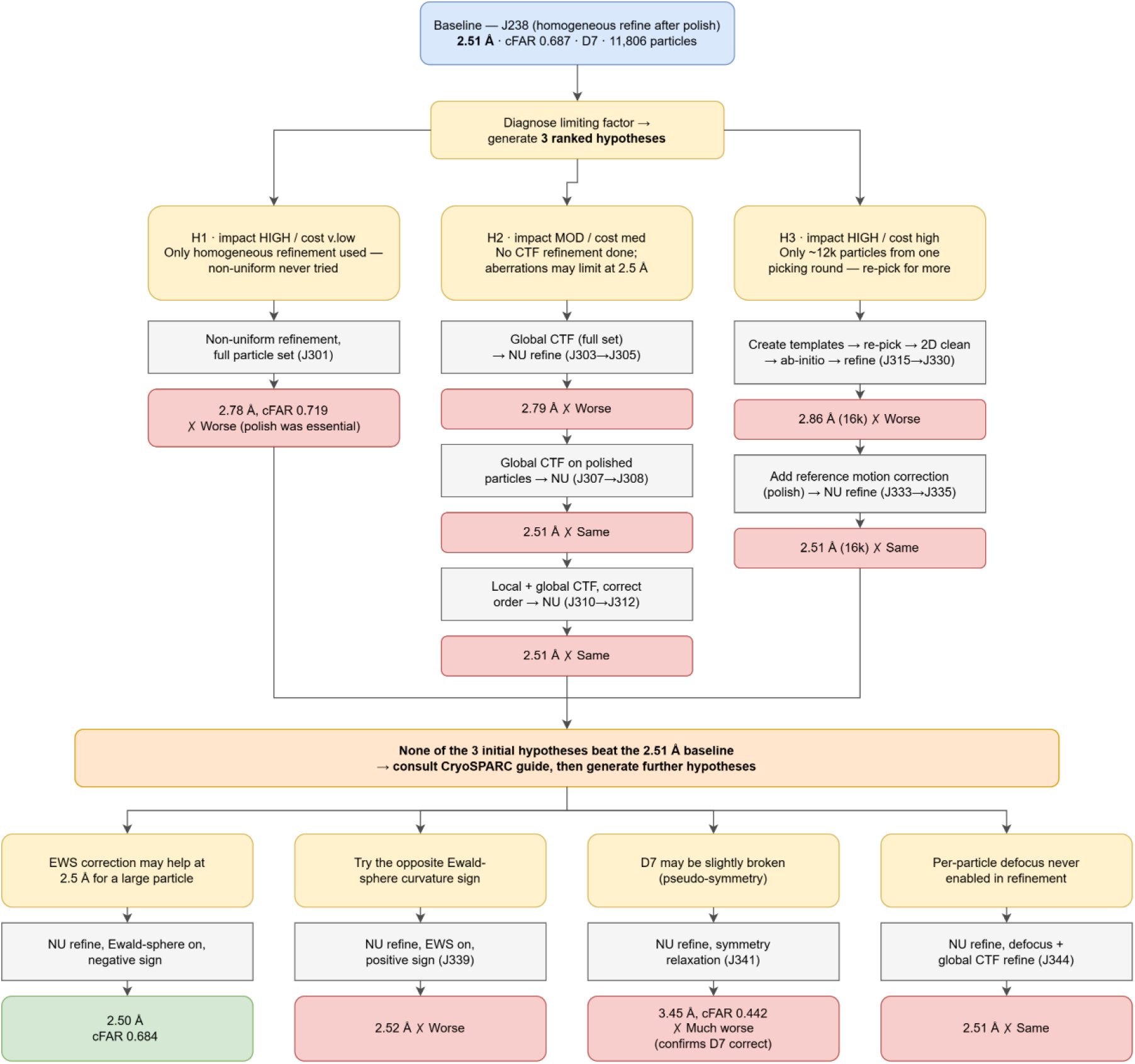
Two hypothesis-driven exploration trials initialized from the guided optimization result on the EMPIAR-10025. The workflow shows the exploration path followed by the hypothesis-driven exploration mode after initialization from the reconstruction generated by guided optimization on the EMPIAR-10025 dataset.

**Figure S35.**
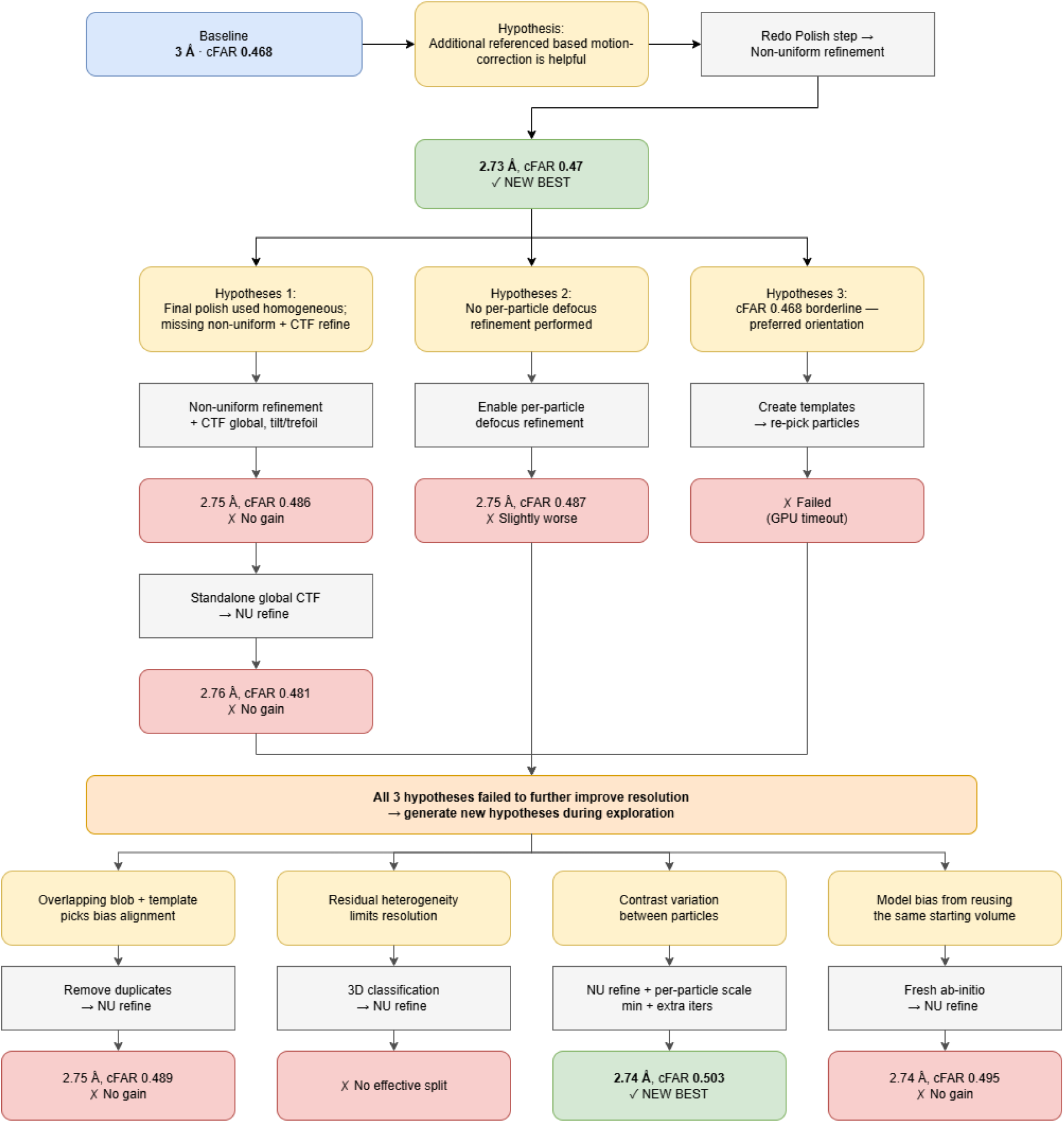
Two hypothesis-driven exploration trials initialized from the guided optimization result on the EMPIAR-10576. The workflow shows the exploration path followed by the hypothesis-driven exploration mode after initialization from the reconstruction generated by guided optimization on the EMPIAR-10576 dataset.

**Figure S36.**
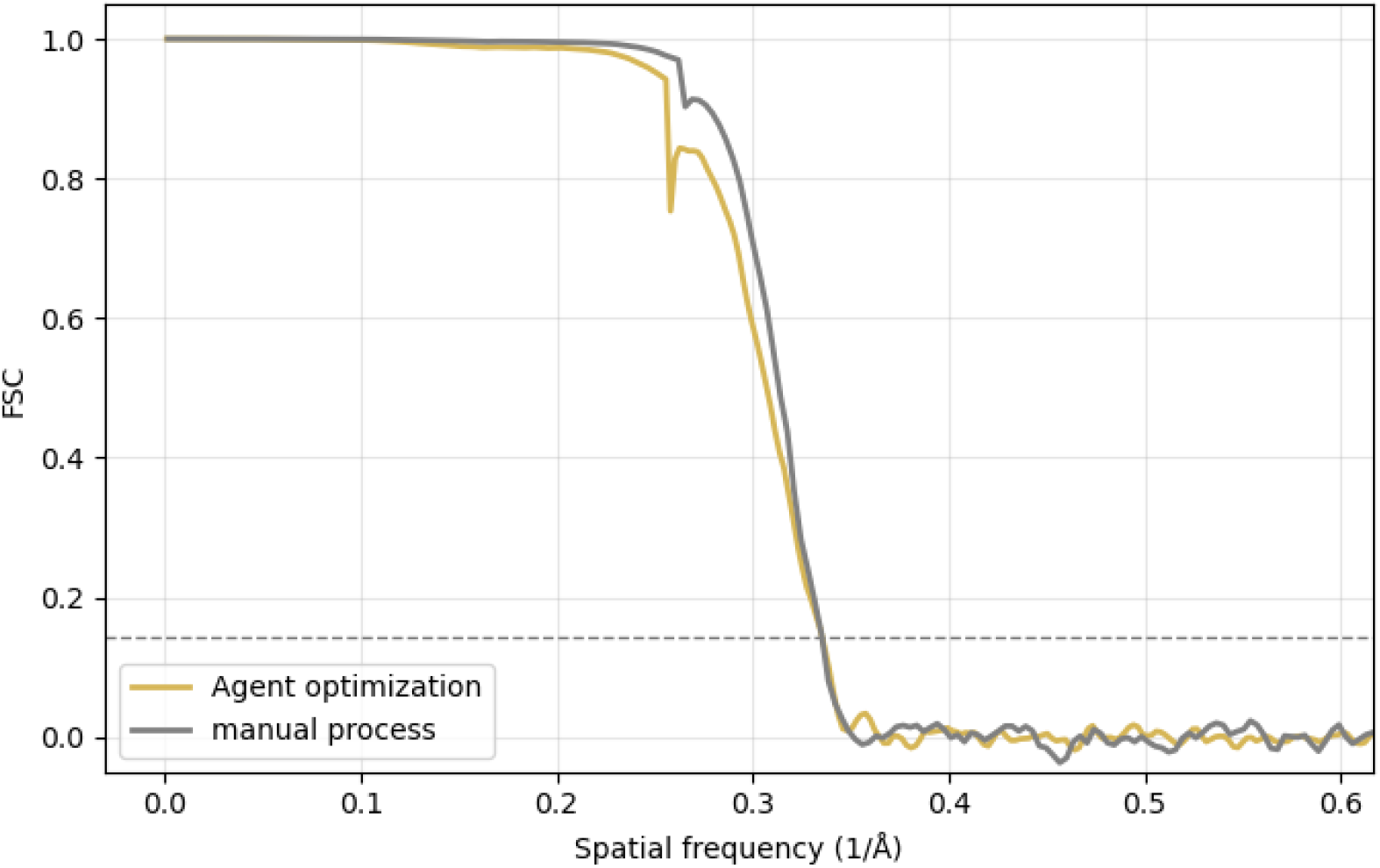
The FSC curve of the density from the guided optimization and manual process.

**Figure S37.**
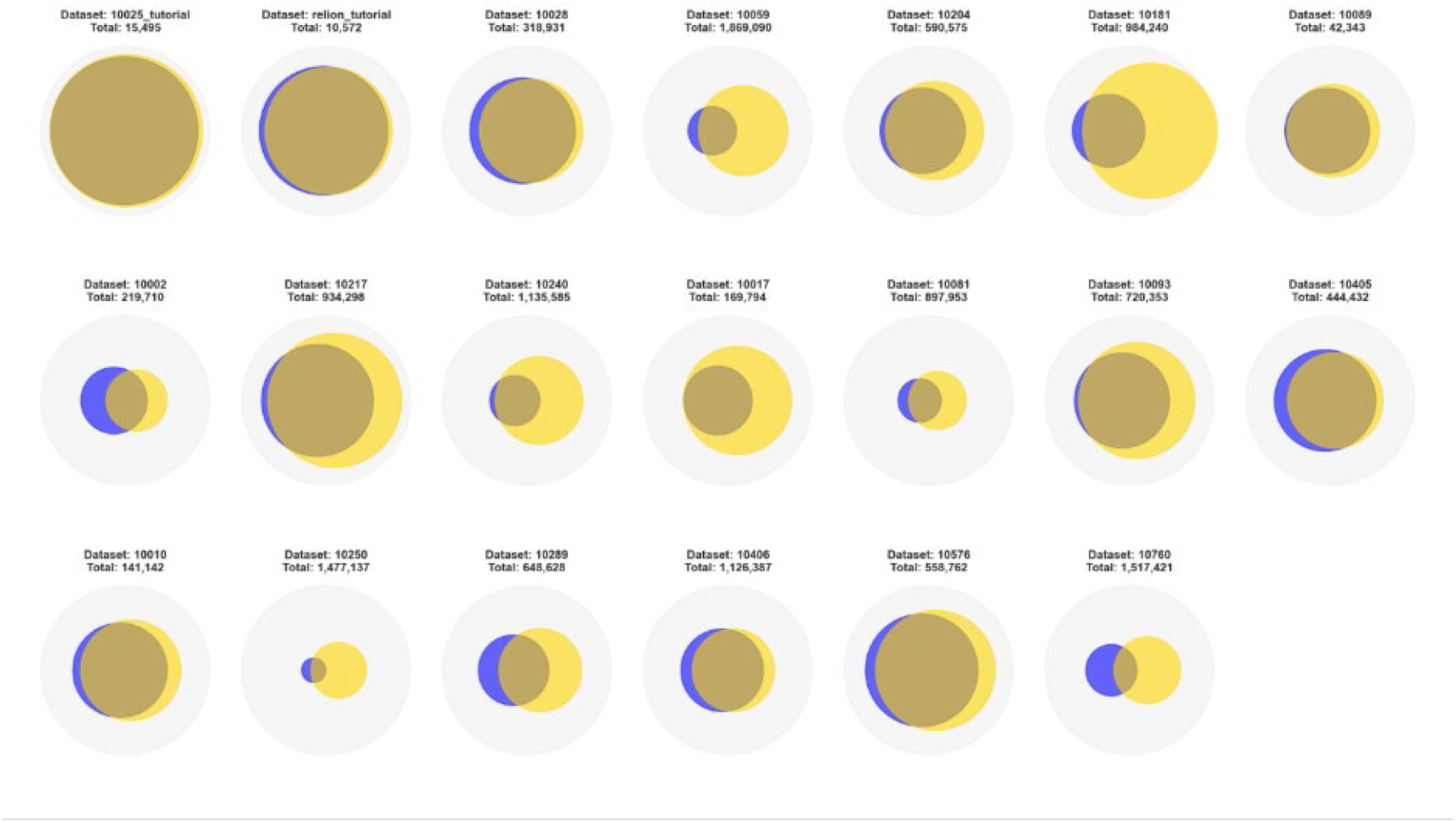
Illustration of the particle set selected by cryoAgent guided optimization. Gray circle: Particle set from the initial template picker. Yellow circle: Particle set from the conventional rigid automated workflow. Blue circle: Particle set selected by cryoAgent.

**Figure S38.**
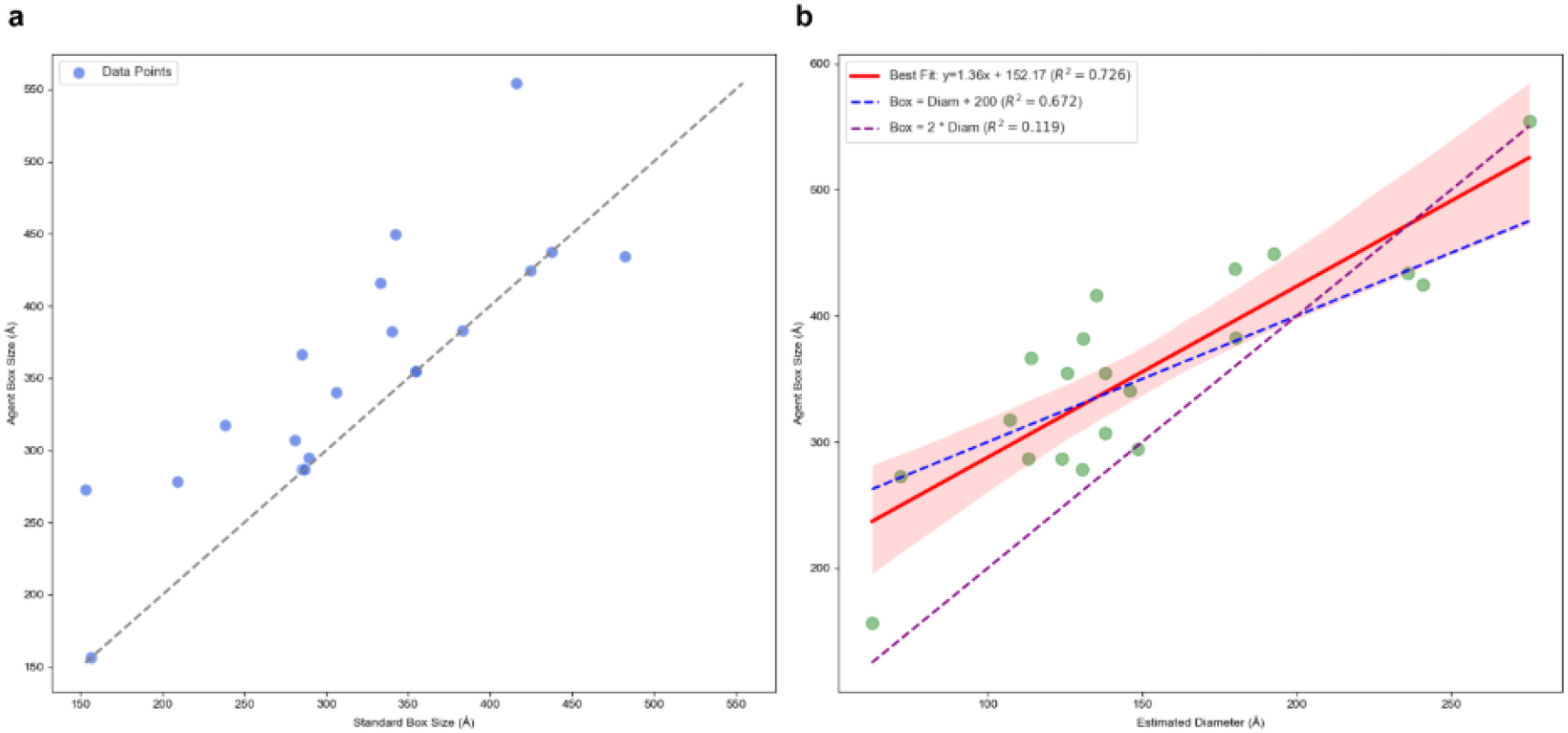
Illustration of the box size estimated by cryoAgent guided optimization. a, Comparison of box size of the original EMDB density and the final box size estimated by cryoAgent. b, Comparison between the particle diameter estimated from the radial profile of the final cryoAgent density and the final box size estimated by cryoAgent.

---

**Table S1.** Summary of cryo-EM datasets statistics and sample properties.

| ID | Optimization<br>System<br>Processed | Heterogen<br>eity<br>System<br>Processed | Protein Type | Strutcutr<br>al Weight<br>(kDa) | Accelerati<br>on<br>Voltage<br>(kV) | Spherical<br>Aberration<br>(mm) | Electron<br>Dose<br>(e/A^2) | Defocu<br>s range<br>(µm) | Microscope | Detector | Total<br>Particle<br>Number |
| --- | --- | --- | --- | --- | --- | --- | --- | --- | --- | --- | --- |
| cryosp<br>arc_tut<br>orial | Yes | Yes | T20S proteasome | 659 | 300 | 2.7 | 53 | 0.9 -<br>2.4 | FEI TITAN<br>KRIOS | GATAN K2 (4k x 4k) | 15495 |
| relion_<br>tutorial | Yes | Yes | β -galactosidase | 466.41 | 200 | 1.4 |  | 0.3 - 2 | JEOL CRYO<br>ARM 200 | GATAN K2 IS (4k x<br>4k) | 10572 |
| 10002 | Yes | No | S.cereviseae 80S<br>ribosome | 6,757.34 | 300 | 2 | 16 | 1.191 -<br>3.44 | FEI POLARA<br>300 | FEI FALCON I (4k x<br>4k) | 219710 |
| 10010 | Yes | No | brome mosaic<br>virus | 3673.8 | 300 | 5.6 |  | 0.5 -<br>2.0 | JEOL<br>3200FSC | DIRECT<br>ELECTRON DE-12<br>(4k x 3k) | 141142 |
| 10017 | Yes | No | β -galactosidase | 466.41 | 300 | 2.7 | 24 | 3 | FEI POLARA<br>300 | FEI FALCON II<br>(CCD, 4k x 4k) | 169794 |
| 10028 | Yes | Yes | Ribosome (80S) | 2,135.89 | 300 | 2.7 | 20 | 0.8 to<br>3.8 | Titan Krios<br>Microscope<br>(FEI) | FEI Falcon II<br>Detector | 318931 |
| 10059 | Yes | No | TRPV1 | 317.88 | 300 | 2 | 41 | 0.7 -<br>2.2 | FEI POLARA<br>300 | K2 Summit direct<br>electron detector<br>camera (Gatan) | 1869090 |
| 10076 | No | Yes | bL17-depleted<br>large ribosomal<br>subunit | 4,270.48 | 300 | 2.7 | 34.27 | 1.0 -<br>3.5 | FEI TITAN<br>KRIOS | GATAN K2<br>SUMMIT (4k x 4k) | 131899 |
| 10081 | Yes | No | Human HCN1<br>hyperpolarization-activated cyclic<br>nucleotide-gated<br>ion channel | 298.5 | 300 | 2.7 | 1.26 | 1.5 -<br>3.3 | FEI TITAN<br>KRIOS | GATAN K2<br>SUMMIT (4k x 4k) | 897953 |
| 10089 | Yes | Yes | TcdA1 | 1,416.15 | 300 | 0 | 2.5 |  | FEI TITAN<br>KRIOS | FEI FALCON II (4k<br>x 4k) | 42343 |
| 10093 | Yes | No | Mechanotransduct<br>ion channel<br>NOMPC | 779.4 | 300 | 2.7 | 54 | 1.5 -<br>3.0 | FEI POLARA<br>300 | GATAN K2<br>SUMMIT (4k x 4k) | 720353 |
| 10181 | Yes | No | Rabbit Muscle<br>Aldolase | 157 | 200 | 2.7 | 68 | 0.6 -<br>2.7 | FEI TALOS<br>ARCTICA | GATAN K2<br>SUMMIT (4k x 4k) | 984240 |
| 10204 | Yes | No | $\beta$ -galactosidase | 466.41 | 200 | 1.4 | 70 | 0.3 to 2 | JEOL CRYO<br>ARM 200 | GATAN K2 IS (4k x<br>4k) | 590575 |
| 10217 | Yes | No | bovine liver<br>glutamate<br>dehydrogenase | 347 | 300 | 2.7 | 61.25 | 1.0 -<br>3.0 | FEI TITAN<br>KRIOS | GATAN K2<br>SUMMIT (4k x 4k) | 934298 |
| 10240 | Yes | No | afTMEM16 | 171.72 | 300 | 2.7 | 62.61 | 1.5 to<br>2.5 | Titan Krios Microscope |  | 1135585 |
| 10250 | Yes | No | human<br>methemoglobin | 64 | 200 | 2.7 | 69 | 5.0 -<br>16.0 | FEI TALOS<br>ARCTICA | GATAN K2<br>SUMMIT (4k x 4k) | 1477137 |
| 10289 | Yes | No | octameric<br>complex of<br>innexin-6 | 361.39 | 300 | 2.7 | 35 |  | JEM-3000SFF<br>(JEOL)<br>Microscope | K2 Summit Direct<br>Electron Detector<br>(Gatan) | 648628 |
| 10405 | Yes | No | Apoferitin | 440 | 300 | 2.7 | 60 |  | FEI TITAN<br>KRIOS | GATAN K2<br>SUMMIT (4k x 4k) | 444432 |
| 10406 | Yes | No | *acinetobacter*<br>*baumannii*<br>ribosome-amikacin complex<br>- 50S subunit | 632.89 | 300 | 2.7 | 58 | 0.8 -<br>2.7 |  | Gatan K2 Summit<br>detector | 1126387 |
| 10576 | Yes | No | Nuclear Protein<br>(DNA) | 290.21 | 300 | 2.7 | 43 |  | Titan Krios<br>Microscope | K2 Summit Direct<br>Electron Detector<br>(Gatan) | 558762 |
| 10760 | Yes | No | human pannexin-1 | 321.69 | 300 | 2.7 | 56 | 1.4-3.5 | JEM-3000SFF<br>(JEOL)<br>Microscope | K2 Summit Direct<br>Electron Detector<br>(Gatan) | 1517421 |

**Table S2.**
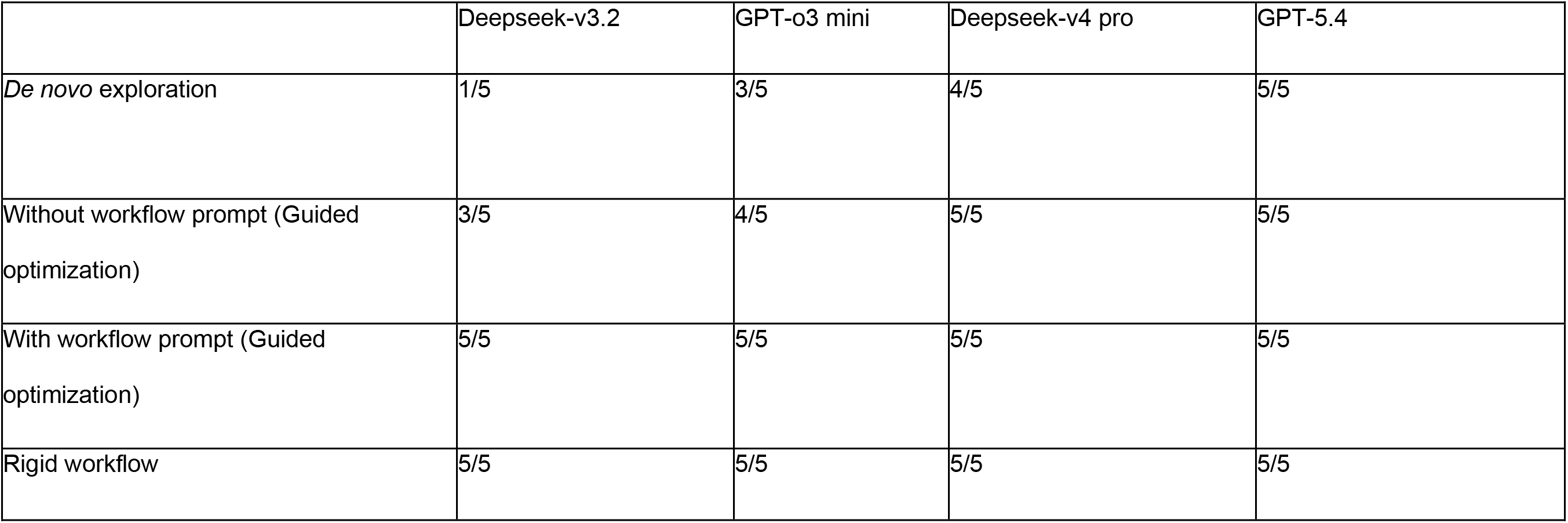
Success rates of different cryoAgent operation modes across differentss language models.

**Table S3.** The available tools and versions in cryoAgent.

| Tools | Version |
| --- | --- |
| cryoSPARC | 4.6 |
| Relion | 5.0.1 |
| Helicon | 1.0 |
| cryoAlign | 2.0 |
| cryoSIFT | NA |
| chimerax | 1.10 |
| CtfFInd | 4.1 |
| MotionCor2 | 1.4 |

**Table S4.**
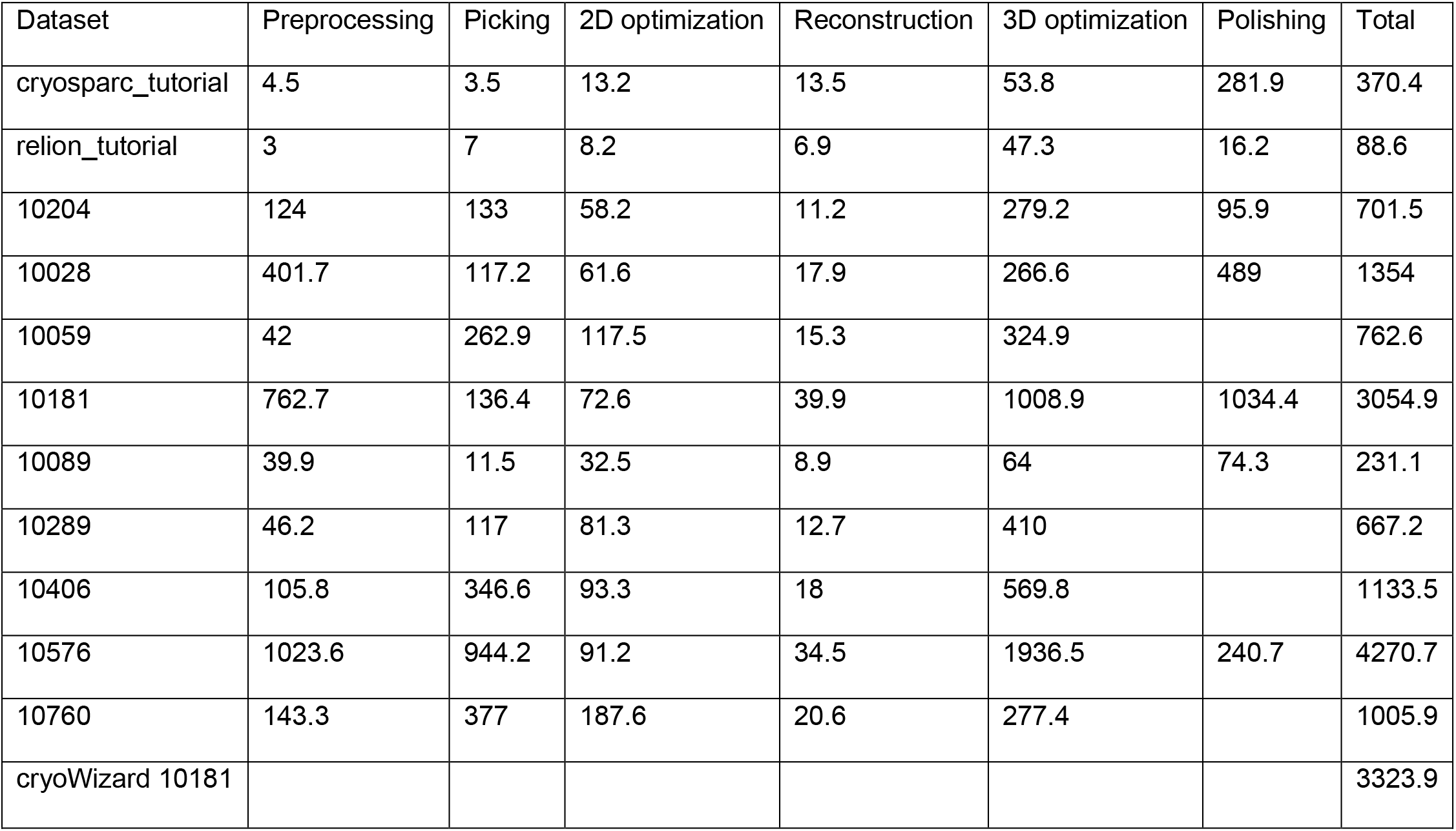
Running times across datasets for cryoAgent using an RTX 5090 GPU with SSD cache in minutes.

| Dataset | Preprocessing | Picking | 2D optimization | Reconstruction | 3D optimization | Polishing | Total |
| --- | --- | --- | --- | --- | --- | --- | --- |
| cryosparc_tutorial | 4.5 | 3.5 | 13.2 | 13.5 | 53.8 | 281.9 | 370.4 |
| relion_tutorial | 3 | 7 | 8.2 | 6.9 | 47.3 | 16.2 | 88.6 |
| 10204 | 124 | 133 | 58.2 | 11.2 | 279.2 | 95.9 | 701.5 |
| 10028 | 401.7 | 117.2 | 61.6 | 17.9 | 266.6 | 489 | 1354 |
| 10059 | 42 | 262.9 | 117.5 | 15.3 | 324.9 |  | 762.6 |
| 10181 | 762.7 | 136.4 | 72.6 | 39.9 | 1008.9 | 1034.4 | 3054.9 |
| 10089 | 39.9 | 11.5 | 32.5 | 8.9 | 64 | 74.3 | 231.1 |
| 10289 | 46.2 | 117 | 81.3 | 12.7 | 410 |  | 667.2 |
| 10406 | 105.8 | 346.6 | 93.3 | 18 | 569.8 |  | 1133.5 |
| 10576 | 1023.6 | 944.2 | 91.2 | 34.5 | 1936.5 | 240.7 | 4270.7 |
| 10760 | 143.3 | 377 | 187.6 | 20.6 | 277.4 |  | 1005.9 |
| cryoWizard 10181 |  |  |  |  |  |  | 3323.9 |

**Table S5.** Running times across datasets for cryoAgent using an RTX 3090 GPU without SSD cache in minutes.

| Dataset | Preprocessing | Picking | 2D optimization | 3D optimization | Reconstruction | Polishing | Total |
| --- | --- | --- | --- | --- | --- | --- | --- |
| 10002 | 64.8 | 45.5 | 317.8 | 194.4 | 22.7 |  | 645.2 |
| 10217 | 182.2 | 282.6 | 251.9 | 2773.6 | 96.6 |  | 3586.9 |
| 10240 | 246.5 | 236.5 | 142.3 | 641.8 | 27.1 |  | 1294.2 |
| 10017 | 16.5 | 18.6 | 82 | 171.9 | 17.5 |  | 306.5 |
| 10081 | 246.5 | 172.8 | 156.9 | 738.6 | 28.3 |  | 1343.1 |
| 10093 | 218 | 271.9 | 376.4 | 1354.4 | 54.5 |  | 2275.2 |
| 10405 | 299 | 151.6 | 197.5 | 5885.2 | 93.1 | 776.1 | 7402.5 |
| 10010 | 196.7 | 73.2 | 264.9 | 3160.6 | 55 |  | 3750.4 |
| 10250 | 387 | 284.1 | 503.6 | 1028.1 | 30.6 | 348.7 | 2582.1 |

## Notes

### Competing Interest Statement

The authors have declared no competing interest.

### Summary of Updates

We have extended the content of the paper, including providing more adaptive ability to the agent and also the designing logic of this agentic workflow.

## References

1. E. Y. D. Chua, et al., Better, Faster, Cheaper: Recent Advances in Cryo–Electron Microscopy. Annu. Rev. Biochem. 91, 1–32 (2022).

2. R. G. Efremov, A. Stroobants, Coma-corrected rapid single-particle cryo-EM data collection on the CRYO ARM 300. Acta Crystallogr. Sect. Struct. Biol. 77, 555–564 (2021).

3. M. Stabrin, et al., TranSPHIRE: automated and feedback-optimized on-the-fly processing for cryo-EM. Nat. Commun. 11, 5716 (2020).

4. D. Tegunov, P. Cramer, Real-time cryo-electron microscopy data preprocessing with Warp. Nat. Methods 16, 1146–1152 (2019).

5. Y. Li, J. N. Cash, J. J. G. Tesmer, M. A. Cianfrocco, High-Throughput Cryo-EM Enabled by User-Free Preprocessing Routines. Structure 28, 858–869.e3 (2020).

6. C. Team, et al., End-to-end automation of repeat-target cryo-EM structure determination in CryoSPARC. [Preprint] (2025). Available at: https://www.biorxiv.org/content/10.1101/2025.10.17.682689v1 [Accessed 10 February 2026].

7. D. Kimanius, L. Dong, G. Sharov, T. Nakane, S. H. W. Scheres, New tools for automated cryo-EM single-particle analysis in RELION-4.0. Biochem. J. 478, 4169–4185 (2021).

8. Y. Yan, S. Fan, F. Yuan, H. Shen, A comprehensive foundation model for cryo-EM image processing. Nat. Methods 23, 88–95 (2026).

9. J.-H. Schaefer, et al., CryoSift - An accessible and automated CNN-driven tool for cryo-EM 2D class selection. [Preprint] (2025). Available at: https://www.biorxiv.org/content/10.1101/2025.07.28.667259v1 [Accessed 4 August 2025].

10. J. Zhou, et al., An AI Agent for Fully Automated Multi-Omic Analyses. Adv. Sci. 11, 2407094 (2024).

11. K. Huang, et al., Biomni: A General-Purpose Biomedical AI Agent. [Preprint] (2025). Available at: https://www.biorxiv.org/content/10.1101/2025.05.30.656746v1 [Accessed 10 February 2026].

12. C. S. Xia, Y. Deng, S. Dunn, L. Zhang, Agentless: Demystifying LLM-based Software Engineering Agents. [Preprint] (2024). Available at: http://arxiv.org/abs/2407.01489 [Accessed 8 July 2026].

13. H. Cao, et al., Mozi: Governed Autonomy for Drug Discovery LLM Agents. [Preprint] (2026). Available at: http://arxiv.org/abs/2603.03655 [Accessed 7 July 2026].

14. S. Yao, et al., ReAct: Synergizing Reasoning and Acting in Language Models. arXiv.org (2022). Available at: https://arxiv.org/abs/2210.03629v3 [Accessed 10 February 2026].

15. CryoSPARC Discuss. *CryoSPARC Discuss*. Available at: https://discuss.cryosparc.com/ [Accessed 8 July 2026].

16. Single particle tutorial — RELION documentation. Available at: https://relion.readthedocs.io/en/release-5.0/SPA_tutorial/ [Accessed 9 July 2026].

17. Z. Liu, et al., CryoAlign2: efficient global and local Cryo-EM map retrieval based on parallel-accelerated local spatial structural features. Bioinformatics 41, btaf296 (2025).

18. E. F. Pettersen, et al., UCSF ChimeraX: Structure visualization for researchers, educators, and developers. Protein Sci. Publ. Protein Soc. 30, 70–82 (2021).

19. W. Wong, et al., Cryo-EM structure of the Plasmodium falciparum 80S ribosome bound to the anti-protozoan drug emetine. eLife 3, e03080 (2014).

20. A. Levy, et al., CryoDRGN-AI: neural ab initio reconstruction of challenging cryo-EM and cryo-ET datasets. Nat. Methods 22, 1486–1494 (2025).

21. E. D. Zhong, T. Bepler, B. Berger, J. H. Davis, CryoDRGN: reconstruction of heterogeneous cryo-EM structures using neural networks. Nat. Methods 18, 176–185 (2021).

22. M. A. Herzik, M. Wu, G. C. Lander, Achieving better-than-3-Å resolution by single-particle cryo-EM at 200 keV. Nat. Methods 14, 1075–1078 (2017).

23. G. Wang, et al., Voyager: An Open-Ended Embodied Agent with Large Language Models. [Preprint] (2023). Available at: http://arxiv.org/abs/2305.16291 [Accessed 8 July 2026].

24. V. Hsiao, M. Roberts, L. Smith, Procedural Knowledge Improves Agentic LLM Workflows. [Preprint] (2025). Available at: http://arxiv.org/abs/2511.07568 [Accessed 13 July 2026].

25. C. Zhang, B. I. Yakobson, MatClaw: An Autonomous Code-First LLM Agent for End-to-End Materials Exploration. [Preprint] (2026). Available at: http://arxiv.org/abs/2604.02688 [Accessed 13 July 2026].

26. A. Punjani, J. L. Rubinstein, D. J. Fleet, M. A. Brubaker, cryoSPARC: algorithms for rapid unsupervised cryo-EM structure determination. Nat. Methods 14, 290–296 (2017).

27. N. Shinn, et al., Reflexion: Language Agents with Verbal Reinforcement Learning. arXiv.org (2023). Available at: https://arxiv.org/abs/2303.11366v4 [Accessed 29 March 2026].

28. C. Packer, et al., MemGPT: Towards LLMs as Operating Systems.

29. S. H. W. Scheres, RELION: Implementation of a Bayesian approach to cryo-EM structure determination. J. Struct. Biol. 180, 519–530 (2012).

30. S. Q. Zheng, et al., MotionCor2: anisotropic correction of beam-induced motion for improved cryo-electron microscopy. Nat. Methods 14, 331–332 (2017).

31. A. Rohou, N. Grigorieff, CTFFIND4: Fast and accurate defocus estimation from electron micrographs. J. Struct. Biol. 192, 216–221 (2015).

32. K. Lasker, O. Dror, M. Shatsky, R. Nussinov, H. J. Wolfson, EMatch: Discovery of High Resolution Structural Homologues of Protein Domains in Intermediate Resolution Cryo-EM Maps. IEEE/ACM Trans. Comput. Biol. Bioinform. 4, 28–39 (2007).

33. Z. Zhou, A. A. Amini, Analysis of spectral clustering algorithms for community detection: the general bipartite setting. arXiv.org (2018). Available at: https://arxiv.org/abs/1803.04547v2 [Accessed 29 March 2026].

34. P. B. Rosenthal, R. Henderson, Optimal Determination of Particle Orientation, Absolute Hand, and Contrast Loss in Single-particle Electron Cryomicroscopy. J. Mol. Biol. 333, 721– 745 (2003).

35. T. Moriya, et al., Size matters: optimal mask diameter and box size for single-particle cryogenic electron microscopy. [Preprint] (2020). Available at: https://www.biorxiv.org/content/10.1101/2020.08.23.263707v1 [Accessed 29 March 2026].

36. F. J. Sigworth, Principles of cryo-EM single-particle image processing. Microscopy 65, 57–67 (2016).

37. EMPIAR-10217 bovine liver glutamate dehydrogenase. Available at: https://www.ebi.ac.uk/empiar/EMPIAR-10217/ [Accessed 17 March 2026].

38. Get Started with CryoSPARC: Introductory Tutorial (v4.0+) | CryoSPARC Guide. (2026). Available at: https://guide.cryosparc.com/processing-data/get-started-with-cryosparc-introductory-tutorial [Accessed 29 March 2026].

39. Single particle tutorial — RELION documentation. Available at: https://relion.readthedocs.io/en/latest/SPA_tutorial/ [Accessed 29 March 2026].

40. Y. Gao, E. Cao, D. Julius, Y. Cheng, TRPV1 structures in nanodiscs reveal mechanisms of ligand and lipid action. Nature 534, 347–351 (2016).

41. EMPIAR-10204 The first reconstruction of beta-galactosidase solved by cryoARM200. Available at: https://www.ebi.ac.uk/empiar/EMPIAR-10204/ [Accessed 29 March 2026].

42. T. Moriya, et al., High-resolution Single Particle Analysis from Electron Cryo-microscopy Images Using SPHIRE. J. Vis. Exp. JoVE 55448 (2017). 10.3791/55448.

43. X. Bai, I. S. Fernandez, G. McMullan, S. H. Scheres, Ribosome structures to near-atomic resolution from thirty thousand cryo-EM particles. eLife 2, e00461 (2013).

44. M. E. Falzone, et al., Structural basis of Ca2+-dependent activation and lipid transport by a TMEM16 scramblase. eLife 8, e43229 (2019).

45. S. H. W. Scheres, Semi-automated selection of cryo-EM particles in RELION-1.3. J. Struct. Biol. 189, 114–122 (2015).

46. C.-H. Lee, R. MacKinnon, Structures of the Human HCN1 Hyperpolarization-Activated Channel. Cell 168, 111–120.e11 (2017).

47. P. Jin, et al., Electron cryo-microscopy structure of the mechanotransduction channel NOMPC. Nature 547, 118–122 (2017).

48. X. Huang, et al., Amorphous nickel titanium alloy film: A new choice for cryo electron microscopy sample preparation. Prog. Biophys. Mol. Biol. 156, 3–13 (2020).

49. Z. Wang, et al., An atomic model of brome mosaic virus using direct electron detection and real-space optimization. Nat. Commun. 5, 4808 (2014).

50. M. A. Herzik, M. Wu, G. C. Lander, High-resolution structure determination of sub-100 kDa complexes using conventional cryo-EM. Nat. Commun. 10, 1032 (2019).

51. B. Burendei, et al., Cryo-EM structures of undocked innexin-6 hemichannels in phospholipids. Sci. Adv. 6, eaax3157 (2020).

52. D. Nicholson, T. A. Edwards, A. J. O’Neill, N. A. Ranson, Structure of the 70S Ribosome from the Human Pathogen Acinetobacter baumannii in Complex with Clinically Relevant Antibiotics. Structure 28, 1087–1100.e3 (2020).

53. B.-R. Zhou, et al., Distinct Structures and Dynamics of Chromatosomes with Different Human Linker Histone Isoforms. Mol. Cell 81, 166–182.e6 (2021).

54. M. Kuzuya, et al., Structures of human pannexin-1 in nanodiscs reveal gating mediated by dynamic movement of the N terminus and phospholipids. Sci. Signal. 15, eabg6941 (2022).

55. Y. Z. Tan, et al., Addressing preferred specimen orientation in single-particle cryo-EM through tilting. Nat. Methods 14, 793–796 (2017).

56. A. Kucukelbir, F. J. Sigworth, H. D. Tagare, Quantifying the local resolution of cryo-EM density maps. Nat. Methods 11, 63–65 (2014).

